# Live Imaging of SARS-CoV-2 Infection in Mice Reveals Neutralizing Antibodies Require Fc Function for Optimal Efficacy

**DOI:** 10.1101/2021.03.22.436337

**Authors:** Irfan Ullah, Jérémie Prévost, Mark S Ladinsky, Helen Stone, Maolin Lu, Sai Priya Anand, Guillaume Beaudoin-Bussières, Kelly Symmes, Mehdi Benlarbi, Shilei Ding, Romain Gasser, Corby Fink, Yaozong Chen, Alexandra Tauzin, Guillaume Goyette, Catherine Bourassa, Halima Medjahed, Matthias Mack, Kunho Chung, Craig B Wilen, Gregory A. Dekaban, Jimmy D. Dikeakos, Emily A. Bruce, Daniel E Kaufmann, Leonidas Stamatatos, Andrew T. McGuire, Jonathan Richard, Marzena Pazgier, Pamela J. Bjorkman, Walther Mothes, Andrés Finzi, Priti Kumar, Pradeep D. Uchil

**Affiliations:** Department of Internal Medicine, Section of Infectious Diseases, Yale University School of Medicine, New Haven, CT 06520, USA; Centre de Recherche du CHUM, Montreal, QC, H2X0A9, Canada; Département de Microbiologie, Infectiologie et Immunologie, Université de Montréal, Montreal, QC, H2X0A9, Canada; Division of Biology and Biological Engineering, California Institute of Technology, Pasadena, CA 91125, USA; Department of Microbial Pathogenesis, Yale University School of Medicine, New Haven CT 06510, USA; Department of Microbiology and Immunology, McGill University, Montreal, Qc, H3A 2B4, Canada; Department of Microbiology and Immunology, University of Western Ontario, London, ON, N6A 5B7, Canada; Infectious Disease Division, Department of Medicine, Uniformed Services University of the Health Sciences, Bethesda, MD 20814, USA; Universitätsklinikum Regensburg, Innere Medizin II – Nephrologie, 93042 Regensburg, Germany; Departments of Laboratory Medicine and Immunobiology, Yale University School of Medicine, New Haven, CT 06520, USA; Division of Immunobiology, Dept. of Medicine, Larner College of Medicine, University of Vermont. Burlington, VT 05405. USA.; Vaccine and Infectious Disease Division, Fred Hutchinson Center, Seattle, WA 98195, USA; Molecluar Medicine Research Laboratories, Robarts Research Institute, University of Western Ontario, London, ON, N6A 5B7, Canada; Department of Global Health, University of Washington, Seattle, WA 98195, USA; Department of Laboratory Medicine and Pathology, University of Washington, Seattle, WA

**Keywords:** SARS-CoV-2, COVID-19, nanoluciferase, bioluminescence imaging, neutralizing antibodies, convalescent patients, human ACE2 transgenic mice, monocytes, natural killer cells, pathogenesis, inflammatory cytokines, Fc effector functions

## Abstract

Neutralizing antibodies (NAbs) are effective in treating COVID-19 but the mechanism of immune protection is not fully understood. Here, we applied live bioluminescence imaging (BLI) to monitor the real-time effects of NAb treatment in prophylaxis and therapy of K18-hACE2 mice intranasally infected with SARS-CoV-2-nanoluciferase. We could visualize virus spread sequentially from the nasal cavity to the lungs and thereafter systemically to various organs including the brain, which culminated in death. Highly potent NAbs from a COVID-19 convalescent subject prevented, and also effectively resolved, established infection when administered within three days. In addition to direct Fab-mediated neutralization, Fc effector interactions of NAbs with monocytes, neutrophils and natural killer cells were required to effectively dampen inflammatory responses and limit immunopathology. Our study highlights that both Fab and Fc effector functions of NAbs are essential for optimal *in vivo* efficacy against SARS-CoV-2.

## Introduction

SARS-CoV-2-neutralizing monoclonal antibodies (NAbs) are an attractive countermeasure for both COVID-19 prevention and therapy (Schafer et al., 2021; Voss et al., 2020; Weinreich et al., 2021). To date, multiple NAbs against the spike (S) glycoprotein of SARS-CoV-2 have been isolated from convalescent subjects. The majority of NAbs bind to the receptor binding domain (RBD) in the S1 subunit and inhibit virus attachment to the human Angiotensin Converting Enzyme 2 (hACE2) receptor. NAbs against the N-terminal domain (NTD) of S1 as well as the S2 subunit have also been isolated (Liu et al., 2020; Voss et al., 2020). NAbs have demonstrated varying levels of efficacy and protection in multiple animal models of SARS-CoV-2 (Alsoussi et al., 2020; Baum et al., 2020; Fagre et al., 2020; Hansen et al., 2020; Hassan et al., 2020; Li et al., 2020; Rogers et al., 2020; Shi et al., 2020b; Winkler et al., 2020; Zost et al., 2020a; Zost et al., 2020b). However, the *in vitro* neutralization potency of NAbs has not consistently correlated with *in vivo* protection (Bournazos et al., 2014; Schafer et al., 2021). While the antigen binding domain (Fab) of antibodies are critical for neutralization, the fragment crystallizable (Fc) domain can contribute significantly to their *in vivo* efficacy (Bournazos et al., 2019; Bournazos et al., 2014; DiLillo et al., 2014; Lu et al., 2018). Fc effector functions can also be detrimental to the host, especially against respiratory viruses such as respiratory syncytial virus (RSV) and SARS-CoV-1 leading to antibody-dependent enhancement (ADE) and aggravated disease pathology (Bolles et al., 2011; Halstead and Katzelnick, 2020; Ruckwardt et al., 2019). Therefore, a careful investigation of NAb mechanisms that elicit protective or pathological consequences is required prior to clinical deployment.

Animal models evaluated to date (Johansen et al., 2020; Leist et al., 2020a; Leist et al., 2020b) have not fully recapitulated pathological features of human COVID-19. Transgenic mice expressing hACE2 under the cytokeratin 18 promoter (K18-hACE2 mice) have some distinct advantages. Their heightened susceptibility to human-tropic SARS-CoV-2 virus strains with mortality ensuing within a week (McCray et al., 2007; Shi et al., 2020a; Winkler et al., 2020) sets a high bar for identifying effective intervention strategies. In humans, SARS-CoV-2 infection disables innate immunity and elicits an imbalanced inflammatory cytokine response in the lungs leading to acute respiratory distress syndrome (ARDS) which is the major cause of death (Graham and Baric, 2020). K18-hACE2 mice also display lung inflammation, cytokine storm and impaired respiratory function (Winkler et al., 2020). However, mortality is due to neuroinvasion (Carossino et al., 2021; Golden et al., 2020; Leist et al., 2020a). Interestingly, many SARS-CoV-2 patients display a myriad of neurological symptoms (Ellul et al., 2020). Finally, mouse FcγRs display similar affinities to human antibodies (Dekkers et al., 2017). Therefore, K18-hACE2 mice are excellent models for evaluating candidate human anti-SARS-CoV-2 Abs and anti-viral interventions.

Bioluminescence imaging (BLI)-guided studies permit live visualization of pathogen spread, in relevant tissues enabling real-time outcome assessments for treatment regimens. A BLI-driven platform has not yet been harnessed for studying infectious respiratory pathogens like SARS-CoV-2 that require level 3 biosafety containment. Here, we established a BLI-driven approach to study SARS-CoV-2 infection with a well characterized replication competent SARS-CoV-2 virus carrying a nanoluciferase (nLuc) reporter in the place of the ORF7A gene (Xie et al., 2020a; Xie et al., 2020b). SARS-CoV-2-nLuc closely mimics the wildtype virus replication kinetics and stably maintains the nLuc reporter over five generations *in vitro*. Further, ORF7a deletion was recently shown not to affect pathology of the wildtype virus (Silvas et al., 2021). *In vivo* BLI revealed that the virus spreads from the nasal cavity to lungs to establish infection. This is followed by sequential infection of cervical lymph nodes (cLNs), brain, and finally systemic dissemination. Upon neuroinvasion, the virus replicates rapidly in the brain leading to fulminant infection and death by 6-7 days post infection (dpi). A single prophylactic intraperitoneal (i.p.) administration of highly potent NAbs isolated from a convalescent COVID-19 subject prevented SARS-CoV-2-induced mortality in K18-hACE2 mice. Protection was associated with widespread localization of administered NAbs and Fc-mediated effector functions with contributions from neutrophils, monocytes and NK cells as well as reduced induction of inflammatory cytokines. BLI also revealed a therapeutic window of 3 dpi for NAb for successfully halting progression and spread of infection from the lungs. Thus, our BLI-driven study highlights that both neutralizing and Fc effector functions of NAbs are essential for optimal *in vivo* efficacy against SARS-CoV-2.

## Results

### BLI allows Visualization of SARS-CoV-2 Replication Dynamics and Pathogenesis

We tracked spread of SARS-CoV-2-nLuc using BLI after intranasal (i.n.) challenge in K18-hACE2 mice **(Figure 1A)**. 1 x 10^5^ FFU of SARS-CoV-2 generated sufficient photon flux to allow non-invasive BLI with luciferase signal detected only in C57BL/6J (B6) mice expressing hACE2 **(Figure 1B)**. Temporal tracking of emitted light intensities revealed that the virus replicated in the nasal cavity in a biphasic manner **(Figure 1C)**. Luminescent signal in the nose increased the first two days of infection and diminished before increasing again between 5 to 6 dpi when systemic spread occurred. The first signs of infection in the lungs were observed at 1 dpi. The nLuc signal then steadily increased in until 3 dpi. nLuc signals in the cLNs and brain region were detected (imaging in ventral position) at 4 dpi. A steep rise in brain nLuc activity occurred from 4 to 6 dpi indicating neuroinvasion and robust virus replication **(Figure 1B, C, Video S1)**. This was accompanied by widespread virus replication in the gut and genital tract with loss in body weight. By 6 dpi, infected K18-hACE2 mice lost 20% of their initial body weight, became moribund and succumbed **(Figure 1D, E)**. In contrast, as expected, B6 mice did not experience any weight loss and survived the virus challenge.

**Figure 1.**
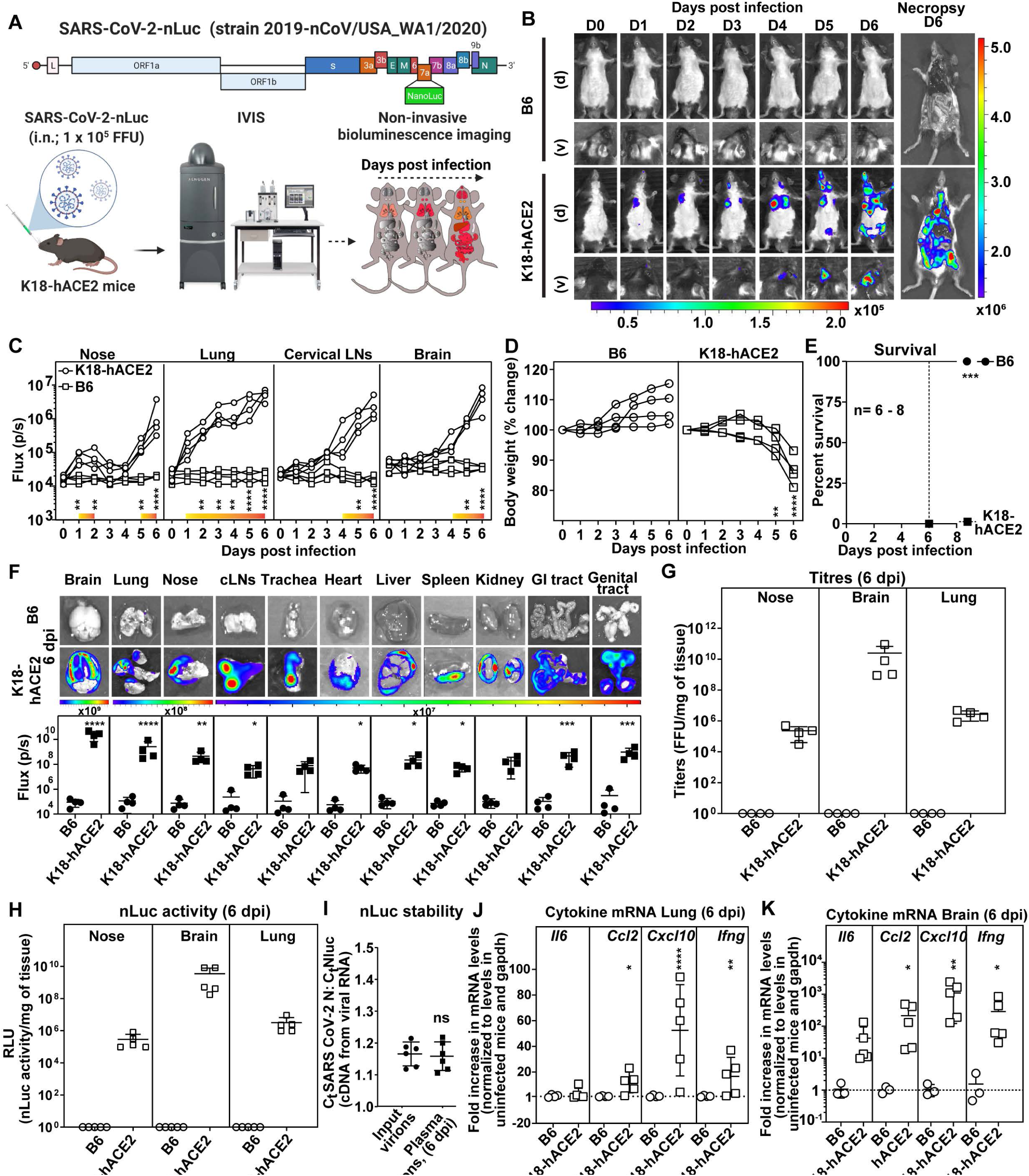
Visualization of SARS-CoV-2 Replication Dynamics in hACE2 Transgenic Mice. (A) Experimental strategy utilizing SARS-CoV-2 carrying nLuc reporter in ORF7a for non-invasive BLI of virus spread following intranasal (i.n.) challenge of B6 or K18-hACE2 mice. (B) Representative images from temporal BLI of SARS-CoV-2-nLuc-infected mice in ventral (v) and dorsal (d) positions at the indicated dpi and after necropsy (C) Temporal quantification of nLuc signal as flux (photons/sec) acquired non-invasively in the indicated tissues of each animal. The color bar above the x-axis (yellow to orange) represents computed signal intensities in K18-hACE2 mice that are significantly above those in B6 mice. (D) Temporal changes in mouse body weight with initial body weight set to 100%. (E) Kaplan-Meier survival curves of mice for experiment as in A statistically compared by log-rank (Mantel-Cox) test. (F) *Ex vivo* imaging of indicated organs and quantification of nLuc signal as flux(photons/sec) at 6 dpi after necropsy. (G, H) Viral loads (FFUs/mg or nLuc activity/mg) in indicated tissue measured on Vero E6 cells as targets. Non-detectable virus amounts were set to 1. (I) Ratio of C_t_ values for SARS-CoV-2 nucleocapsid (N) and nLuc estimated by RT-PCR using RNA extracted from input virions (inoculum) and virions from sera of mice at 6 dpi. (J, K) Fold changes in cytokine mRNA levels in lung and brain tissues at 6 dpi after normalization to *GAPDH* mRNA in the same sample and that in uninfected mice. Each curve in (C) and (D) and each data point in (F), (I), (J), and (K) represents an individual mouse. Scale bars in (B and (F) denote radiance (photons/sec/cm^2^/steradian). *p* values obtained by non-parametric Mann-Whitney test for pairwise comparison. ∗, p < 0.05; ∗∗, p < 0.01; ∗∗∗, p < 0.001; ∗∗∗∗, p < 0.0001; ns, not significant; Mean values ± SD are depicted.

To visualize the extent of viral spread with enhanced sensitivity and resolution, we imaged individual organs after necropsy **(Figure 1B, F)**. Most organs analyzed from K18-hACE2 mice showed nLuc activity with maximum signal in the brain followed by the lung and nasal cavity **(Figure 1F)** and mirrored the viral loads [Focus Forming Units (FFUs) and nLuc activity] measured in these tissues **(Figure 1G, H)**. Real-time PCR and histology to detect hACE2 and SARS-CoV-2 N confirmed widespread infection in keeping with hACE2 expression in individual tissues/organs **(Table S1, Figure S1A)**.

Reporter-expressing viruses often purge foreign genes, particularly *in vivo*, due to fitness and immune pressure (Falzarano et al., 2014; Ventura et al., 2019). The ratio of copy numbers of SARS-CoV-2 nucleocapsid (N) to nLuc in the viral RNA by real-time PCR analyses of input virions and virions isolated from sera of mice at 6 dpi, however, remained unchanged **(Figure 1I)** indicating that the reporter was stable throughout the experimental timeline. Thus, nLuc activity was a good surrogate for virus replication *in vivo*.

SARS-CoV-2 infection triggers an imbalanced immune response and a cytokine storm that contributes substantially to pathogenesis (Del Valle et al., 2020). mRNA levels of inflammatory cytokines *IL6, CCL2, CXCL10* and *IFNγ* in the lungs and brains of mice after necropsy at 6 dpi were significantly upregulated in infected K18-hACE2 mice compared to B6 **(Figure 1J, K)**. Consistent with enhanced inflammation, we observed infiltration of Ly6G^+^ neutrophils and Ly6C^+^ monocytes in both lung and brain (Figure S1C). Overall, cytokine mRNAs were highest in the brain with *CXCL10* mRNA copy numbers ∼1000 fold higher in K18-hACE2 than in B6 mice corroborating extensive infection **(Figure 1J, K)**.

We used BLI data to pinpoint infected regions within lungs, brain, and testis for directed histology and electron tomographic studies **(Figure 2)**. Higher resolution imaging revealed that SARS-CoV-2 viruses were associated to large extent with capillary endothelial cell and/or alveolar type-1 cells in close vicinity to alveolar macrophages in the lungs **(Figure 2A-D; Video S2, S3)**. In the brain, neuronal cells (hACE2^+^MAP2^+^GFAP^-^CD68^-^CD11b^-^) were positive for SARS-CoV-2 N and EM tomography revealed an array of SARS-CoV-2 viruses associated within the dendrites **(Figure 2E-J, Figure S1B, Video S4)**. In the testis, Sertoli cells stained positively for N **(Figure 2K-N, Video S5)**. EM tomography also showed a large population of virions within pleomorphic membrane-bound compartments of Sertoli cells.

**Figure 2.**
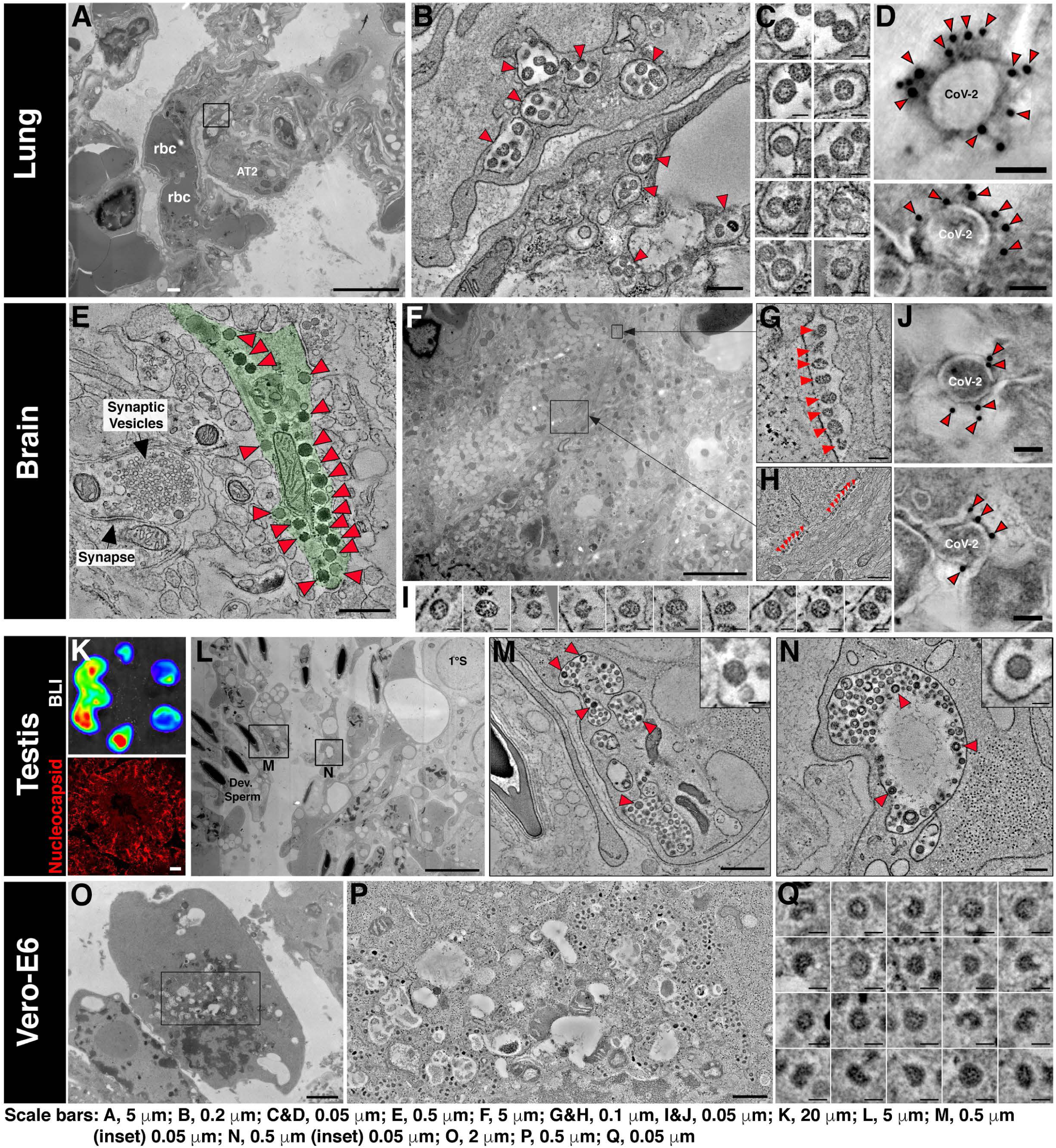
EM localization of SARS-CoV-2 Virions in Lung, Brain and Testis of Infected K18-hACE2 Mice. (A) 2D overview of a lung region featuring red blood cells (rbc) within a pulmonary capillary, an alveolar Type 2 cell (AT2). (B) Slice from a 3D tomogram of square region in A showing membrane-enclosed cytoplasmic compartments (arrowheads) containing presumptive SARS-CoV-2 virions in capillary endothelial cells. (C) Presumptive virions from tomogram in B displayed at equatorial views. Presumptive virions were identified as described in *Methods* and are directly comparable to those in SARS-CoV-2 infected Vero-E6 cells (panels O-Q). (D) ImmunoEM tomography of presumptive SARS-CoV-2 virions from infected lung tissue, labeled with antiserum against Spike protein and gold (10 nm) conjugated secondary antibodies. Gold particles localized to the outer peripheries of virions indicate specific labeling of SARS-CoV-2 Spikes. (E) Tomography of SARS-CoV-2 infected brain tissue. Presumptive SARS-CoV-2 virions (red arrowheads) are present within a neuron (pale green). A dendritic synaptic terminal to the left of the virus-containing neuron shows that presumptive SARS-CoV-2 virions are easily distinguished from typical synaptic neurotransmitter vesicles. (F) 2D overview of brain tissue illustrating the complex spatial relationship among neurons and other brain cell types. Presumptive SARS-CoV-2 virions are present in two compartments (black squares) within a single neuron. (G, H) Tomographic slices of black squares in F. Presumptive SARS-CoV-2 virions (red arrowheads) appear to be aligned within compartments that border the edges of a neural projection. (I) Presumptive SARS-CoV-2 virions from tomograms in G and H. (J) ImmunoEM tomography as in D of presumptive SARS-CoV-2 virions from infected brain tissue. (K) (Upper) BLI of testis from a SARS-CoV-2 infected mouse to identify infected regions for IF and EM analyses. (Lower) IF image of an infected testis region stained with antibodies to SARS-CoV-2 Nucleocapsid (red) (L) 2D overview of testis corresponding to region of high intensity (red) in the upper panel of K, showing Sertoli cells surrounded by developing sperm (left) and one primary spermatocyte (1°S, upper right). Presumptive SARS-CoV-2 virions are localized to membrane-bound compartments in Sertoli cells (black squares). (M, N) Slices from two 3D tomograms of squares in L. Presumptive SARS-CoV-2 virions (arrowheads) are present within membrane-enclosed cytoplasmic compartments. These compartments contain additional structures amongst the discernable SARS-CoV-2 virions (insets). (O) EM localization of virions in SARS-CoV-2 infected Vero-E6 cells, processed for EM as above tissue samples. Virions were characterized (see Methods) and compared to presumptive virions in the tissue samples to confidently verify their identities. 2D overview of infected Vero-E6 cell in a 150 nm section. (P) Tomogram of rectangle in O showing >100 presumptive SARS-CoV-2 virions contained within cytoplasmic exit compartments. (Q) Virions from the tomogram in P showing common features of dense RNC puncta, discernable surface spikes, vary in size (∼60-120 nm) and shape. Virions are directly comparable to those shown for the tissue samples in C and I. Scale bar length for each image is shown at the bottom.

### Highly Potent SARS-CoV-2 NAbs CV3-1 and CV3-25 from a Convalescent Donor

We recently characterized plasma from a COVID-19 convalescent subject (S006) with potent neutralizing activity and high levels of SARS-CoV-1 cross-reactive Abs (Lu et al., 2020). We probed the B cell receptor (BCR) repertoire from this donor to isolate broad and potent NAbs. Using recombinant SARS-CoV-2 S ectodomain (S2P) as bait for antigen-specific B cells, we screened a library of S-targeted BCR clones to identify two potent NAb candidates: CV3-1 and CV3-25 (Jennewein et al., 2021). We characterized their epitope specificity using ELISA, cell-surface staining, virus capture assay and surface plasmon resonance (SPR) (Ding et al., 2020; Prevost et al., 2020). Both NAbs recognized SARS-CoV-2 S, as a stabilized ectodomain (S-6P) or when displayed on cells and virions, with low-nanomolar affinity **(Figure 3A-E)**. While CV3-1 bound the SARS-CoV-2 RBD, CV3-25 targeted the S2 subunit **(Figure 3A, D-E)** and cross-reacted with SARS-CoV-1 S on cells or virions, but not with S from other human coronaviruses **(Figure 3B-C).** In agreement with the previous smFRET data for S006 plasma, CV3-1 stabilized S in the RBD-up (∼0.1 FRET) conformation **(Figure 3F-H)**, as seen with hACE2 and most RBD-directed NAbs (Lu et al., 2020). Interestingly, CV3-25-bound S showed a partial shift towards downstream conformations (∼0.1 and ∼0.3 FRET), suggesting a distinct inhibitory mechanism from CV3-1 **(Figure 3F-H)**.

**Figure 3.**
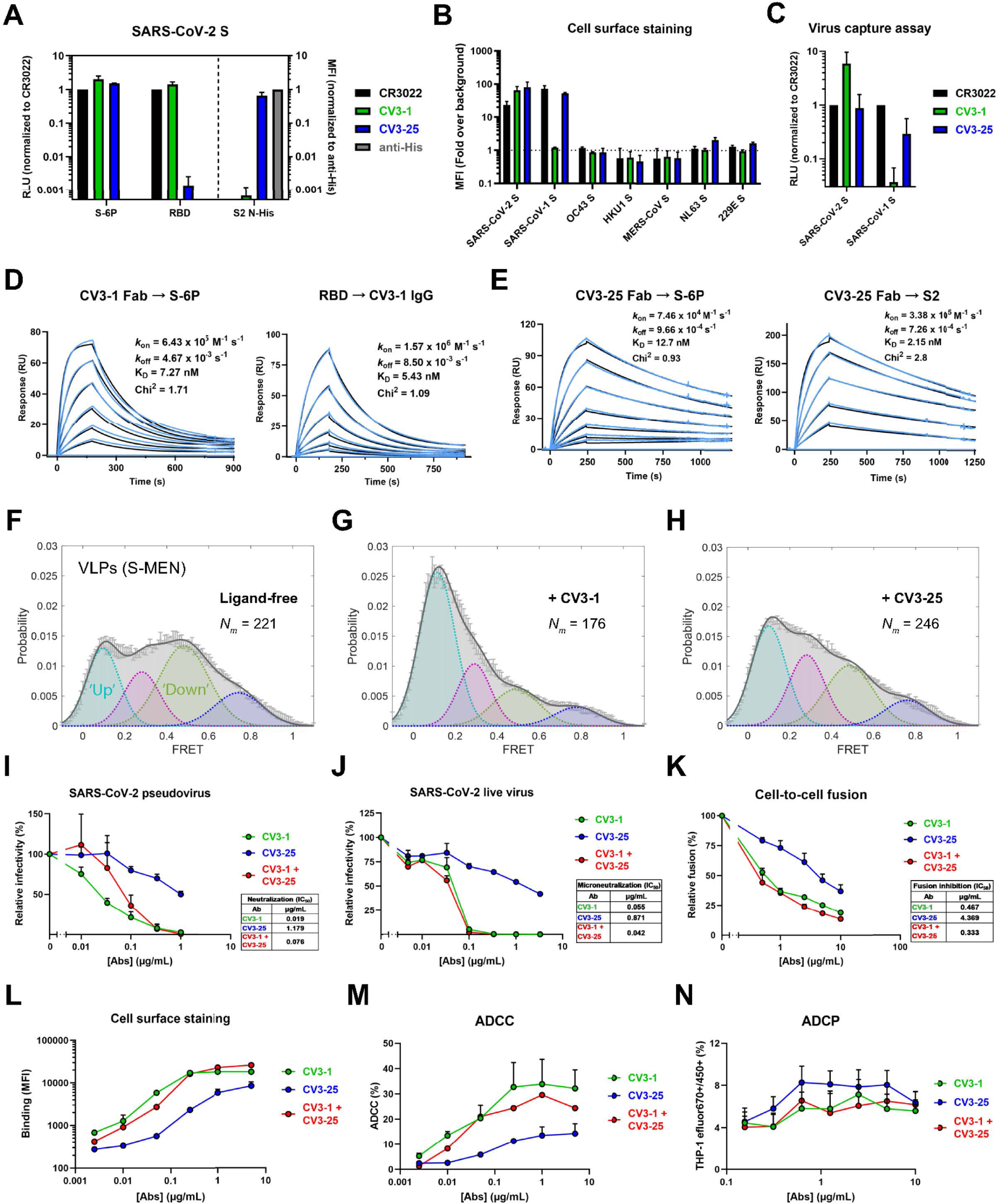
*In vitro* Characterization of CV3-1 and CV3-25 Nabs. (A) NAb binding to SARS-CoV-2 Spike ectodomain (S-6P) or RBD estimated by ELISA. Relative light units (RLU) were normalized to the cross-reactive SARS-CoV-1 mAb CR3022. NAb binding to SARS-CoV-2 S2 N-His tag protein on cell-surface of transfected 293T cells analyzed by flow cytometry. Median fluorescence intensities (MFIs) for anti-Spike NAbs were normalized to the signal obtained with an anti-His tag mAb. (B) Flow cytometric detection of 293T cells expressing S from the indicated human CoVs. MFI from 293T cells transfected with empty vector was used for normalization. (C) Pseudoviruses bearing SARS-CoV-2 or SARS-CoV-1 S were tested for capture by anti-Spike NAbs. The cross-reactive CR3022 mAb was used for normalization. (D-E) NAb binding affinity and kinetics to SARS-CoV-2 S using Surface Plasmon Resonance (SPR). SARS-CoV-2 S-6P or S2 ectodomain was immobilized as the ligand on the chip and CV3-1 or CV3-25 Fab was used as analyte at concentrations ranging from 1.56 to 100 nM for both Fabs to S-6P and 3.125nM to 200nM for CV3-25 to S2 (2-fold serial dilution, see Methods for details). Alternatively, CV3-1 IgG was immobilized on the chip and SARS-CoV-2 RBD used as analyte from 1.56 to 50 nM (2-fold serial dilution). Kinetic constants were determined using a 1:1 Langmuir model in BIA evaluation software (experimental readings depicted in blue and fitted curves in black). (F-H) FRET histograms of ligand-free S on S-MEN coronavirus-like particles (VLPs) or in presence of 50 µg/mL of CV3-1 (G) or CV3-25 (H). VLPs were incubated for 1 h at 37°C before smFRET imaging. *N_m_* is the number of individual FRET traces compiled into a conformation-population FRET histogram (gray lines) and fitted into a 4-state Gaussian distribution (solid black) centered at 0.1-FRET (dashed cyan), 0.3-FRET (dashed red), 0.5-FRET (dashed green), and 0.8- FRET (dashed magenta). (I) Neutralizing activity of CV3-1 and CV3-25 alone or in combination (1:1 ratio) on SARS-CoV-2 S bearing pseudoviruses using 293T-ACE2 cells. (J) Microneutralization activity of anti-Spike NAbs on live SARS-CoV-2 virus using Vero E6 cells. (K) Inhibition of cell-to-cell fusion between 293T cells expressing HIV-1 Tat and SARS-CoV-2 S and TZM-bl-ACE2 cells by NAbs. Half maximal inhibitory antibody concentration (IC_50_) values in I-K were determined by normalized non-linear regression analyses. (L) MFI of CEM.NKr cells expressing SARS-CoV-2 Spike (CEM.NKr-Spike) stained with indicated amounts of NAbs and normalized to parental CEM.NKr. (M) % ADCC in the presence of titrated amounts of NAbs using 1:1 ratio of parental CEM.NKr cells and CEM.NKr-Spike cells as targets when PBMCs from non-infected donors were used as effector cells (N) % ADCP in the presence of titrated amounts of NAbs using CEM.NKr-Spike cells as targets and THP-1 cells as phagocytic cells.

We next measured neutralization and Fc-dependent functions of CV3-1 and CV3-25. While both NAbs blocked infection by SARS-CoV-2 pseudovirus or live virus and interfered with S-driven cell-to-cell fusion, CV3-1 was ∼10 times more potent than CV3-25 **(Figure 3I-L)**. To evaluate Fc-mediated effector functions, we used assays that quantify the antibody-dependent cellular cytotoxicity (ADCC) and antibody-dependent cellular phagocytosis (ADCP) activities. CV3-1 and CV3-25 efficiently bound and eliminated S-expressing cells by stimulating cytotoxic and phagocytic responses in immune effector cells **(Figure 3L-N)**. Overall, both NAbs displayed significant neutralization and Fc-dependent antibody functions, although CV3-1 was found to be more effective. The combinatorial effect of the two NAbs (1:1 ratio) was similar to the response with CV3-1 alone **(Figure 3I-N).**

### Prophylactic Treatment with NAbs Protects K18-hACE2 Mice from SARS-CoV-2 Infection

We first monitored the tissue biodistribution of Alexa Fluor 647 (AF_647_) conjugated CV3-1 and AF_594_-conjugated CV3-25 in various tissues 24 h after i.p. delivery in mice by fluorescence imaging, histology and ELISA. All three approaches revealed widespread distribution of both NAbs in multiple organs including the SARS-CoV-2 target tissues nasal cavity, lung and the brain **(Figure S2, S3A-D)**. We next tested a prophylactic regimen where each NAb was delivered i.p. alone (12.5 mg/kg body weight) or in 1:1 combination (6.25 mg each NAb/kg body weight) 24 h before i.n. challenge with SARS-CoV-2 nLuc **(Figure 4A)**. Temporal monitoring by whole-body BLI revealed that all three prophylactic regimens substantially reduced SARS-CoV-2 infection in the lungs and subsequent spread **(Figure 4B-D)**. Remarkably, pretreatment with CV3-1 alone or in combination with CV3-25 (cocktail 1:1) produced near complete protection from SARS-CoV-2 infection with no signals detected in most organs by live non-invasive imaging or after terminal necropsy at 22 dpi **(Figure 4B-D, G, H)**. Moreover, all test cohorts survived with no discernible weight loss, nLuc activity or viral loads signifying complete control of virus infection **(Figure 4E-I)**. In CV3-25-pretreated animals, lung infection and subsequent neuroinvasion occurred at reduced intensity, and was also reflected in individual organs after necropsy **(Figure 4B, G, H)**. CV3-25 delayed mortality by ∼2 days in 4 out of the 6 animals and viral loads in the nasal cavity, lungs, and brain at the time of necropsy (8 dpi) were similar to that in the control cohorts treated with isotype-matched antibodies at 6 dpi **(Figure 4F, I)**.

**Figure 4.**
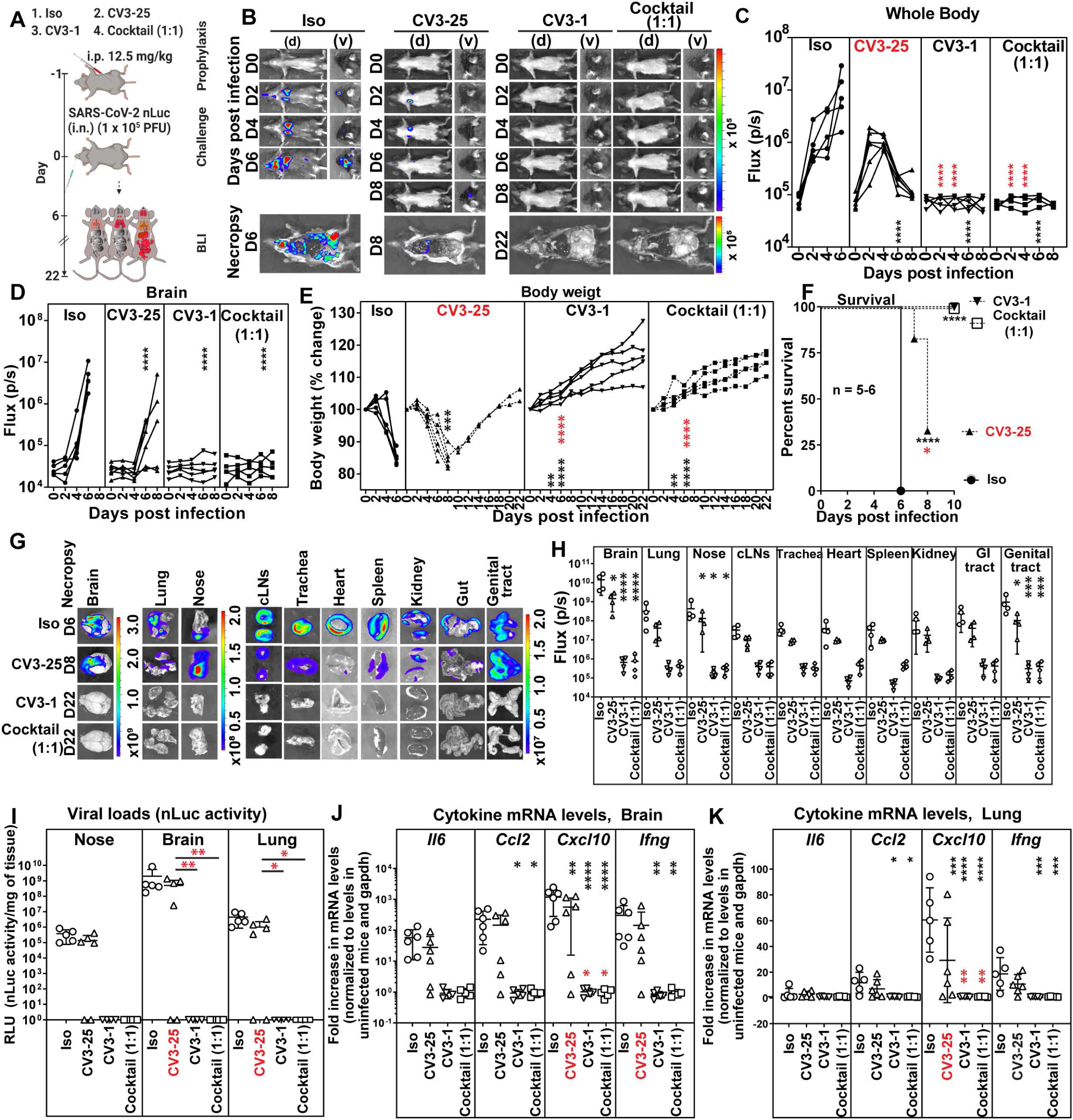
Prophylactic Treatment with CV3-1 Protects Mice from Lethal SARS-CoV-2 Infection. (A) Experimental design to test *in vivo* efficacy of NAbs CV3-1 and CV3-25 administered alone (12.5 mg/kg body weight) or as a 1:1 cocktail (6.25 mg/kg body weight each) 1 day prior to challenging K18-hACE2 mice (i.n.) with SARS-CoV-2-nLuc followed by non-invasive BLI every 2 days. Human IgG1-treated (12.5 mg/kg body weight) mice were the control cohort (Iso). (B) Representative BLI images of SARS-CoV-2-nLuc-infected mice in ventral (v) and dorsal (d) positions. (C-D) Temporal quantification of nLuc signal as flux (photons/sec) computed non-invasively. (E) Temporal changes in mouse body weight with initial body weight set to 100%. (F) Kaplan-Meier survival curves of mice statistically compared by log-rank (Mantel-Cox) test. (G, H) *Ex-vivo* images of organs and nLuc signal quantified as flux (photons/sec) after necropsy. (I) Viral loads (nLuc activity/mg tissue) measured in Vero E6 cells as targets. Non-detectable virus amounts were set to 1. (J, K) Fold changes in cytokine mRNA levels in lung and brain tissues normalized to *GAPDH* mRNA in the same sample and that in non-infected mice after necropsy. Viral loads (I) and inflammatory cytokine profile (J, K) were determined after necropsy for mice that succumbed to infection at 6dpi and in mice surviving at 22 dpi. Scale bars in (B) and (G) denote radiance (photons/sec/cm^2^/steradian). Each curve in (C)-(E) and each data point in (H)-(K) represents an individual mouse. Grouped data in (C)-(K) were analyzed by 2-way ANOVA followed by Dunnett’s or Tukey’s multiple comparison tests. Statistical significance for group comparisons to isotype control are shown in black and for those to CV3-25 are shown in red. ∗, p < 0.05; ∗∗, p < 0.01; ∗∗∗, p < 0.001; ∗∗∗∗, p < 0.0001; Mean values ± SD are depicted.

Pre-treatment with CV3-1 or NAb cocktail also prevented the inflammatory cytokine induction **(Figure 4J, K)**. In contrast, heightened levels of cytokine mRNA were detected in mice that had succumbed to infection in control as well as in CV3-25 cohorts **(Figure 4J, K)**. Mice that survived in the CV3-25 cohorts, regained body weight and at 22 dpi, had no detectable virus in organs and base-line inflammatory cytokine induction **(Figure 4E-K)**. Overall, our data indicated that CV3-1 alone, was sufficient to inhibit establishment of virus infection and prophylactically protect K18-hACE2 mice. Histology of brain tissue revealed that NAbs CV3-1 and CV3-25 persisted even at 6 dpi **(Figure S3E)**. However, while CV3-1 localization remained unaltered post-infection due to the absence of viral neuroinvasion in this cohort, CV3-25 localized heavily onto the surface of infected neurons at 6 dpi, consistent with virus infection in the brain **(Figure S3E)**. Additionally, neutrophils infiltrated the brain in mice treated with CV3-25 alone **(Figure S3F)**. These data, together with the imaging analyses indicated that CV3-1 inhibited virus dissemination rather than neutralizing virus in peripheral tissues. In *in vivo* dose response studies, just 0.75 mg CV3-1/kg body weight was sufficient to afford 50% efficacy against lethal SARS-CoV-2 infection **(Figure S4)**. These data indicated that CV3-1 is highly potent at halting SARS-CoV-2 at early infection sites and was primarily responsible for protecting the cohort treated with the NAb cocktail.

### CV3-1 Therapy Rescues Mice from Lethal SARS-CoV-2 Infection

We explored if CV3-1 could also cure mice infected with SARS-CoV-2-nLuc. Mice were administered CV3-1 at 1, 3, and 4 dpi after confirming SARS-CoV-2 infection was established in the lungs **(Figure 5A)**. Temporal imaging and quantification of nLuc signal revealed that CV3-1, administered at 1 and 3 dpi, controlled virus spread successfully preventing neuroinvasion **(Figure 5B-D, G, H)**. This was corroborated by the absence of weight loss and/or recuperation of body weight, undetectable viral loads, and near-baseline levels of inflammatory cytokines in tissues **(Figure 5E-K)**. CV3-1 therapy at 4 dpi, however, could neither control virus spread nor neuroinvasion resulting in 75% mortality of the cohort **(Figure 5B-F)** with loss in body weight, high inflammatory cytokines levels and tissue viral loads, similar to the control cohort **(Figure 5E-K)**. Thus, the therapeutic window for maximal efficacy of CV3-1 treatment extends for up to 3 days from the initiation of SARS-CoV-2 infection.

**Figure 5.**
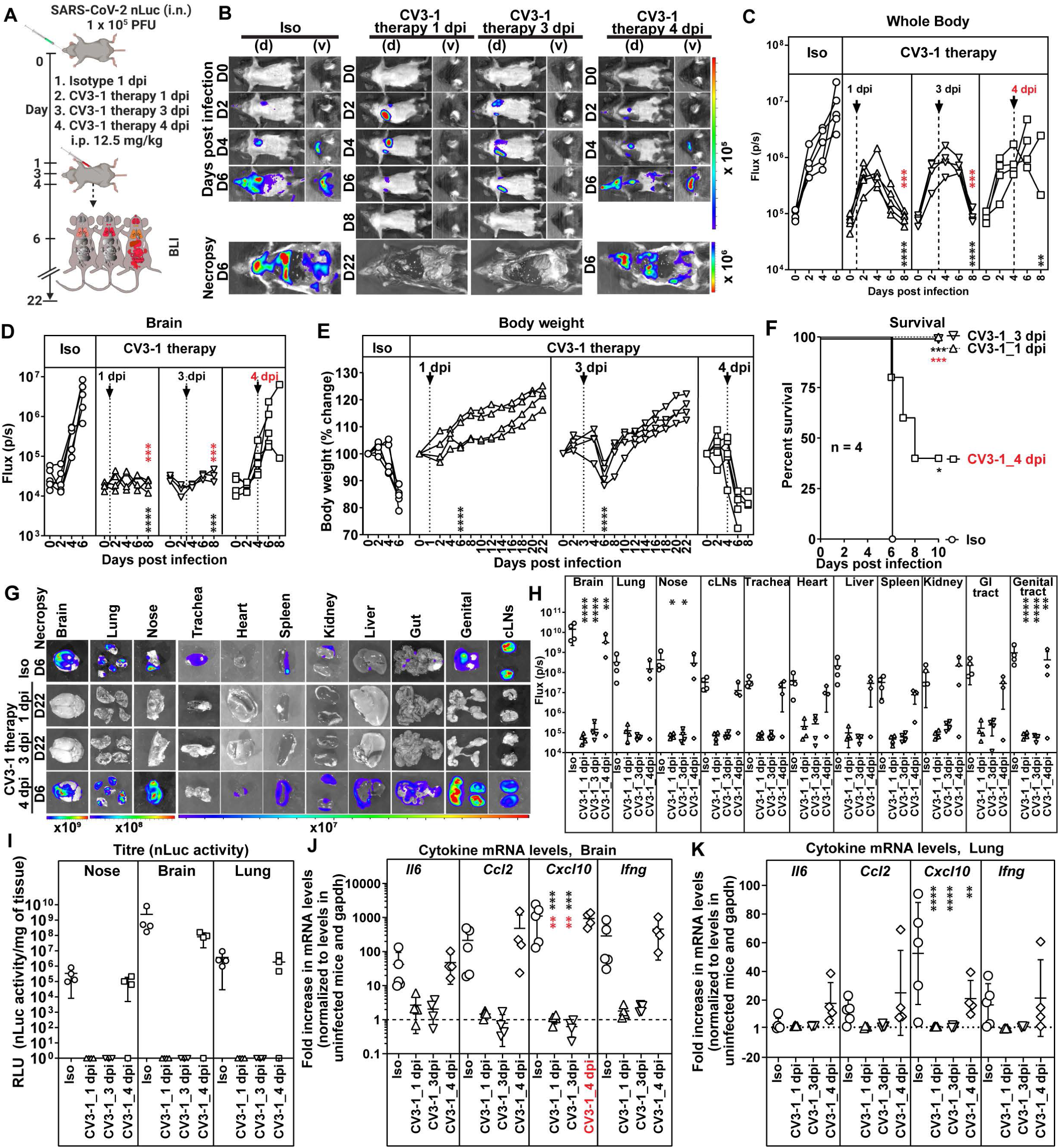
CV3-1 Therapy Protects Mice from Lethal SARS-CoV-2 Infection. (A) Experimental design to test *in vivo* efficacy of CV3-1 administered i.p. (12.5 mg/kg body weight) at indicated times after i.n. challenge of K18-hACE2 mice with SARS-CoV-2 nLuc followed by non-invasive BLI every 2 days. Human IgG1 treated (12.5 mg/kg body weight) mice were the control cohort (Iso). (B) Representative images from temporal BLI of SARS-CoV-2-nLuc-infected mice in ventral (v) and dorsal (d) positions. Scale bars denote radiance (photons/sec/cm^2^/steradian). (C-D) Temporal quantification of nLuc signal as flux (photons/sec) computed non-invasively. (E) Temporal changes in mouse body weight with initial body weight set to 100%. (F) Kaplan-Meier survival curves of mice statistically compared by log-rank (Mantel-Cox) test. (G, H) *Ex vivo* imaging of organs and quantification of nLuc signal as flux(photons/sec) after necropsy. (I) Viral loads (nLuc activity/mg tissue) measured in Vero E6 cells as targets. Non-detectable virus amounts were set to 1. (J, K) Fold changes in cytokine mRNA levels in lung and brain tissues normalized to GAPDH mRNA in the same sample and that in non-infected mice after necropsy. Viral loads (I) and inflammatory cytokine profile (J, K) were determined after necropsy at 6dpi. Each curve in (C)-(E) and each data point in (H)-(K) represents an individual mouse. CV3-1 treatment times are indicated in (C)-(E). Grouped data in (C)-(K) were analyzed by 2-way ANOVA followed by Dunnett’s or Tukey’s multiple comparison tests. Statistical significance for group comparisons to isotype control are shown in black and for groups under CV3-1 therapies to 4 dpi-treatment shown in red. ∗, p < 0.05; ∗∗, p < 0.01; ∗∗∗, p < 0.001; ∗∗∗∗, p < 0.0001; Mean values ± SD are depicted.

### CV3-1 and CV3-25 Require Antibody Effector Functions For *In vivo* Efficacy

Antibodies may also mediate Fc-recruitment of immune cells to eliminate infected cells (Lu et al., 2018). We explored a role for Fc-mediated effector functions in the *in vivo* protection afforded by NAbs. We generated mutant versions with Leucine to Alanine (L234A/L235A, LALA) changes of both NAbs to impair interaction with Fc receptors (Saunders, 2019) and a G236A/S239D/A330L/I332E (GASDALIE) version of CV3-25 known to enhance affinity to FcγRs. (Bournazos et al., 2014). LALA and GASDALIE mutations had no impact on S binding and neutralizing capabilities of NAbs **(Figure S5A, B)**. As expected, LALA mutations compromised ADCC and ADCP activities whereas CV3-25 GASDALIE displayed enhanced ADCC activity **(Figure S5C, D)**. Biodistribution analyses of AF_647_-conjugated mutants of CV3-1, CV3-25 24h after i.p. administration revealed penetration into most tissues **(Figure S5E)**.

We next tested the impact of Fc-effector altering mutations on the prophylactic efficacy of NAbs **(Figure S6A)**. Longitudinal non-invasive BLI and terminal imaging analyses after necropsy, body weight changes, survival and viral load estimations revealed that LALA mutations had indeed compromised efficacy of both antibodies **(Figure S6A-I)**. SARS-CoV-2 replicated better, invaded the brain and induced body weight loss in cohorts treated with LALA NAbs compared to the corresponding wildtype NAbs **(Figure S6D-E)**. Histology at 6 dpi revealed that both LALA NAbs had penetrated the brain tissue during the course of infection and bound the surface of infected neurons **(Figure S5F, G)**. At 6 dpi, both the CV3-1 LALA and CV3-25 LALA cohorts had higher tissue viral loads compared to the respective wildtype cohorts indicating compromised protective efficacy **(Figure S6G)**. Similarly, while tissue viral loads in CV3-25-pretreated mice were reduced by a log, those in CV3-25 LALA-pretreated mice were comparable to that in control cohorts. The delayed mortality and 25% protective efficacy offered by CV3-25 was abrogated and the protective efficacy of CV3-1 fell from 100% to 62.5% with the corresponding LALA mutants **(Figure S6F)**. There was also an overall increase in the inflammatory cytokine signature in the LALA cohorts **(Figure S6H, I)**. The requirement for Fc effector function for CV3-1 prophylaxis was surprising as no infection was detected in CV3-1-pretreated mice both by non-invasive and post-necropsy tissue imaging at 6 dpi **(Figure 4)**. However, examination of tissues at 3 dpi did reveal weak nLuc signals in the nasal cavity and lungs despite absence of signal by non-invasive imaging **(Figure S5H-M)**. PCR analyses also confirmed the presence SARS-CoV-2 N RNA in these tissues at 3 dpi **(Figure S5M)**. Thus, some incoming viruses managed to establish infection despite CV3-1 prophylaxis and Fc effector functions were required to eliminate infected cells. Correspondingly, Fc-effector enhanced CV3-25 GASDALIE reduced virus dissemination and provided 100% protective efficacy despite a transient loss in body weight in mice compared to the wild-type CV3-25 **(Figure S6B-F)**. Viral titers and inflammatory cytokines were also significantly reduced compared to control cohorts **(Figure S6G-I)**. We also tested the requirement for Fc- effector functions using mouse-adapted SARS-CoV-2 MA10 in WT B6 mice (Dinnon et al., 2020; Leist et al., 2020), an alternative model where mice succumb due to ARDS. CV3-1 successfully neutralized SARS-CoV-2 MA10 with similar efficacies to the WA1 strain *in vitro* (IC_50_ = 0.01837 µg/mL). Though we had to use 5-fold more virus and older B6 mice (12-14 weeks) to induce mortality, we observed a similar Fc effector requirement for CV3-1 to prophylactically protect mice and reduce viral loads and inflammation in lung and brain **(Figure S6J-O)**. The requirement for Fc-effector function for CV3-1 prophylaxis suggested that the same should be critical for CV3-1 therapy **(Figure 6A).** Indeed, while CV3-1 treatment at 3 dpi controlled infection, cohorts treated with CV3-1 LALA displayed rapidly spreading lung infection and fully succumbed by 6 dpi after an accelerated loss in body weight **(Figure 6B-F)**. High viral loads and cytokine levels in tissues also reflected the failure of the LALA NAbs to treat pre-established viral infection **(Figure 6G)**. Notably, while the lung viral loads in CV3-1 LALA cohorts were similar to that in the control, inflammatory cytokine mRNA levels in lungs, *CXCL10* in particular, were significantly higher suggesting a crucial requirement for Fc-engagement in curbing a cytokine-storm like phenotype **(Figure 6H-I).**

**Figure 6.**
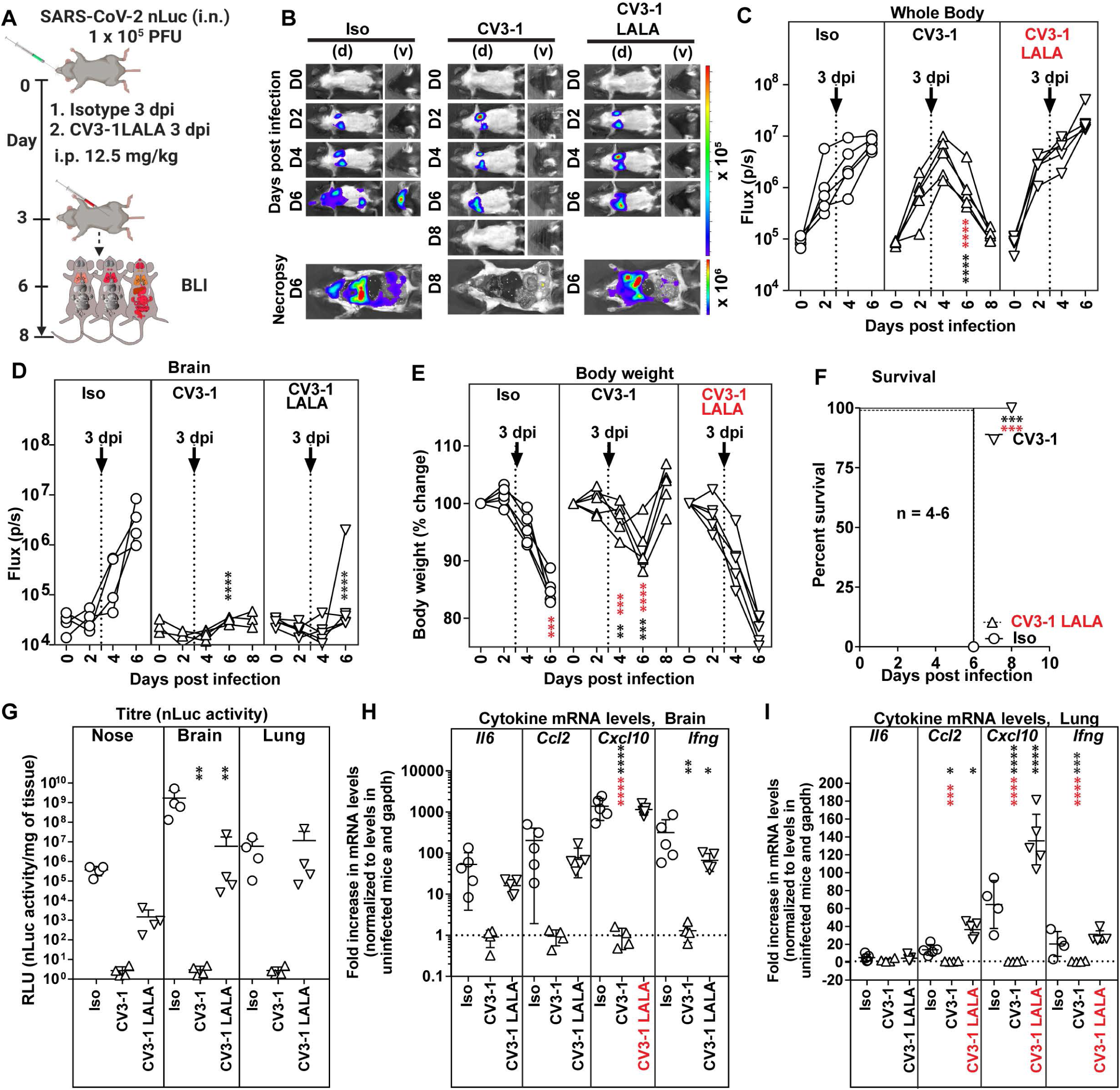
Fc-mediated Antibody Effector Functions Contribute to the *In Vivo* Efficacy of CV3-1. (A) Experimental design to test therapeutic efficacy of NAb CV3-1 and its corresponding Leucine to Alanine (LALA) mutant administered ip (12.5 mg/kg body weight) in K18-hACE2 mice 3 dpi with SARS-CoV-2 nLuc followed by non-invasive BLI every 2 days. Human IgG1-treated (12.5 mg/kg body weight) mice were used as the control cohort (Iso). (B) Representative images from temporal BLI of SARS-CoV-2-nLuc-infected mice in ventral (v) and dorsal (d) positions. Scale bars denote radiance (photons/sec/cm^2^/steradian). (C-D) Temporal quantification of nLuc signal as flux (photons/sec) computed non-invasively. (E) Temporal changes in mouse body weight with initial body weight set to 100%. (F) Kaplan-Meier survival curves of mice statistically compared by log-rank (Mantel-Cox) test. (G) Viral loads (nLuc activity/mg tissue) measured in Vero E6 cells as targets. Non-detectable virus amounts were set to 1. (H, I) Fold changes in cytokine mRNA levels in lung and brain tissues normalized to *Gapdh* mRNA in the same sample and that in non-infected mice after necropsy. Viral loads (G) and inflammatory cytokine profile (H, I) were determined after necropsy at 6dpi. Each curve in (C)-(E) and each data point in (G)-(I) represents an individual mouse. CV3-1 treatment times are indicated in (C)-(E). Grouped data in (C)-(I) were analyzed by 2-way ANOVA followed by Dunnett’s or Tukey’s multiple comparison tests. Statistical significance for group comparisons to isotype control are shown in black and between CV3-1 and CV3-1 LALA treated cohorts are shown in red. ∗, p < 0.05; ∗∗, p < 0.01; ∗∗∗, p < 0.001; ∗∗∗∗, p < 0.0001; Mean values ± SD are depicted.

### Monocytes, Neutrophils and Natural Killer (NK) cells Contribute to Antibody-mediated Effector Functions *In Vivo*

Fc can recruit NK cells, monocytes, or neutrophils to facilitate clearance of infected cells and shape the cytokine response produced by these cells for enhancing adaptive and cell-mediated immune responses (Lu et al., 2018). When NK cells were depleted prior to CV3-1 prophylaxis **(Figure S7J, K),** weak nLuc signals appeared in lungs of infected mice; however, this did not progress to the levels seen in control cohorts with no CV3-1 treatment **(Figure S7A-D)**. They also experienced a temporary but significant decrease in body weight **(Figure S7E)**. Nevertheless, NK cell depletion did not decrease the survival statistics of CV3-1 prophylaxis and all the mice survived despite marginal increases in viral loads in target organs and significant increases in inflammatory cytokine levels **(Figure S7F-K)**. Thus, while NK cells do contribute to *in vivo* efficacy of CV3-1, their absence did not compromise the protection offered by CV3-1 prophylaxis.

In therapeutic regimen format, when CV3-1 treatment was initiated at 3 dpi **(Figure 7A),** depleting NK cells compromised protective efficacy with 25% of the mice succumbing to SARS-CoV-2 infection compared to a 100% survival rate of mice cohorts replete with NK cells **(Figure 7B-F, S7J, K)**. Depleting Ly6G^+^ neutrophils and Ly6C^hi^ CD11b^+^ classical monocytes under CV3-1 therapy. resulted in 75% and 80% of the mouse cohorts respectively failing to control SARS-CoV-2 spread, with loss in body weight resulting in death **(Figure 7A-G, S7L-O)** (Mack et al., 2001). This was accompanied by increased viral burden and enhanced expression of *CCL2*, *CXCL10*, and *Il6* mRNA in target tissues **(Figure 7H-J)**. Overall, neutrophils, monocytes and NK cells contributed to the antibody-dependent cure of mice from lethal SARS-CoV-2 infection and were critical for the success of SARS-CoV-2 NAb-directed therapies.

**Figure 7.**
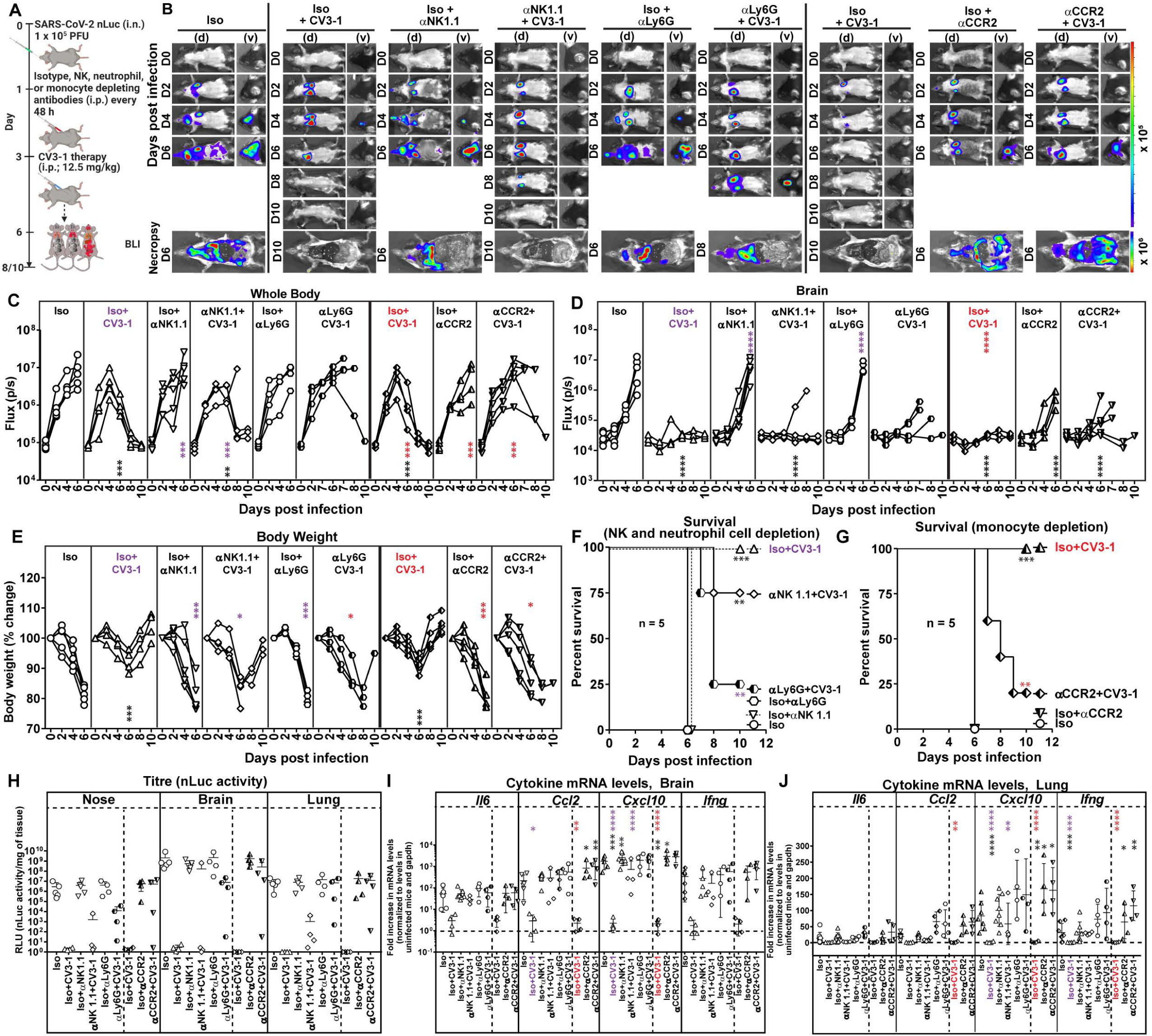
Monocytes, Neutrophils and Natural Killer Cells Contribute to Antibody Effector Functions *In Vivo*. (A) Experimental design to test the contribution of NK cells, neutrophils (CD11b^+^Ly6G^+^) and monocytes (CCR2^+^Ly6^hi^ CD11b^+^) in K18-hACE2 mice therapeutically treated with CV3-1 NAb (i.p.,12.5 mg/kg body weight) at 3 dpi after challenge with SARS-CoV-2-nLuc. αNK1.1, αLy6G and αCCR2 mAbs (i.p., 20, 20 and 2.5 mg/kg body weight respectively) were used to deplete NK cells, neutrophils and monocytes respectively every 48h starting at 1 dpi. Human and/or rat isotype mAb treated cohorts served as controls (Iso). The mice were followed by non-invasive BLI every 2 days from the start of infection. (B) Representative images from temporal BLI of SARS-CoV-2-nLuc-infected mice in ventral (v) and dorsal (d) positions. Scale bars denote radiance (photons/sec/cm^2^/steradian). (C-D) Temporal quantification of nLuc signal as flux (photons/sec) computed non-invasively. (E) Temporal changes in mouse body weight with initial body weight set to 100%. (F) Kaplan-Meier survival curves of mice statistically compared by log-rank (Mantel-Cox) test. (H) Viral loads (nLuc activity/mg tissue) measured in Vero E6 cells as targets. Non-detectable virus amounts were set to 1. (I, J) Fold change in cytokine mRNA levels in lung and brain tissues normalized to *GAPDH* mRNA in the same sample and that in non-infected mice after necropsy. Viral loads (H) and inflammatory cytokine profile (I, J) were determined after necropsy at 6dpi. Each curve in C-E and each data point in H-J represents an individual mouse. Grouped data in (C)-(I) were analyzed by 2-way ANOVA followed by Dunnett’s or Tukey’s multiple comparison tests. Statistical significance: group comparisons to isotype control are shown in black; group comparisons to Iso+CV3-1 within the NK and neutrophil depleted cohorts are shown in purple; group comparisons to Iso+CV3-1 within the monocyte-depleted cohorts are shown in red. ∗, p < 0.05; ∗∗, p < 0.01; ∗∗∗, p < 0.001; ∗∗∗∗, p < 0.0001; Mean values ± SD are depicted.

In summary, our data demonstrate the utility of a BLI-guided platform for temporo-spatial visualization of SARS-CoV-2 replication, pathogenesis and the mechanisms contributing to an effective outcome with NAb-based interventions in K18-ACE2 mice.

## Discussion

NAb therapies are being explored to augment current vaccination strategies against SARS-CoV-2 to expand the protection afforded towards emerging variants of concern. However, prior evidence for antibody-dependent enhancement of pathology caused by respiratory viruses like RSV and SARS-CoV-1 warrants careful investigation of antibody effects *in vivo* before clinical implementation (Iwasaki and Yang, 2020; Klasse and Moore, 2020). We have established a whole-body imaging approach to follow the dynamics and pathogenesis of SARS-CoV-2 infection in mice to facilitate preclinical studies for identifying effective therapeutic measures against COVID-19. Temporal tracking revealed that SARS-CoV-2 first replicates in the nasal cavity, reaches the lungs at 1 dpi where the infection expands till 3 dpi before spreading systemically to other organs including the brain at 4 dpi. BLI also helped illuminate how the highly potent human NAbs CV3-1 (targets Spike RBD) and CV3-25 (binds S2 domain) differed in their ability to protect or treat SARS-CoV-2 infection in the highly susceptible K18-hACE2 mouse model. Imaging analyses revealed widespread distribution of NAb within the animals, including in the nasal cavity and lungs where the virus infection is initially established, and persistence for at least a week after administration, features that were critical for efficacy in this acute model for SARS-CoV-2. BLI also revealed a therapeutic window of 3 dpi for CV3-1 NAb to successfully halt progression of infection from lungs to distal tissues. As previously reported SARS-CoV-2 NAbs have a therapeutic window of 1 dpi (Alsoussi et al., 2020; Hassan et al., 2020; Schafer et al., 2021; Winkler et al., 2021), CV3-1 displays one of the most potent *in vivo* efficacy profiles with a broad therapeutic window till 3 dpi. Most protective human NAbs for SARS-CoV-2 tested in animal models and in humans, target RBD (Baum et al., 2020; Chen et al., 2021; Rogers et al., 2020; Schafer et al., 2021; Tortorici et al., 2020; Weinreich et al., 2021), some NTD targeting NAbs have also display potent antiviral activity *in vivo* (Li et al., 2021; Noy-Porat et al., 2021; Voss et al., 2020). We show that the S2-directed CV3-25 NAb also conferred protection, albeit not as potently as CV3-1. This of significance as newly emerging variants display fewer mutations in the S2 subunit compared to the S1. Indeed, CV3-25 can efficiently neutralize the B.1.351 variant, while neutralization by anti-NTD and anti-RBD NAbs was greatly diminished (Stamatatos et al., 2021) Hence epitopes in S2 targeted by CV3-25 can be explored to generate future antigenic templates for potent pan-coronavirus antibodies (Sauer et al., 2021).

Our data also establishes that neutralizing capacity of NAbs alone is insufficient to garner clinical protection. LALA variants of CV3-1 revealed a crucial role for Fc-mediated interactions in augmenting *in vivo* protection not only for therapy, but also in prophylaxis in contrast to a recent report where Fc-effector was involved only during NAb therapy (Winkler et al., 2021). CV3-1 Fc effector functions were needed to eliminate infected cells originating from viruses that eluded neutralization during prophylaxis in both the B6 mouse model with mouse-adapted SARS-CoV-2 MA10 and in K18-hACE2. However, in agreement with previous study (Winkler et al., 2021), diminishing Fc function of CV3-1 completely compromised its ability to therapeutically cure mice. Surprisingly, body weight loss and inflammatory responses (CCL2, CXCL10, IFNγ) were aggravated in mice administered CV3-1 LALA therapeutically compared to control cohorts. Thus, the Fc region plays an additional protective role in limiting immunopathology by dampening inflammatory responses. A previously reported NAb engaged only monocytes for *in vivo* activity (Winkler et al., 2021). In contrast, CV3-1 engaged Fc-interacting neutrophils, monocytes and NK cells for its *in vivo* efficacy. Thus, in addition to potent neutralizing activity, effective engagement of innate immune components contributed to the high *in vivo* potency of CV3-1.

CV3-1, at low doses, did not enhance infection but displayed protective efficacy of only 50%. Thus, our data add to the growing body of evidence suggesting absence of an ADE mechanism during SARS-CoV-2 infection with a protective rather than pathogenic role for Fc (Schafer et al., 2021; Winkler et al., 2021). However, as the expression pattern of murine FcγRs on immune cells differs from that in humans, additional investigations, in other animal models, are required to confirm definitive absence of ADE in SARS-CoV-2 infection (Gorman et al., 2021). Moreover, elucidation of the major FcγR(s) (FcγRI, FcγRIII and/or FcγRIV) engaged by NAbs will help design ultrapotent SARS-CoV-2 Nab therapies (Smith et al., 2012). Here we have taken a step in this direction by introducing GASDALIE mutations to augment FcγR interactions and enhance *in vivo* potency of CV3-25.

In summary, our study demonstrates the utility of the BLI-guided approach to study SARS-CoV-2 pathogenesis and identify effective antiviral therapies for rapid translation to clinical use in humans.

## Supplemental information

7 Supplementary figures, 1 table and 5 Videos

## Author contributions

Conceptualization, PDU, PK, AF, WM, IU and JP; Methodology, PDU, IU, JP, MSL, ML.WM, AF, PK; Investigation, IU, PDU, JP, HS, KS, LS, ATM, SPA, GBB, MB, SD, RG, CF, YC, AT, GG, CB, HM, GAD, JDD, DEK, JR, MP; Writing – Original Draft, PDU; Writing – Review & Editing, PDU, PK, AF, WM, IU, JP, LS, ATM; Funding Acquisition, AF, LS, ATM, PJB; Resources, AF, PJB, CBW, MM; Supervision, PDU, WM, PK, AF, PJB.

## Acknowledgements

This work was supported by George Mason University Fast Grants to MSL and PJB; P20GM125498 (awarded to UVM Translational Global Infectious Disease Research Center) to EAB; le Ministère de l’Économie et de l’Innovation du Québec, Programme de soutien aux organismes de recherche et d’innovation, Foundation du CHUM, Canadian Institutes of Health Research (CIHR) foundation grant #352417 & Rapid Research Funding Opportunity #FRN440388 to JDD and G.A.D, Canada Research Chair on Retroviral Entry no. RCHS0235 950-232424 to AF; Canada’s COVID-19 Immunity Task Force (CITF) & Canada Foundation for Innovation (CFI) #41027 to AF and DEK & #36287 to JDD. and GAD; FRQS Merit Research Scholarship to DEK; CIHR fellowships to JP, SPA and GBB, MITACS Accélération postdoctoral fellowship to RG; Fred Hutch COVID-19 Research Fund to LS and ATM.

## Disclaimer

The views expressed in this presentation are those of the authors and do not reflect the official policy or position of the Uniformed Services University, US Army, the Department of Defense, or the US Government.

## STAR Methods

### KEY RESOURCES TABLE

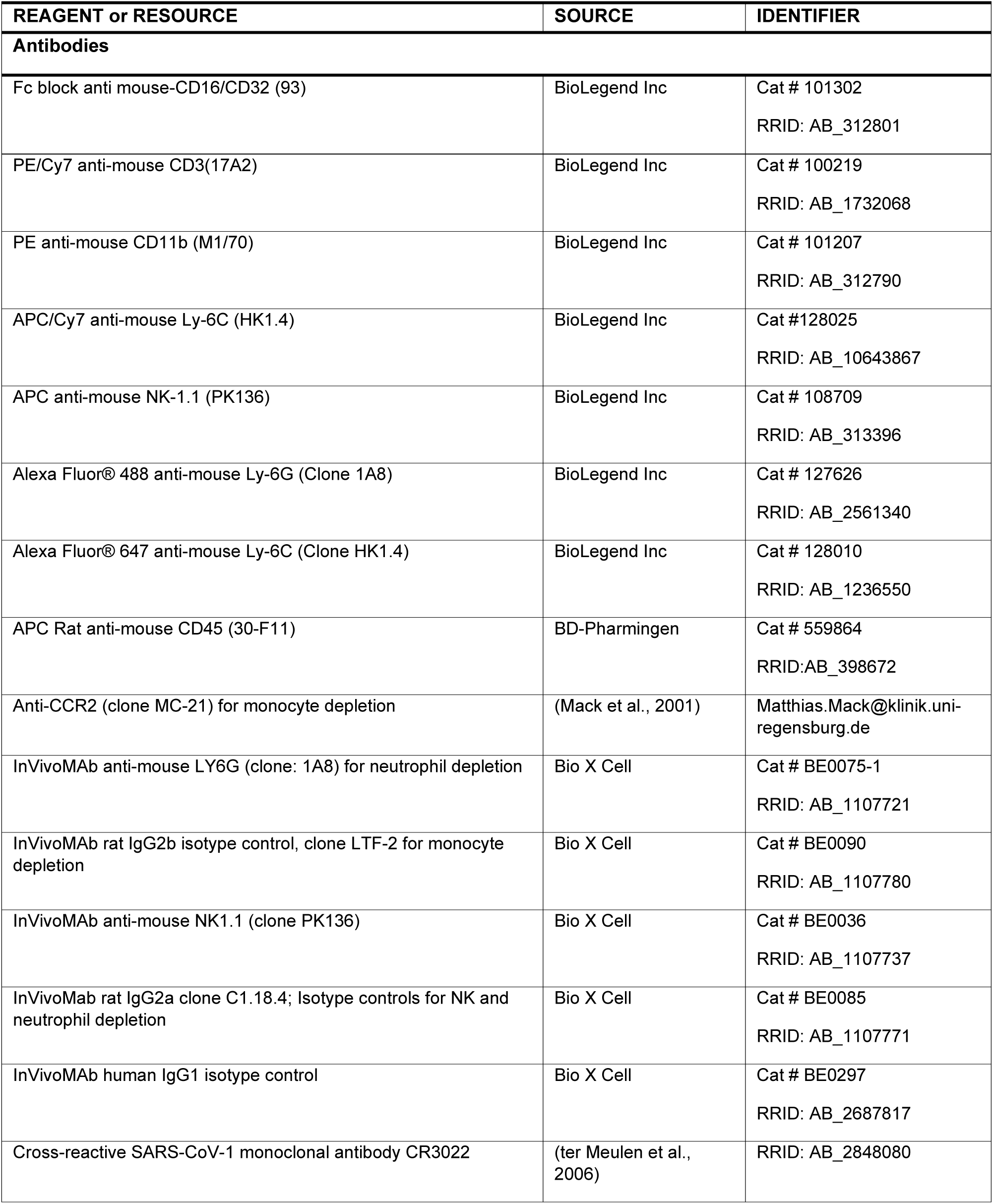
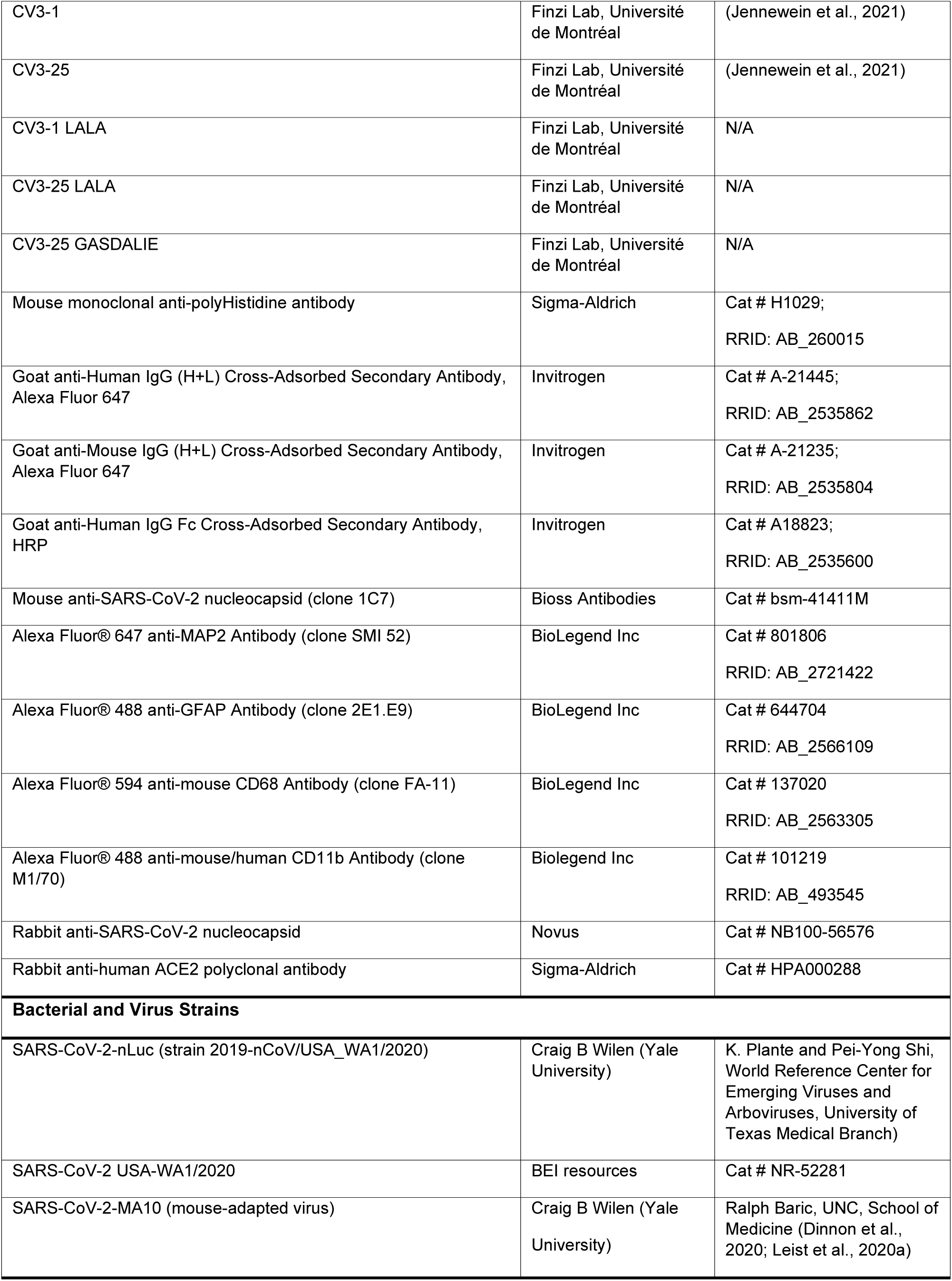
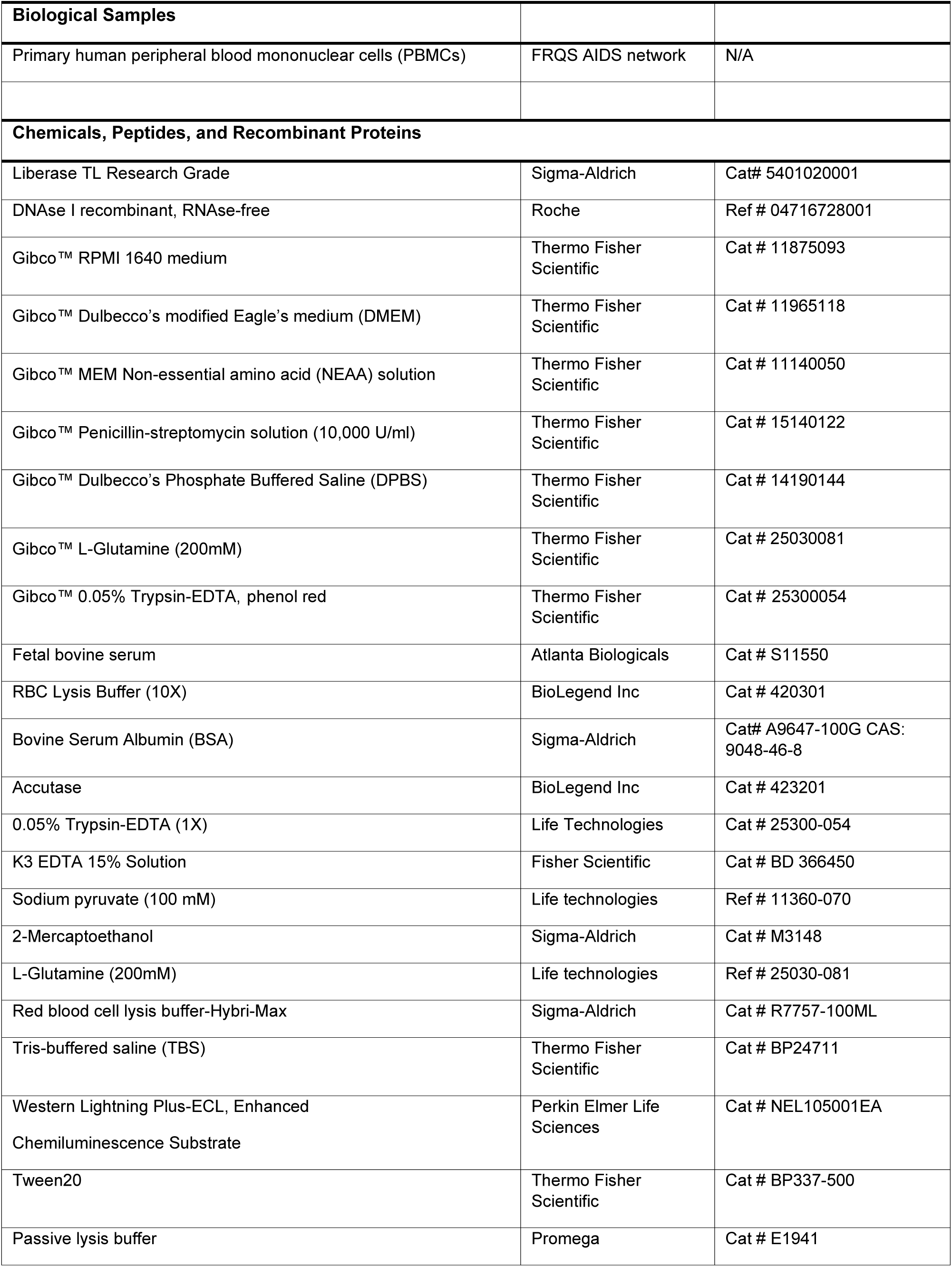
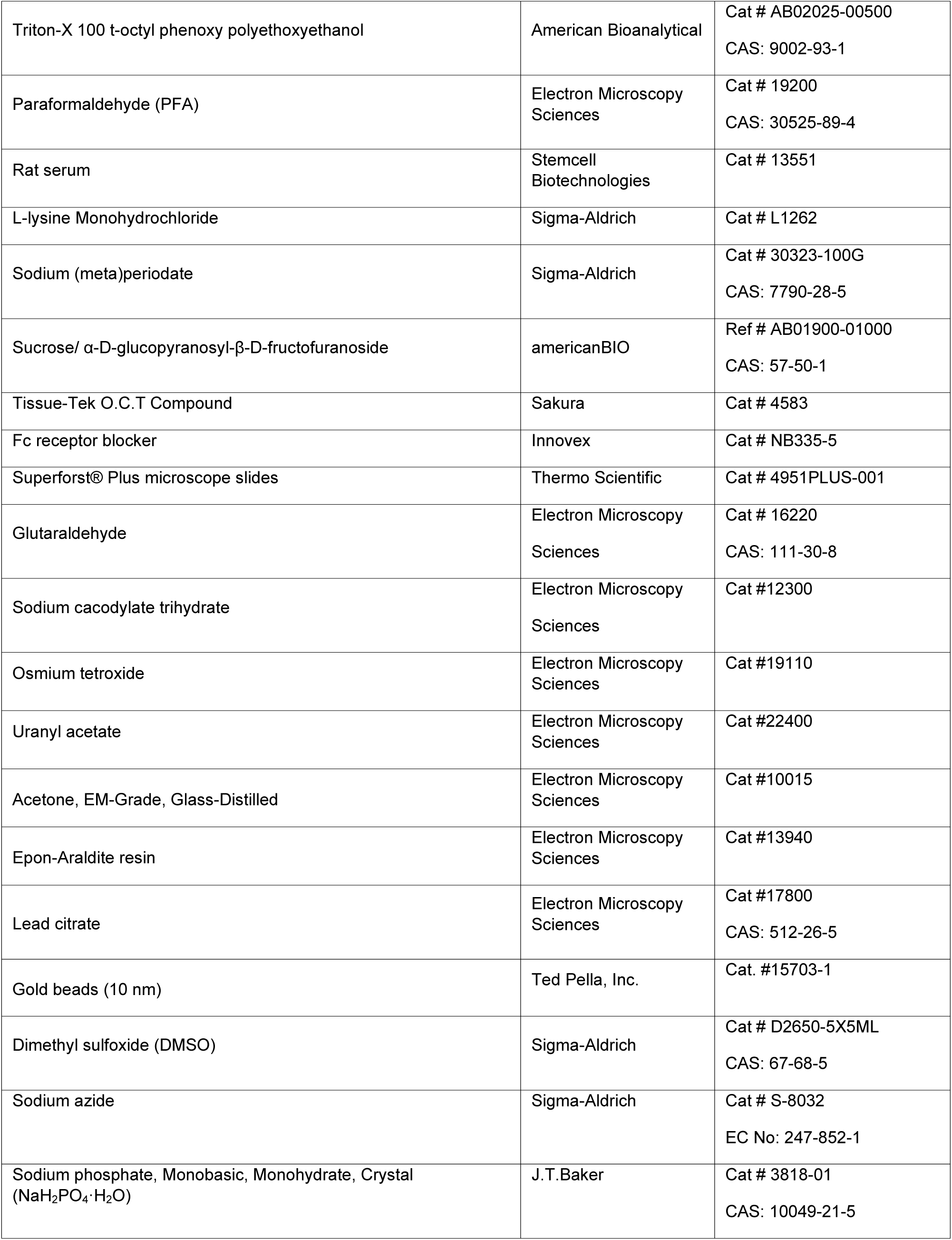
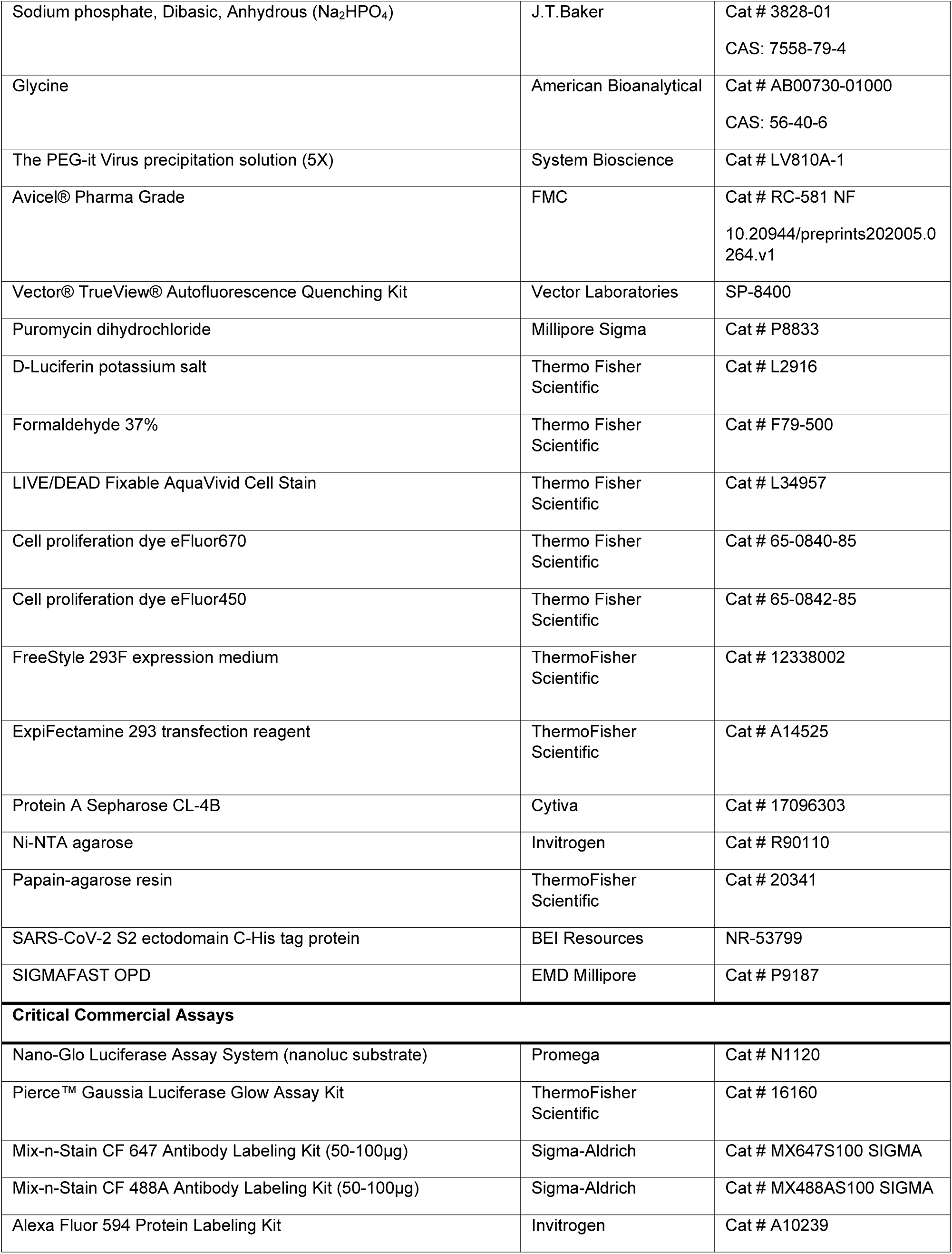
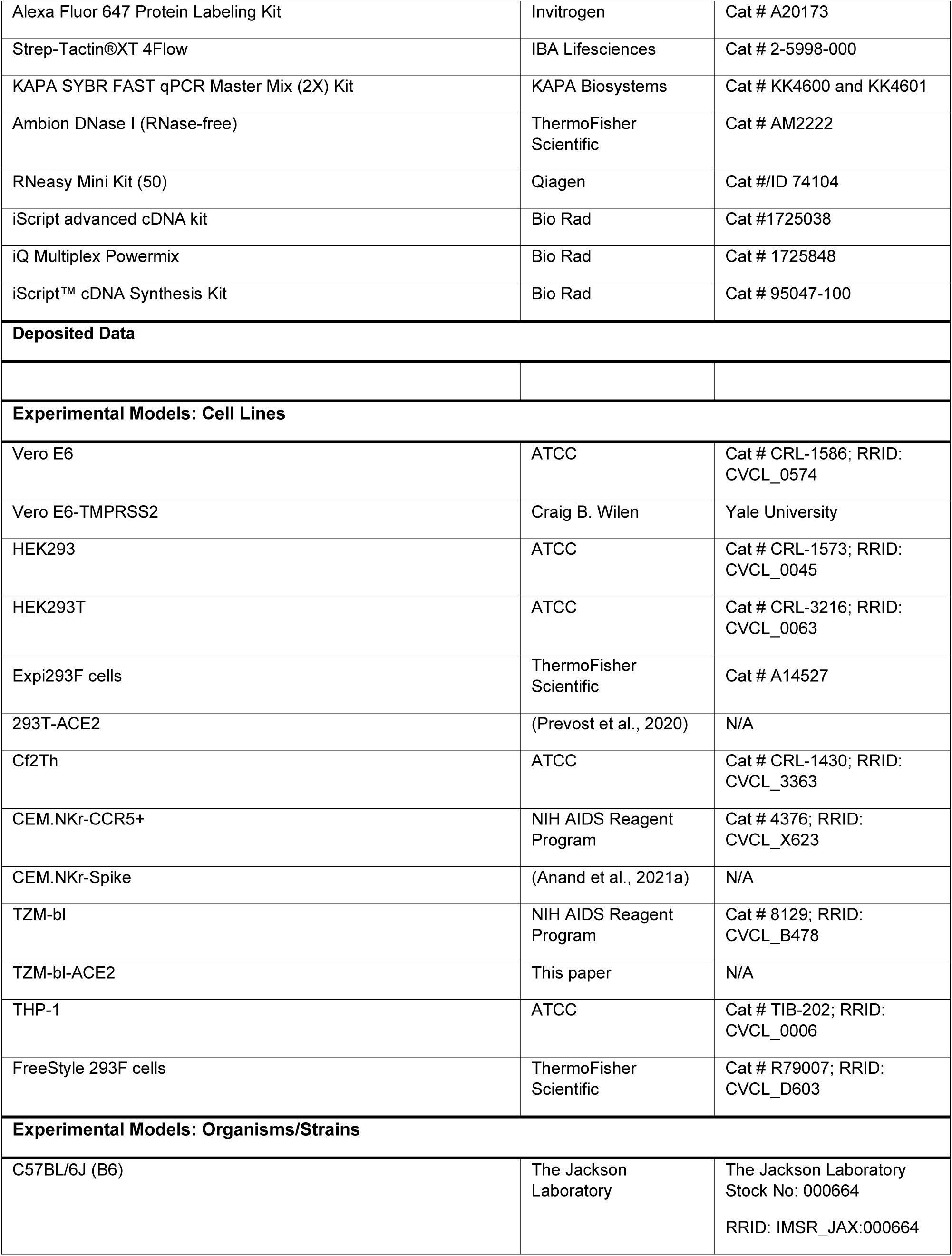
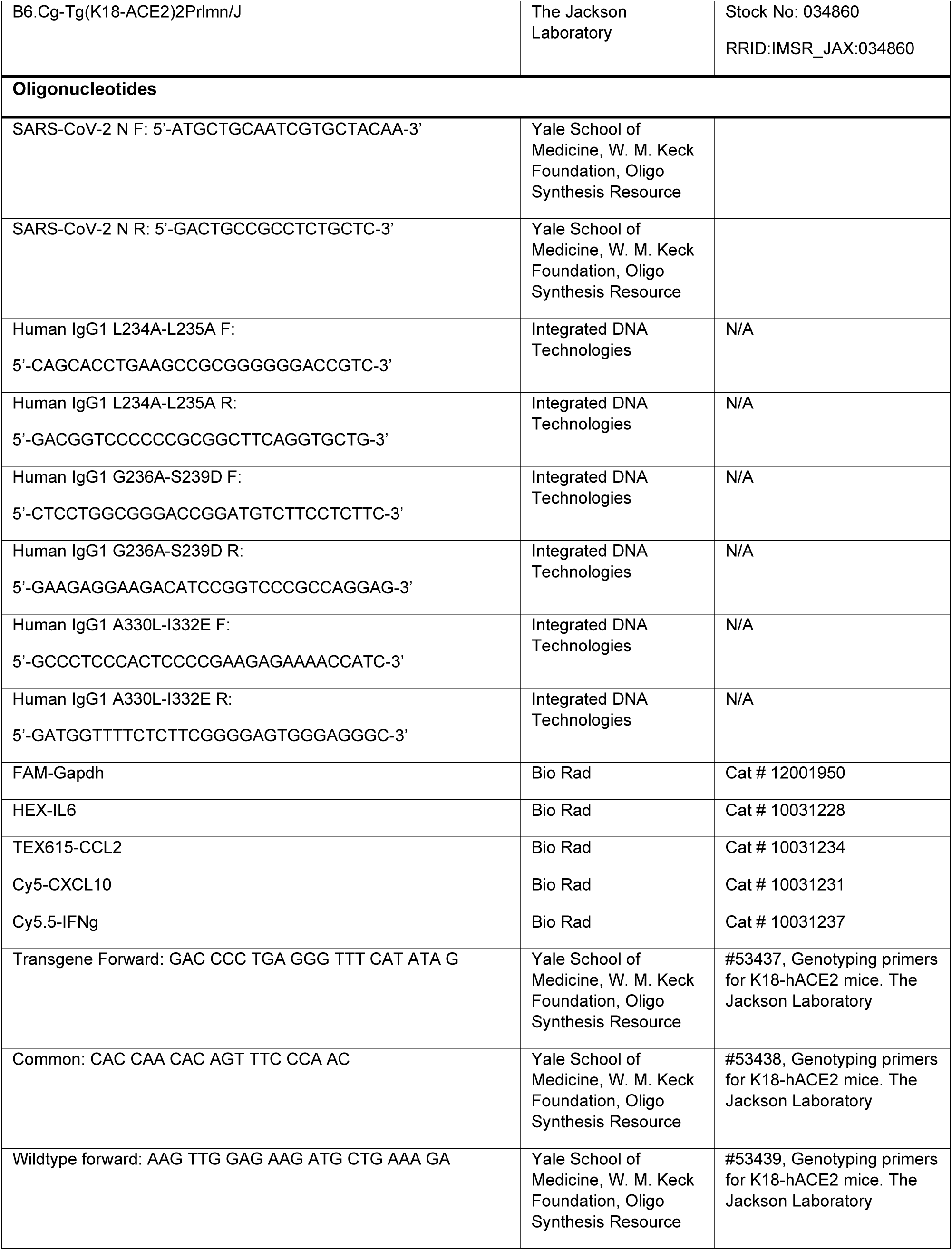
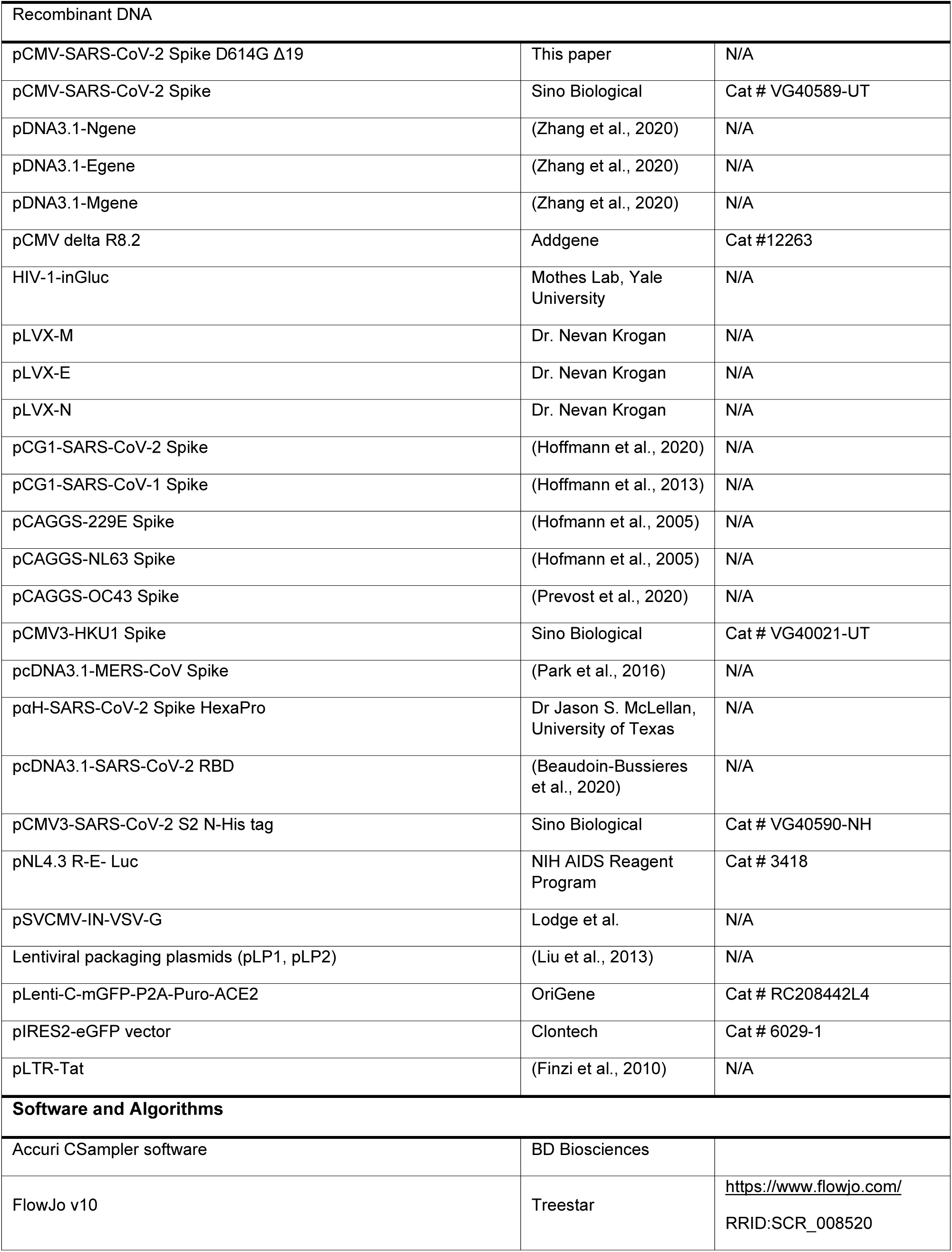
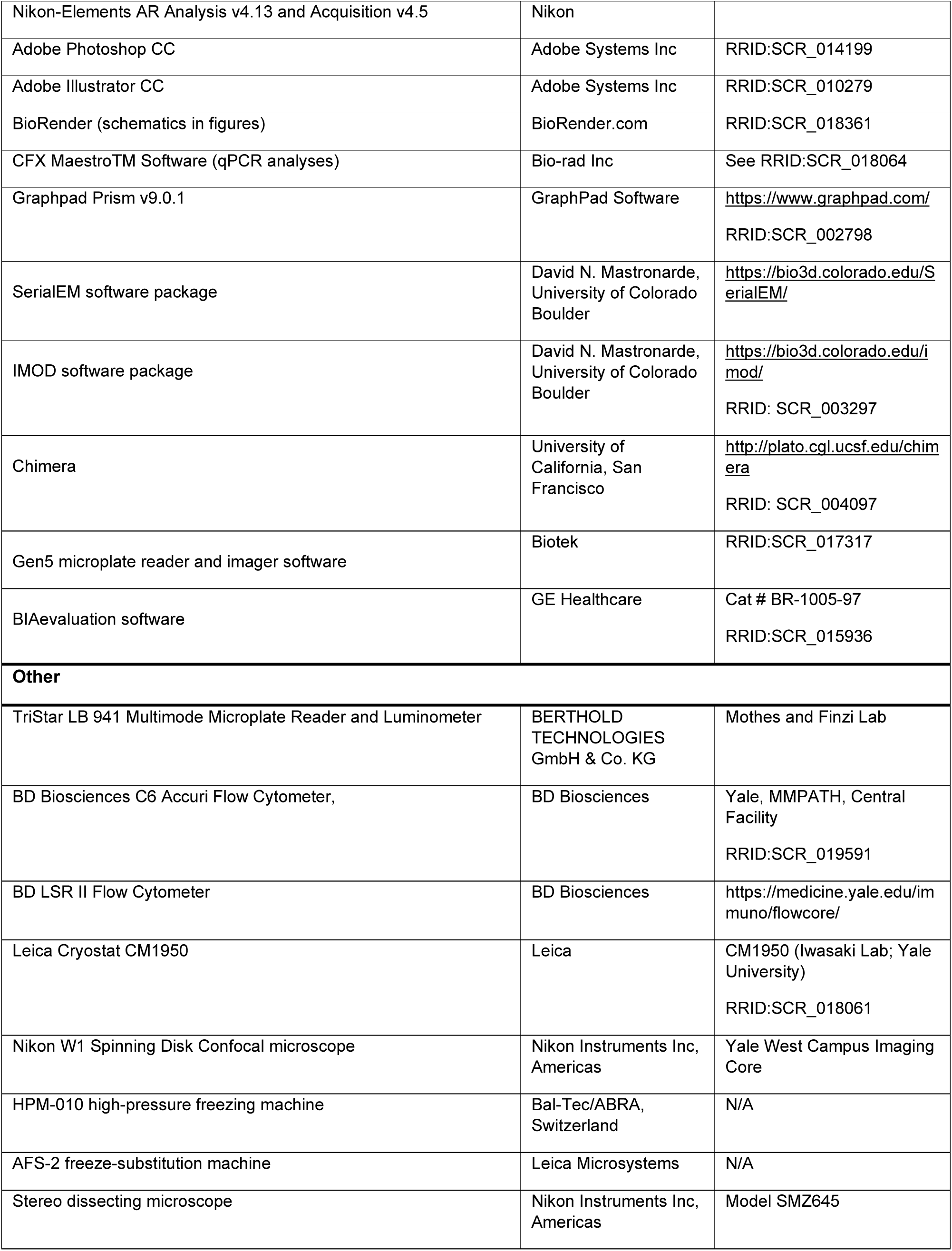
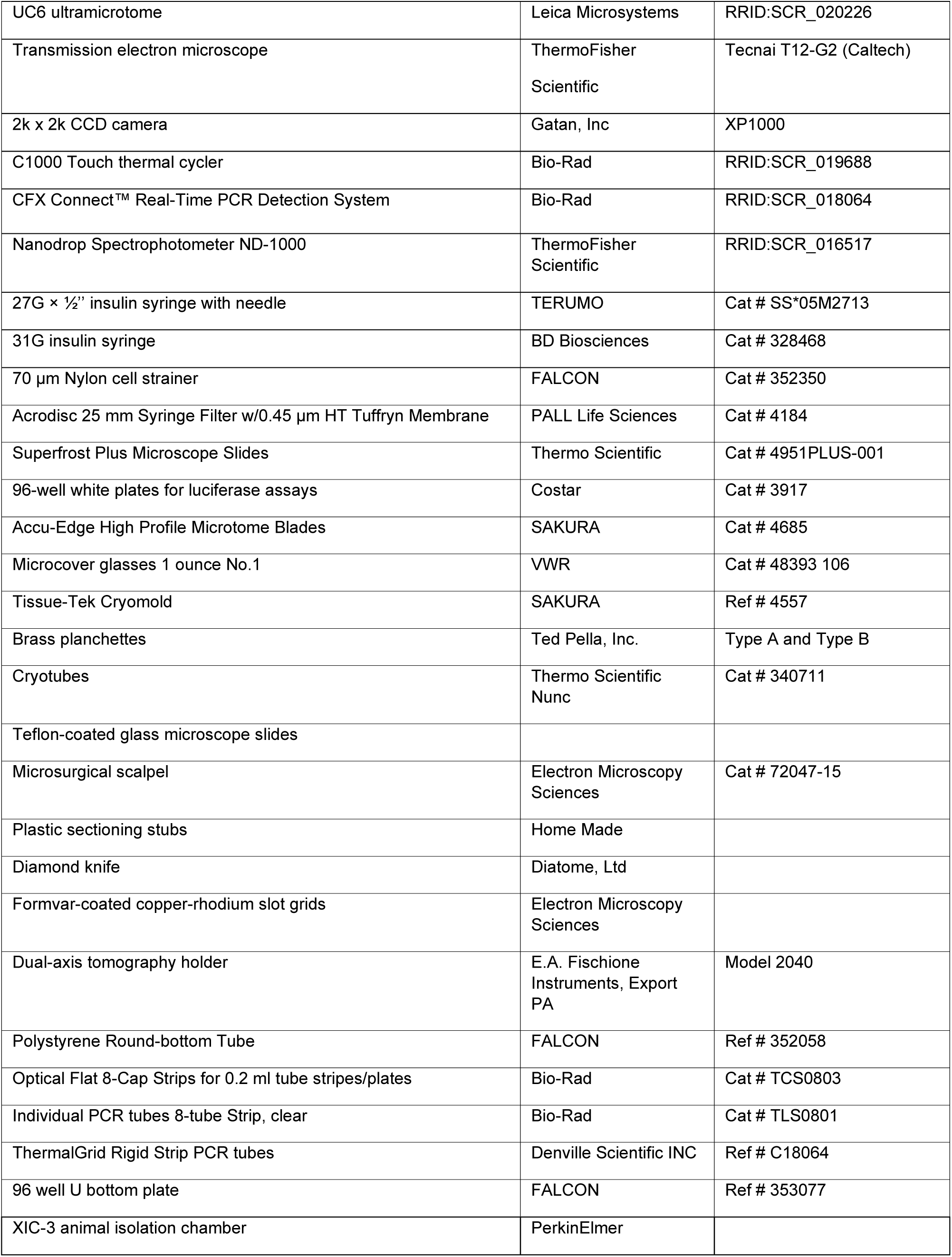
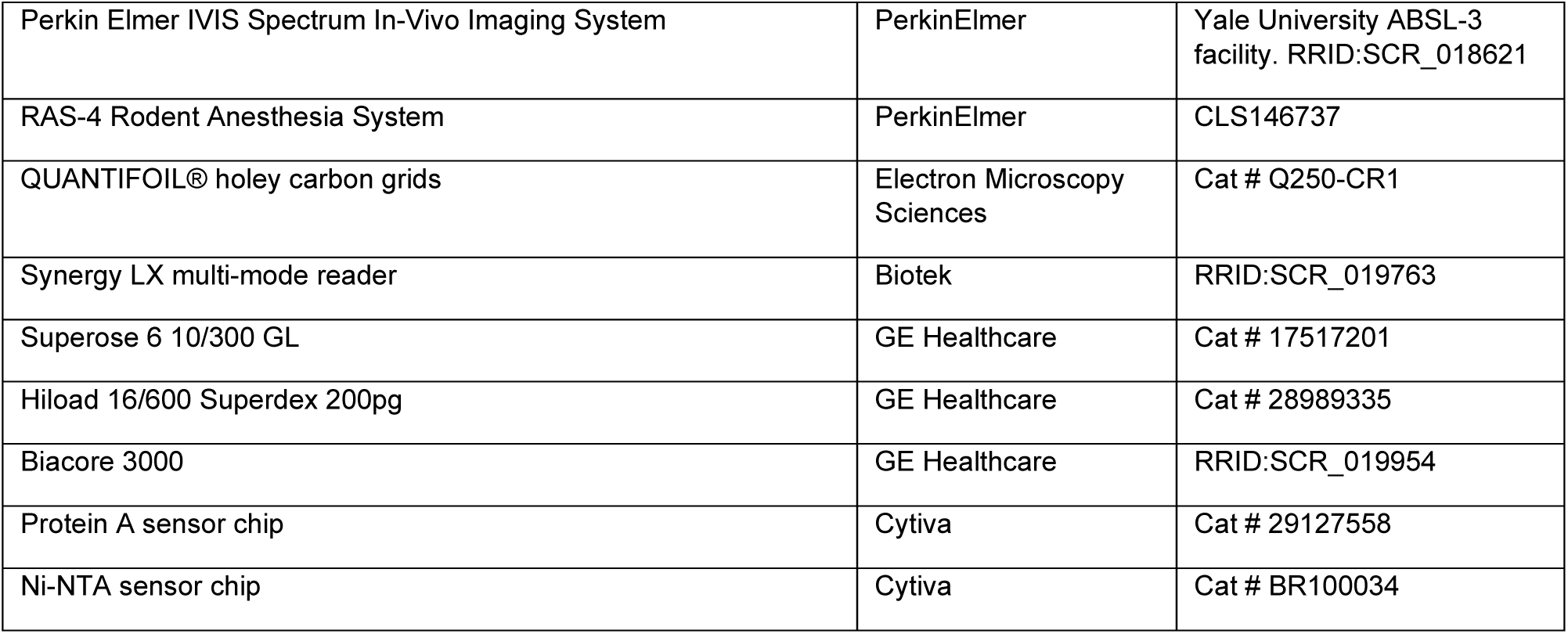

### RESOURCE AVAILABILITY

**Lead Contact** Pradeep Uchil (

Requests for resources and reagents should be directed to and will be fulfilled by the Lead Contact, Pradeep Uchil, Priti Kumar, Andrés Finzi and Walther Mothes.

#### Materials Availability

All other unique reagents generated in this study are available from the corresponding authors with a completed Materials Transfer Agreement.

#### Data and Code Availability

All the data that support the findings of this study are available from the corresponding authors upon reasonable request.

### EXPERIMENTAL MODEL AND SUBJECT DETAILS

#### Cell and Viruses

Vero E6 (CRL-1586, American Type Culture Collection (ATCC), were cultured at 37°C in RPMI supplemented with 10% fetal bovine serum (FBS), 10 mM HEPES pH 7.3, 1 mM sodium pyruvate, 1× non-essential amino acids, and 100 U/ml of penicillin–streptomycin. The 2019n- CoV/USA_WA1/2019 isolate of SARS-CoV-2 expressing nanoluciferase was obtained from Craig B Wilen, Yale University and generously provided by K. Plante and Pei-Yong Shi, World Reference Center for Emerging Viruses and Arboviruses, University of Texas Medical Branch) (Xie et al., 2020a; Xie et al., 2020b). Mouse-adapted SARS-CoV-2 MA10 was obtained from Craig B. Wilen, Yale University and generously provided by Ralph S Baric, Department of Epidemiology, University of North Carolina at Chapel Hill (Leist et al., 2020). The SARS-CoV-2 USA-WA1/2020 virus strain used for microneutralization assay was obtained through BEI Resources. Viruses (WA1 or MA10) were propagated in Vero-E6 or Vero E6 TMPRSS2 by infecting them in T150 cm^2^ flasks at a MOI of 0.1. The culture supernatants were collected after 72 h when cytopathic effects were clearly visible. The cell debris was removed by centrifugation and filtered through 0.45-micron filter to generate virus stocks. Viruses were concentrated by adding one volume of cold (4 °C) 4x PEG-it Virus Precipitation Solution (40 % (w/v) PEG-8000 and 1.2 M NaCl; System Biosciences) to three volumes of virus-containing supernatant. The solution was mixed by inverting the tubes several times and then incubated at 4 °C overnight. The precipitated virus was harvested by centrifugation at 1,500 × g for 60 minutes at 4 °C. The concentrated virus was then resuspended in PBS then aliquoted for storage at −80°C. All work with infectious SARS-CoV-2 was performed in Institutional Biosafety Committee approved BSL3 and A-BSL3 facilities at Yale University School of Medicine or the University of Western Ontario using appropriate positive pressure air respirators and protective equipment. CEM.NKr, CEM.NKr-Spike, THP-1 and peripheral blood mononuclear cells (PBMCs) were maintained at 37°C under 5% CO_2_ in RPMI media, supplemented with 10% FBS and 100 U/mL penicillin/ streptomycin. 293T (or HEK293T), 293T-ACE2, CF2Th, TZM-bl and TZM-bl-ACE2 cells were maintained at 37°C under 5% CO_2_ in DMEM media, supplemented with 5 % FBS and 100 U/mL penicillin/ streptomycin. CEM.NKr (NIH AIDS Reagent Program) is a T lymphocytic cell line resistant to NK cell-mediated lysis. CEM.NKr-Spike stably expressing SARS-CoV-2 Spike were used as target cells in ADCC and ADCP assays (Anand et al., 2021a). THP-1 monocytic cell line (ATCC) was used as effector cells in the ADCP assay. PBMCs were obtained from healthy donor through leukapheresis and were used as effector cells in ADCC assay. 293T cells (obtained from ATCC) were derived from 293 cells, into which the simian virus 40 T-antigen was inserted. 293T-ACE2 cells stably expressing human ACE2 is derived from 293T cells (Prevost et al., 2020). Cf2Th cells (obtained from ATCC) are SARS-CoV-2-resistant canine thymocytes and were used in the virus capture assay. TZM-bl (NIH AIDS Reagent Program) were derived from HeLa cells and were engineered to contain the Tat-responsive firefly luciferase reporter gene. For the generation of TZM-bl cells stably expressing human ACE2, transgenic lentiviruses were produced in 293T using a third-generation lentiviral vector system. Briefly, 293T cells were co-transfected with two packaging plasmids (pLP1 and pLP2), an envelope plasmid (pSVCMV-IN-VSV-G) and a lentiviral transfer plasmid coding for human ACE2 (pLenti-C-mGFP-P2A-Puro-ACE2) (OriGene). Forty-eight hours post-transfection, supernatant containing lentiviral particles was used to infect TZM-bl cells in presence of 5 µg/mL of polybrene. Stably transduced cells were enriched upon puromycin selection. TZM-bl-ACE2 cells were then cultured in medium supplemented with 2 mg/mL of puromycin (Millipore Sigma).

#### Ethics statement

PBMCs from healthy individuals as a source of effector cells in our ADCC assay were obtained under CRCHUM institutional review board (protocol #19.381). Research adhered to the standards indicated by the Declaration of Helsinki. All participants were adults and provided informed written consent prior to enrollment in accordance with Institutional Review Board approval.

#### Antibodies

The human antibodies (CV3-1 and CV3-25) used in the work were isolated from blood of male convalescent donor S006 (male) recovered 41 days after symptoms onset using fluorescent recombinant stabilized Spike ectodomains (S2P) as probes to identify antigen-specific B cells as previously described (Lu et al., 2020; Seydoux et al., 2020; Jennewein et al., 2021). Site-directed mutagenesis was performed on plasmids expressing CV3-1 and CV3-25 antibody heavy chains in order to introduce the LALA mutations (L234A/L235A) or the GASDALIE mutations (G236A/S239D/A330L/I332E) using to the QuickChange II XL site-directed mutagenesis protocol (Stratagene).

#### Mouse Experiments

All experiments were approved by the Institutional Animal Care and Use Committees (IACUC) of and Institutional Biosafety Committee of Yale University (IBSCYU). All the animals were housed under specific pathogen-free conditions in the facilities provided and supported by Yale Animal Resources Center (YARC). All IVIS imaging, blood draw and virus inoculation experiments were done under anesthesia using regulated flow of isoflurane:oxygen mix to minimize pain and discomfort to the animals.

C57BL/6 (B6), hACE2 transgenic B6 mice (heterozygous) were obtained from Jackson Laboratory. 6–8-week-old male and female mice were used for all the experiments. The heterozygous mice were crossed and genotyped to select heterozygous mice for experiments by using the primer sets recommended by Jackson Laboratory.

### METHOD DETAILS

#### SARS-CoV-2 infection and treatment conditions

For all *in vivo* experiments, the 6 to 8 weeks male and female mice were intranasally challenged with 1 x 10^5^ FFU in 25-30 µl volume under anesthesia (0.5 - 5 % isoflurane delivered using precision Dräger vaporizer with oxygen flow rate of 1 L/min). For mouse-adapted SARS-CoV-2 MA10 challenge experiments, 12–14-week-old male and female mice were used with a challenge dose of 5 x 10^5^ FFU delivered in 25-30 µl volume under anesthesia as above. For NAb treatment using prophylaxis regimen, mice were treated with 250 µg (12.5 mg/kg body weight) of indicated antibodies (CV3-1, CV3-25, CV3-1 LALA, CV3-25 LALA or CV3-25 GASDALIE) or in combination (CV3-1:CV3-25; 6.25 mg/kg body weight of each) via intraperitoneal injection (i.p.) 24 h prior to infection. For neutralizing mAb treatment under therapeutic regimen, mice were treated at 1, 3 and 4 dpi intraperitoneally with CV3-1 or 3 dpi with CV3-1 LALA (12.5 mg/kg body weight). Body weight was measured and recorded daily. The starting body weight was set to 100 %. For survival experiments, mice were monitored every 6-12 h starting six days after virus administration. Lethargic and moribund mice or mice that had lost more than 20 % of their body weight were sacrificed and considered to have succumbed to infection for Kaplan-Meier survival plots.

#### Bioluminescence Imaging (BLI) of SARS-CoV-2 infection

All standard operating procedures and protocols for IVIS imaging of SARS-CoV-2 infected animals under ABSL-3 conditions were approved by IACUC, IBSCYU and YARC. All the imaging was carried out using IVIS Spectrum® (PerkinElmer) in XIC-3 animal isolation chamber (PerkinElmer) that provided biological isolation of anesthetized mice or individual organs during the imaging procedure. All mice were anesthetized via isoflurane inhalation (3 - 5 % isoflurane, oxygen flow rate of 1.5 L/min) prior and during BLI using the XGI-8 Gas Anesthesia System. Prior to imaging, 100 µL of nanoluciferase substrate, furimazine (NanoGlo^TM^, Promega, Madison, WI) diluted 1:40 in endotoxin-free PBS was retroorbitally administered to mice under anesthesia. The mice were then placed into XIC-3 animal isolation chamber (PerkinElmer) pre-saturated with isothesia and oxygen mix. The mice were imaged in both dorsal and ventral position at indicated days post infection. The animals were then imaged again after euthanasia and necropsy by spreading additional 200 µL of substrate on to exposed intact organs. Infected areas of interest identified by carrying out whole-body imaging after necropsy were isolated, washed in PBS to remove residual blood and placed onto a clear plastic plate. Additional droplets of furimazine in PBS (1:40) were added to organs and soaked in substrate for 1-2 min before BLI.

Images were acquired and analyzed with the manufacturer’s Living Image v4.7.3 *in vivo* software package. Image acquisition exposures were set to auto, with imaging parameter preferences set in order of exposure time, binning, and f/stop, respectively. Images were acquired with luminescent f/stop of 2, photographic f/stop of 8. Binning was set to medium. Comparative images were compiled and batch-processed using the image browser with collective luminescent scales. Photon flux was measured as luminescent radiance (p/sec/cm2/sr). During luminescent threshold selection for image display, luminescent signals were regarded as background when minimum threshold levels resulted in displayed radiance above non-tissue-containing or known uninfected regions. To determine the pattern of virus spread, the image sequences were acquired every day following administration of SARS-CoV-2 (i.n). Image sequences were assembled and converted to videos using Image J.

#### Biodistribution of therapeutic neutralizing antibodies using IVIS

Mice were intraperitoneally (i.p) administered with 250 µg of unconjugated (12.5 mg/kg body weight), Alexa Fluor 647 or Alexa Fluor 594-labeled antibodies to non-infected or SARS-CoV-2 infected hACE2 mice. 24 h later all organs (nose, trachea, lung, cervical lymph nodes, brain, liver, spleen, kidney, gut, testis and seminal vesicles) were isolated after necropsy and images were acquired with an IVIS Spectrum® (PerkinElmer) and fluorescence radiance intensities were analyzed with the manufacturer’s Living Image v4.7.3 *in vivo* software package. Organs were cut into half and weighed. One half was fixed in 4 % PFA and processed for cryoimmunohistology. The other half was resuspended in serum-free RPMI and homogenized in a bead beater for determination of antibody levels using quantitative ELISA.

#### Measurement of therapeutic antibody levels in organs by quantitative ELISA

Recombinant SARS-CoV-2 RBD and S-6P proteins were used to quantify CV3-1 and CV3-25 antibody levels, respectively, in mice organs. SARS-CoV-2 proteins (2.5 μg/ml), or bovine serum albumin (BSA) (2.5 μg/ml) as a negative control, were prepared in PBS and were adsorbed to plates (MaxiSorp; Nunc) overnight at 4 °C. Coated wells were subsequently blocked with blocking buffer (Tris-buffered saline [TBS], 0.1% Tween20, 2% BSA) for 1 hour at room temperature. Wells were then washed four times with washing buffer (TBS 0.1% Tween20). Titrated concentrations of CV3-1 or CV3-25 or serial dilutions of mice organ homogenates were prepared in a diluted solution of blocking buffer (0.1 % BSA) and incubated in wells for 90 minutes at room temperature. Plates were washed four times with washing buffer followed by incubation with HRP-conjugated anti-IgG secondary Abs (Invitrogen) (diluted in a diluted solution of blocking buffer [0.4% BSA]) for 1 hour at room temperature, followed by four washes. HRP enzyme activity was determined after the addition of a 1:1 mix of Western Lightning oxidizing and luminol reagents (Perkin Elmer Life Sciences). Light emission was measured with a LB941 TriStar luminometer (Berthold Technologies). Signal obtained with BSA was subtracted for each organ. Titrated concentrations of CV3-1 or CV3-25 were used to establish a standard curve of known antibody concentrations and the linear portion of the curve was used to infer the antibody concentration in tested organ homogenates.

#### Cryo-immunohistology of organs

Organs were isolated after necropsy and fixed in 1X PBS containing freshly prepared 4% PFA for 12 h at 4 °C. They were then washed with PBS, cryoprotected with 10, 20 and 30% ascending sucrose series, snap-frozen in Tissue-Tek® O.C.T.^TM^ compound and stored at -80 °C. The nasal cavity was snap-frozen in 8% gelatin prepared in 1X PBS and stored at -80 °C. 10 - 30 µm thick frozen sections were permeabilized with 0.2% Triton X-100 and treated with Fc receptor blocker (Innovex Biosciences) before staining with indicated conjugated primary, secondary antibodies or Phalloidin in PBS containing 2% BSA containing 10% fetal bovine serum. Stained sections were treated with TrueVIEW Autofluorescence Quenching Kit (Vector Laboratories) and mounted in VECTASHIELD® Vibrance™ Antifade Mounting Medium. Images were acquired using Nikon W1 spinning disk confocal microscope equipped with 405, 488, 561 and 647 nm laser lines or EVOS M7000 imaging system. The images were processed using Nikon Elements AR version 4.5 software (Nikon Instruments Inc, Americas) and figures assembled with Photoshop CC and Illustrator CC (Adobe Systems, San Jose, CA, USA).

#### Focus forming assay

Titers of virus stocks was determined by standard plaque assay. Briefly, the 4 x 10^5^ Vero-E6 cells were seeded on 12-well plate. 24 h later, the cells were infected with 200 µL of serially diluted virus stock. After 1 hour, the cells were overlayed with 1ml of pre-warmed 0.6% Avicel (RC-581 FMC BioPolymer) made in complete RPMI medium. Plaques were resolved at 48 h post infection by fixing in 10 % paraformaldehyde for 15 min followed by staining for 1 hour with 0.2 % crystal violet made in 20 % ethanol. Plates were rinsed in water to visualize plaques.

#### Measurement of viral burden

Indicated organs (nasal cavity, brain, lungs from infected or uninfected mice were collected, weighed, and homogenized in 1 mL of serum free RPMI media containing penicillin-streptomycin and homogenized in 2 mL tube containing 1.5 mm Zirconium beads with BeadBug 6 homogenizer (Benchmark Scientific, TEquipment Inc). Virus titers were measured using three highly correlative methods. Frist, the total RNA was extracted from homogenized tissues using RNeasy plus Mini kit (Qiagen Cat # 74136), reverse transcribed with iScript advanced cDNA kit (Bio-Rad Cat #1725036) followed by a SYBR Green Real-time PCR assay for determining copies of SARS-CoV-2 N gene RNA using primers SARS-CoV-2 N F: 5’-ATGCTGCAATCGTGCTACAA-3’ and SARS-CoV-2 N R: 5’-GACTGCCGCCTCTGCTC-3’.

Second, serially diluted clarified tissue homogenates were used to infect Vero-E6 cell culture monolayer. The titers per gram of tissue were quantified using standard plaque forming assay described above. Third, we used nanoluciferase activity as a shorter surrogate for plaque assay. Infected cells were washed with PBS and then lysed using 1X Passive lysis buffer. The lysates transferred into a 96-well solid white plate (Costar Inc) and nanoluciferase activity was measured using Tristar multiwell Luminometer (Berthold Technology, Bad Wildbad, Germany) for 2.5 seconds by adding 20 µl of Nano-Glo® substrate in nanoluc assay buffer (Promega Inc, WI, USA). Uninfected monolayer of Vero cells treated identically served as controls to determine basal luciferase activity to obtain normalized relative light units. The data were processed and plotted using GraphPad Prism 8 v8.4.3.

#### Analyses of signature inflammatory cytokines mRNA

Brain and lung samples were collected from mice at the time of necropsy. Approximately, 20 mg of tissue was suspended in 500 µL of RLT lysis buffer, and RNA was extracted using RNeasy plus Mini kit (Qiagen Cat # 74136), reverse transcribed with iScript advanced cDNA kit (Bio-Rad Cat #1725036). To determine levels of signature inflammatory cytokines, multiplex qPCR was conducted using iQ Multiplex Powermix (Bio Rad Cat # 1725848) and PrimePCR Probe Assay mouse primers FAM-GAPDH, HEX-IL6, TEX615-CCL2, Cy5-CXCL10, and Cy5.5-IFNgamma. The reaction plate was analyzed using CFX96 touch real time PCR detection system. Scan mode was set to all channels. The PCR conditions were 95 °C 2 min, 40 cycles of 95 °C for 10 s and 60 °C for 45 s, followed by a melting curve analysis to ensure that each primer pair resulted in amplification of a single PCR product. mRNA levels of *Il6, Ccl2, Cxcl10 and Ifng* in the cDNA samples of infected mice were normalized to *Gapdh* mRNA with the formula ΔC_t_(target gene)=C_t_(target gene)-C_t_(*Gapdh*). The fold increase was determined using 2^-ΔΔCt^ method comparing treated mice to uninfected controls.

#### Antibody depletion of immune cell subsets

For evaluating the effect of NK cell depletion during CV3-1 prophylaxis, anti-NK1.1 (clone PK136; 12.5 mg/kg body weight) or an isotype control mAb (BioXCell; clone C1.18.4; 12.5 mg/kg body weight) was administered to mice by i.p. injections every 2 days starting at 48 h before SARS-CoV-2-nLuc challenge till 8 dpi. The mice were bled after two days of antibody depletion, necropsy or at 10 dpi (surviving mice) for analyses. To evaluate the effect of NK cell and neutrophil depletion during CV3-1 therapy, anti-NK1.1 (clone PK136; 12.5 mg/kg body weight) or anti-Ly6G (clone: 1A8; 12.5 mg/kg body weight) was administered to mice by i.p injection every two days starting at 1 dpi respectively. Rat IgG2a mAb (BioXCell; clone C1.18.4; 12.5 mg/kg body weight) was used as isotype control. The mice were sacrificed and bled at 10 dpi for analyses. For evaluating the effect of monocyte depletion on CV3-1 therapy, anti-CCR2 (clone MC-21; 2.5 mg/kg body weight) (Mack et al., 2001) or an isotype control mAb (BioXCell; clone LTF-2; 2.5 mg/kg body weight) was administered to mice by i.p injection every two days starting at 1 dpi. The mice were sacrificed and bled 2-3 days after antibody administration or at 10 dpi to ascertain depletion of desired population.

#### Flow Cytometric Analyses

For analysis of immune cell depletion, peripheral blood was collected before infection and on day of harvest. Erythrocytes were lysed with RBC lysis buffer (BioLegend Inc), PBMCs fixed with 4 % PFA and quenched with PBS containing 0.1M glycine. PFA-fixed cells PBMCs were resuspended and blocked in Cell Staining buffer (BioLegend Inc.) containing Fc blocking antibody against CD16/CD32 (BioLegend Inc) before staining with antibodies. NK cells were identified as CD3- NK1.1+ cells using PE/Cy7 anti-mouse CD3(17A2) and APC anti-mouse NK-1.1 (PK136). Neutrophils were identified as CD45^+^CD11b^+^Ly6G^+^ cells using APC Rat anti-mouse CD45 (30-F11), PE anti-mouse CD11b (M1/70) APC/Cy7 and anti-mouse Ly-6G (1A8). Ly6C^hi^ monocytes were identified as CD45^+^CD11b^+^Ly6C^hi^ cells using APC Rat anti-mouse CD45 (30-F11), PE anti-mouse CD11b (M1/70) and APC/Cy7 anti-mouse Ly-6C (HK1.4). Data were acquired on an Accuri C6 (BD Biosciences) and were analyzed with Accuri C6 software. FlowJo software (Treestar) was used to generate FACS plot shown in Figure S7. 100,000 – 200,000 viable cells were acquired for each sample.

#### Sample Preparation for Electron Microscopy

Lung, brain and testis tissue samples from hACE2 transgenic mice challenged intranasally with SARS-CoV-2-nLuc (1 x 10^5^ FFU; 6 dpi) were imaged after necropsy using bioluminescence imaging (IVIS, Perkin Elmer), pruned to isolate regions with high nLuc activity and immediately pre-fixed with 3 % glutaraldehyde, 1 % paraformaldehyde, 5 % sucrose in 0.1 M sodium cacodylate trihydrate to render them safe for handling outside of BSL3 containment. Viral infections of cultured cells were conducted at the UVM BSL-3 facility using an approved Institutional Biosafety protocol. SARS-CoV-2 strain 2019-nCoV/USA_USA-WA1/2020 (WA1; generously provided by K. Plante, World Reference Center for Emerging Viruses and Arboviruses, University of Texas Medical Branch) and propagated in African green monkey kidney (Vero E6) cells. Vero E6 cells were maintained in complete Dulbecco’s Modified Eagle Medium (DMEM; Thermo Fisher, Cat. #11965–092) containing 10% fetal bovine serum (Gibco, Thermo-Fisher, Cat. #16140–071), 1% HEPES Buffer Solution (15630–130), and 1% penicillin– streptomycin (Thermo Fisher, Cat. #15140–122). Cells were grown in a humidified incubator at 37 °C with 5 % CO_2_. Vero E6 cells were seeded into six well dishes and infected with SARS-CoV-2 at a multiplicity of infection of 0.01 for 48 hours before fixing and preparing for electron microscopy. Cells were pre-fixed with 3% glutaraldehyde, 1% paraformaldehyde, 5 % sucrose in 0.1M sodium cacodylate trihydrate, removed from the plates and further prepared by high-pressure freezing and freeze-substitution as described below.

Tissues samples were further cut to ∼0.5 mm^3^ blocks and cultured cells were gently pelleted. Both samples were rinsed with fresh cacodylate buffer and placed into brass planchettes (Type A; Ted Pella, Inc., Redding, CA) prefilled with 10% Ficoll in cacodylate buffer. The tissues were covered with the flat side of a Type-B brass planchette and rapidly frozen with an HPM-010 high-pressure freezing machine (Leica Microsystems, Vienna Austria). The frozen samples were transferred under liquid nitrogen to cryotubes (Nunc) containing a frozen solution of 2.5% osmium tetroxide, 0.05 % uranyl acetate in acetone. Tubes were loaded into an AFS-2 freeze-substitution machine (Leica Microsystems) and processed at -90°C for 72 h, warmed over 12 h to -20°C, held at that temperature for 6 h, then warmed to 4°C for 2 h. The fixative was removed, and the samples rinsed 4 x with cold acetone, following which they were infiltrated with Epon-Araldite resin (Electron Microscopy Sciences, Port Washington PA) over 48 h. The spleen tissue was flat-embedded between two Teflon-coated glass microscope slides. Resin was polymerized at 60°C for 48 h.

#### Electron Microscopy and Dual-Axis Tomography

Flat-embedded tissue samples or portions of cell pellets were observed with a stereo dissecting microscope and appropriate regions were extracted with a microsurgical scalpel and glued to the tips of plastic sectioning stubs. Semi-thin (150-200 nm) serial sections were cut with a UC6 ultramicrotome (Leica Microsystems) using a diamond knife (Diatome, Ltd. Switzerland). Sections were placed on formvar-coated copper-rhodium slot grids (Electron Microscopy Sciences) and stained with 3 % uranyl acetate and lead citrate. Gold beads (10 nm) were placed on both surfaces of the grid to serve as fiducial markers for subsequent image alignment. Sections were placed in a dual-axis tomography holder (Model 2040, E.A. Fischione Instruments, Export PA) and imaged with a Tecnai T12-G2 transmission electron microscope operating at 120 KeV (ThermoFisher Scientific) equipped with a 2k x 2k CCD camera (XP1000; Gatan, Inc. Pleasanton CA). Tomographic tilt-series and large-area montaged overviews were acquired automatically using the SerialEM software package (Mastronarde, 2005, 2008; Mastronarde and Held, 2017). For tomography, samples were tilted +/- 62° and images collected at 1° intervals. The grid was then rotated 90° and a similar series taken about the orthogonal axis. Tomographic data was calculated, analyzed, and modeled using the IMOD software package (Mastronarde, 2005, 2008; Mastronarde and Held, 2017) on iMac Pro and MacPro computers (Apple, Inc., Cupertino, CA). Montaged projection overviews were used to illustrate spatial perspective, identify cell types and frequency within the tissue sections. High-resolution 3D electron tomography was used to confirm virus particles and characterize virus-containing compartments within infected cells.

#### Identification and Characterization of SARS-CoV-2 Virions in infected cells and tissues

Particles resembling virions were examined in 3D by tomography to determine their identity. Presumptive SARS-CoV-2 virions were identified from tomographic reconstructions of tissue samples by observing structures resembling virions described in cryo-electron tomography studies of purified SARS-CoV-2 and of SARS-CoV-2 in infected cells (Ke et al., 2020; Klein et al., 2020; Turonova et al., 2020; Yao et al., 2020). These were compared to identified virions within SARS-CoV-2–infected cultured Vero E6 cells that had been prepared for EM by the same methodology (Figure 2O-Q). We used the following criteria to positively identify SARS-CoV-2 virions in tissues: (*i*) Structures that were spherical in 3D with ∼60-120 nM diameters and were not continuous with other adjacent structures, (*ii*) Spherical structures with densities corresponding to a distinct membrane bilayer, internal puncta consistent with ribonucleoproteins (Yao et al., 2020), and densities corresponding to surface spikes on the external peripheries of the spheres. In further characterization of virions, we noted that the inner vesicles of multivesicular bodies (MVBs) have been mis-identified as SARS-CoV-2 by electron microscopy (Calomeni et al., 2020). We therefore compared measurements of MVB inner vesicles and presumptive coronavirus virions from what we identified as intracellular exit compartments within the same tomogram (data not shown) with our previous tomographic reconstructions of MVBs (He et al., 2008; Ladinsky et al., 2012). We distinguished virions inside of cytoplasmic exit compartments from the inner vesicles of MVBs based on differences in size (MVB inner virions are generally smaller in diameter than coronaviruses) and the presence of surface spikes and internal puncta (MVB inner vesicles do not present surface spikes or internal puncta).

#### Immunoelectron microscopy

SARS-CoV-2 infected tissues were extracted and immediately fixed with 4% paraformaldehyde, 5% sucrose in 0.1M cacodylate buffer. Tissues were cut into ∼0.5 mm3 pieces and infiltrated into 2.1M sucrose in 0.1M cacodylate buffer for 24 h. Individual tissue pieces were placed onto aluminum cryosectioning stubs (Ted Pella, Inc.) and rapidly frozen in liquid nitrogen. Thin (100 nm) cryosections were cut with a UC6/FC6 cryoultramicrotome (Leica Microsystems) using a cryo-diamond knife (Diatome, Ltd., Switzerland) at -110°C. Sections were picked up with a wire loop in a drop of 2.3M sucrose in 0.1M cacodylate buffer and transferred to Formvar-coated, carbon-coated, glow-discharged 100-mesh copper/rhodium grids (Electron Microscopy Sciences). Grids were incubated 1 hr with 10% calf serum in PBS to block nonspecific antibody binding, then incubated 2 hrs with anti-S antiserum (Cohen et al., 2021). Mosaic nanoparticles elicit cross-reactive immune responses to zoonotic coronaviruses in mice (Cohen et al., 2021). diluted 1:500 in PBS with 5% calf serum. Grids were rinsed (4x 10’) with PBS then labeled for 2 hrs with 10 nm gold conjugated goat anti-mouse secondary antibody (Ted Pella, Inc.). Grids were again rinsed (4x 10’) with PBS, then 3x with distilled water and negatively stained with 1% uranyl acetate in 1% methylcellulose (Sigma) for 20’. Grids were air-dried in wire loops and imaged as described for ET.

#### Protein expression and purification

FreeStyle 293F cells (Thermo Fisher Scientific) were grown in FreeStyle 293F medium (Thermo Fisher Scientific) to a density of 1×10^6^ cells/mL at 37°C with 8% CO_2_ with regular agitation (150 rpm). Cells were transfected with a plasmid coding for recombinant stabilized SARS-CoV-2 ectodomain (S-6P; obtained from Dr. Jason S. McLellan) or SARS-CoV-2 RBD (Beaudoin-Bussieres et al., 2020) using ExpiFectamine 293 transfection reagent, as directed by the manufacturer (Thermo Fisher Scientific). One-week post-transfection, supernatants were clarified and filtered using a 0.22 µm filter (Thermo Fisher Scientific). The recombinant S-6P was purified by strep-tactin resin (IBA) following by size-exclusion chromatography on Superose 6 10/300 column (GE Healthcare) in 10 mM Tris pH 8.0 and 200 mM NaCl (SEC buffer). RBD was purified by Ni-NTA column (Invitrogen) and gel filtration on Hiload 16/600 Superdex 200pg using the same SEC buffer. Purified proteins were snap-frozen at liquid nitrogen and stored in aliquots at 80°C until further use. Protein purities were confirmed as one single-band on SDS-PAGE.

#### SARS-CoV-2 Spike ELISA (enzyme-linked immunosorbent assay)

The SARS-CoV-2 Spike ELISA assay used was recently described (Beaudoin-Bussieres et al., 2020; Prevost et al., 2020). Briefly, recombinant SARS-CoV-2 S-6P and RBD proteins (2.5 μg/ml), or bovine serum albumin (BSA) (2.5 μg/ml) as a negative control, were prepared in PBS and were adsorbed to plates (MaxiSorp; Nunc) overnight at 4 °C. Coated wells were subsequently blocked with blocking buffer (Tris-buffered saline [TBS] containing 0.1% Tween20 and 2% BSA) for 1 hour at room temperature. Wells were then washed four times with washing buffer (TBS containing 0.1% Tween20). CV3-1, CV3-25 and CR3022 mAbs (50 ng/ml) were prepared in a diluted solution of blocking buffer (0.1 % BSA) and incubated with the RBD-coated wells for 90 minutes at room temperature. Plates were washed four times with washing buffer followed by incubation with HRP-conjugated anti-IgG secondary Abs (Invitrogen) (diluted in a diluted solution of blocking buffer [0.4% BSA]) for 1 hour at room temperature, followed by four washes. HRP enzyme activity was determined after the addition of a 1:1 mix of Western Lightning oxidizing and luminol reagents (Perkin Elmer Life Sciences). Light emission was measured with a LB941 TriStar luminometer (Berthold Technologies). Signal obtained with BSA was subtracted for each plasma and was then normalized to the signal obtained with CR3022 mAb present in each plate.

#### Flow cytometry analysis of cell-surface Spike staining

Spike expressors of human coronaviruses SARS-CoV-2, SARS-CoV-1, MERS-CoV, OC43, NL63 and 229E were reported elsewhere (Hoffmann et al., 2020; Hoffmann et al., 2013; Hofmann et al., 2005; Park et al., 2016; Prevost et al., 2020). Expressors of HKU1 Spike and SARS-CoV-2 S2 N-His were purchased from Sino Biological. Using the standard calcium phosphate method, 10 μg of Spike expressor and 2 μg of a green fluorescent protein (GFP) expressor (pIRES2-eGFP) was transfected into 2 × 10^6^ 293T cells. At 48 hours post transfection, 293T cells were stained with CV3-1 and CV3-25 antibodies (5μg/mL), using cross-reactive anti-SARS-CoV-1 Spike CR3022 or mouse anti-His tag (Sigma-Aldrich) as positive controls. Alexa Fluor-647- conjugated goat anti-human IgG (H+L) Abs (Invitrogen) and goat anti-mouse IgG (H+L) Abs (Invitrogen) were used as secondary antibodies. The percentage of transfected cells (GFP+ cells) was determined by gating the living cell population based on the basis of viability dye staining (Aqua Vivid, Invitrogen). Samples were acquired on a LSRII cytometer (BD Biosciences) and data analysis was performed using FlowJo v10 (Tree Star).

#### Virus capture assay

The SARS-CoV-2 virus capture assay was previously reported (Ding et al., 2020). Briefly, pseudoviral particles were produced by transfecting 2×10^6^ HEK293T cells with pNL4.3 Luc R-E- (3.5 μg), plasmids encoding for SARS-CoV-2 Spike or SARS-CoV-1 Spike (3.5 μg) protein and VSV-G (pSVCMV-IN-VSV-G, 1 μg) using the standard calcium phosphate method. Forty-eight hours later, supernatant-containing virion was collected, and cell debris was removed through centrifugation (1,500 rpm for 10 min). To immobilize antibodies on ELISA plates, white MaxiSorp ELISA plates (Thermo Fisher Scientific) were incubated with 5 μg/ml of antibodies in 100 μl phosphate-buffered saline (PBS) overnight at 4°C. Unbound antibodies were removed by washing the plates twice with PBS. Plates were subsequently blocked with 3% bovine serum albumin (BSA) in PBS for 1 hour at room temperature. After two washes with PBS, 200 μl of virus-containing supernatant was added to the wells. After 4 to 6 hours incubation, supernatants were removed and the wells were washed with PBS 3 times. Virus capture by any given antibody was visualized by adding 1×10^4^ SARS-CoV-2-resistant Cf2Th cells per well in complete DMEM medium. Forty-eight hours post-infection, cells were lysed by the addition of 30 μL of passive lysis buffer (Promega) and three freeze-thaw cycles. An LB941 TriStar luminometer (Berthold Technologies) was used to measure the luciferase activity of each well after the addition of 100 μL of luciferin buffer (15 mM MgSO_4_, 15 mM KH_2_PO_4_ [pH 7.8], 1 mM ATP, and 1 mM dithiothreitol) and 50 μL of 1 mM D-luciferin potassium salt (ThermoFisher Scientific).

#### Surface plasmon resonance (SPR)

All surface plasma resonance assays were performed on a Biacore 3000 (GE Healthcare) with a running buffer of 10 mM HEPES pH 7.5 and 150 mM NaCl, supplemented with 0.05% Tween 20 at 25°C. The binding affinity and kinetics to the SARS-CoV-2 spike (S) trimer (SARS-CoV-2 S HexaPro [S-6P]) (Hsieh et al., 2020) and SARS-CoV-2 S2 ectodomain (baculovirus produced his-tagged S2(686-1213) from BEI Resources (NR-53799) were evaluated using monovalent CV3-1 and CV3-25 Fab. Fabs were generated by standard papain digestion (Thermo Fisher) and purified by Protein A affinity chromatography and gel filtration. His-tagged SARS-CoV-2 S-6P or SARS-CoV-2 S2 ectodomain was immobilized onto a Ni-NTA sensor chip at a level of ∼1000 and ∼630 RU response units (RUs), respectively. Two-fold serial dilutions of CV3-1 or CV3-25 Fab were injected in a concentration range of 1.56-100 nM over the SARS-CoV-2 S-6P and CV3-25 Fab in a range of 3.125 to 200 to nM over the SARS-CoV-2 S2. After each cycle the Ni-NTA sensor chip was regenerated with a wash step of 0.1 M EDTA and reloaded with 0.1 M nickel sulfate followed by the immobilization of fresh antigens for the next cycle. The binding kinetics of SARS-CoV-2 RBD and CV3-1 were obtained in a format where CV3-1 IgG was immobilized onto a Protein A sensor chip (Cytiva) with ∼300 (RUs) and serial dilutions of SARS-CoV-2 RBD were injected with concentrations ranging from 1.56 to 50 nM. The protein A chip was regenerated with a wash step of 0.1 M glycine pH 2.0 and reloaded with IgG after each cycle.

All sensograms were corrected by subtraction of the corresponding blank channel and the kinetic constant determined using a 1:1 Langmuir model with the BIA evaluation software (GE Healthcare). Goodness of fit of the curve was evaluated by the Chi^2^ value with a value below 3 considered acceptable

#### Pseudovirus neutralization assay

Target cells were infected with single-round luciferase-expressing lentiviral particles. Briefly, 293T cells were transfected by the calcium phosphate method with the pNL4.3 R-E-Luc plasmid (NIH AIDS Reagent Program) and a plasmid encoding for SARS-CoV-2 Spike at a ratio of 5:4. Two days post-transfection, cell supernatants were harvested and stored at –80°C until use. 293T-ACE2 (Prevost et al., 2020) target cells were seeded at a density of 1 × 10^4^ cells/well in 96-well luminometer-compatible tissue culture plates (Perkin Elmer) 24 h before infection. Recombinant viruses in a final volume of 100 μL were incubated with the indicated semi-log diluted antibody concentrations for 1 h at 37°C and were then added to the target cells followed by incubation for 48 h at 37°C; cells were lysed by the addition of 30 μL of passive lysis buffer (Promega) followed by one freeze-thaw cycle. An LB941 TriStar luminometer (Berthold Technologies) was used to measure the luciferase activity of each well after the addition of 100 μL of luciferin buffer (15 mM MgSO_4_, 15 mM KH_2_PO_4_ [pH 7.8], 1 mM ATP, and 1 mM dithiothreitol) and 50 μL of 1 mM d-luciferin potassium salt. The neutralization half-maximal inhibitory dilution (IC_50_) represents the plasma dilution to inhibit 50 % of the infection of 293T-ACE2 cells by recombinant viruses bearing the SARS-CoV-2 S glycoproteins.

#### Microneutralization assay

A microneutralization assay for SARS-CoV-2 serology was performed as previously described (Amanat et al., 2020). Experiments were conducted with the SARS-CoV-2 USA-WA1/2020 virus strain (obtained from BEI resources). One day prior to infection, 2×10^4^ Vero E6 cells were seeded per well of a 96 well flat bottom plate and incubated overnight at 37°C under 5% CO_2_ to permit cell adherence. Titrated antibody concentrations were performed in a separate 96 well culture plate using MEM supplemented with penicillin (100 U/mL), streptomycin (100 mg/mL), HEPES, L-Glutamine (0.3 mg/mL), 0.12% sodium bicarbonate, 2% FBS (all from Thermo Fisher Scientific) and 0.24% BSA (EMD Millipore Corporation). In a Biosafety Level 3 laboratory (ImPaKT Facility, Western University), 10^3^ TCID_50_/mL of SARS-CoV-2 USA-WA1/2020 live virus was prepared in MEM + 2% FBS and combined with an equivalent volume of respective antibody dilutions for one hour at room temperature. After this incubation, all media was removed from the 96 well plate seeded with Vero E6 cells and virus:antibody mixtures were added to each respective well at a volume corresponding to 600 TCID_50_ per well and incubated for one hour further at 37°C. Both virus only and media only (MEM + 2% FBS) conditions were included in this assay. All virus:plasma supernatants were removed from wells without disrupting the Vero E6 monolayer. Each antibody concentration (100 µL) was added to its respective Vero E6-seeded well in addition to an equivalent volume of MEM + 2% FBS and was then incubated for 48 hours. Media was then discarded and replaced with 10% formaldehyde for 24 hours to cross-link Vero E6 monolayer. Formaldehyde was removed from wells and subsequently washed with PBS. Cell monolayers were permeabilized for 15 minutes at room temperature with PBS + 0.1% Triton X-100, washed with PBS and then incubated for one hour at room temperature with PBS + 3% non-fat milk. An anti-mouse SARS-CoV-2 nucleocapsid protein (Clone 1C7, Bioss Antibodies) primary antibody solution was prepared at 1 mg/mL in PBS + 1% non-fat milk and added to all wells for one hour at room temperature. Following extensive washing with PBS, an anti-mouse IgG HRP secondary antibody solution was formulated in PBS + 1% non-fat milk. One-hour post-incubation, wells were washed with PBS, SIGMAFAST OPD developing solution (Millipore Sigma) was prepared as per manufacturer’s instructions and added to each well for 12 minutes. Dilute HCl (3.0 M) was added to quench the reaction and the optical density at 490 nm of the culture plates was immediately measured using a Synergy LX multi-mode reader and Gen5 microplate reader and imager software (BioTek).

#### SARS-CoV-2 MA10 neutralization assay

Serial two-fold dilutions of CV3-1 antibody (100 to 0.0078125 mg/mL) were prepared in triplicates in a volume of 50 μL. 30 µL of MA10 virus (1.2 x 10^4^ FFU/mL stock) was mixed with diluted antibody and incubated for 1 h at 37°C. The virus-antibody mixes were then added to Vero E6 cells (7.5 × 10^5^ cells/well) seeded 24 h earlier, in 6-well tissue culture plates and allowed to interact with cells for 1 h. The cells were then overlayed with 1 mL of pre-warmed 0.6 % Avicel (RC-581 FMC BioPolymer) made in complete RPMI medium. Plaques were resolved after 48 h by fixing cells in 10 % paraformaldehyde for 15 min followed by staining for 1 hour with 0.2 % crystal violet made in 20 % ethanol. Plates were rinsed in water to visualize FFU. The FFU counts from virus samples without antibody incubation were set to 100% (50 - 60 FFU/well). IC_50_ was calculated by plotting the log (antibody dilution) vs normalized FFUs and using non-linear fit option in GraphPad Prism.

#### Cell-to-cell fusion assay

To assess cell-to-cell fusion, 2 × 10^6^ 293T cells were co-transfected with plasmid expressing HIV-1 Tat (1µg) and a plasmid expressing SARS-CoV-2 Spike (4 µg) using the calcium phosphate method. Two days after transfection, Spike-expressing 293T (effector cells) were detached with PBS-EDTA 1mM and incubated for 1 hour with indicated amounts of CV3-1 and/or CV3-25 NAbs at 37°C and 5% CO_2_. Subsequently, effector cells (1 × 10^4^) were added to TZM-bl-ACE2 target cells that were seeded at a density of 1 × 10^4^ cells/well in 96-well luminometer-compatible tissue culture plates 24 h before the assay. Cells were co-incubated for 6 h at 37°C and 5% CO_2_, after which they were lysed by the addition of 40 μL of passive lysis buffer (Promega) and one freeze-thaw cycles. An LB 941 TriStar luminometer (Berthold Technologies) was used to measure the luciferase activity of each well after the addition of 100 μL of luciferin buffer (15 mM MgSO_4_, 15 mM KH_2_PO_4_ [pH 7.8], 1 mM ATP, and 1 mM dithiothreitol) and 50 μL of 1 mM d-luciferin potassium salt (ThermoFisher Scientific).

#### Antibody dependent cellular cytotoxicity (ADCC) assay

For evaluation of anti-SARS-CoV-2 ADCC activity, parental CEM.NKr CCR5+ cells were mixed at a 1:1 ratio with CEM.NKr-Spike cells. These cells were stained for viability (AquaVivid; Thermo Fisher Scientific) and a cellular dye (cell proliferation dye eFluor670; Thermo Fisher Scientific) and subsequently used as target cells. Overnight rested PBMCs were stained with another cellular marker (cell proliferation dye eFluor450; Thermo Fisher Scientific) and used as effector cells. Stained effector and target cells were mixed at a 10:1 ratio in 96-well V-bottom plates. Titrated concentrations of CV3-1 and CV3-25 mAbs were added to the appropriate wells. The plates were subsequently centrifuged for 1 min at 300xg, and incubated at 37°C, 5% CO2 for 5 hours before being fixed in a 2% PBS-formaldehyde solution.

ADCC activity was calculated using the formula: [(% of GFP+ cells in Targets plus Effectors)-(% of GFP+ cells in Targets plus Effectors plus antibody)]/(% of GFP+ cells in Targets) x 100 by gating on transduced live target cells. All samples were acquired on an LSRII cytometer (BD Biosciences) and data analysis performed using FlowJo v10 (Tree Star).

#### Antibody dependent cellular phagocytosis (ADCP) assay

The ADCP assay was performed using CEM.NKr-Spike cells as target cells that were fluorescently labelled with a cellular dye (cell proliferation dye eFluor450). THP-1 cells were used as effector cells and were stained with another cellular dye (cell proliferation dye eFluor670). Stained target and effector cells were mixed at a 5:1 ratio in 96-well U-bottom plates. Titrated concentrations of CV3-1 and CV3-25 mAbs were added to the appropriate wells. After an overnight incubation at 37 °C and 5% CO_2_, cells were fixed with a 2% PBS-formaldehyde solution. Antibody-mediated phagocytosis was determined by flow cytometry, gating on THP-1 cells that were double-positive for efluor450 and efluor670 cellular dyes. All samples were acquired on an LSRII cytometer (BD Biosciences) and data analysis performed using FlowJo v10 (Tree Star).

#### smFRET imaging of S on SARS-CoV-2 VLPs (S-MEN particles)

S-MEN coronavirus-like particles carrying SARS-CoV-2 spikes were prepared similarly as previously described (Lu et al., 2020). The peptides tags-carrying spike plasmid (pCMV-S Q3-1 A4-1: Q3 - GQQQLG; A4 - DSLDMLEM) was used to make S-MEN coronavirus-like particles. Plasmids encoding wildtype pCMV-S, dual-tagged pCMV-S Q3-1 A4-1, pLVX-M, pLVX-E, and pLVX-N were transfected into 293T cells at a ratio of 20:1:21:21:21. Using this very diluted ratio of tagged-S vs. wildtype S, the vast majority of S-MEN particles carry wildtype spikes. For the rest of the virus particles containing tagged S, more than 95 % S trimers will have one dual-tagged protomer and two wildtype protomers within a trimer. Using this strategy, we generated S-MEN particles with an average of one dual-tagged S protomer for conjugating FRET-paired fluorophores among predominantly wildtype S trimers presented on VLP surface. S-MEN particles were harvested 40 h post-transfection, filtered with a 0.45 µm pore size filter, and partially purified using ultra-centrifugation at 25,000 rpm for 2 h through a 15 % sucrose cushion made in PBS. S- MEN particles were then re-suspended in 50 mM pH 7.5 HEPES buffer, labeled with Cy3B(3S) and Cy5 derivative (LD650-CoA) and purified through an optiprep gradient as previously described (Lu et al., 2019; Lu et al., 2020; Munro et al., 2014) smFRET images of S-MEN particles was acquired on a home-built prism-based total internal reflection fluorescence (TIRF) microscope, as described previously (Lu et al., 2020). smFRET data analysis was performed using MATLAB (MathWorks)-based customized SPARTAN software package (Juette et al., 2016). The conformational effects of 50 ug/ml CV3-1 and CV3-25 antibodies on SARS-CoV-2 spike were tested by pre-incubating fluorescently labeled viruses for 60 mins at 37 °C before imaging in the continued presence of the antibodies. During smFRET imaging, fluorescently-labeled S-MEN particles were monitored for 80 seconds, where fluorescence from Cy3B(3S) and LD650-CoA labeled on S-MEN particles was recorded simultaneously at 25 frames per second for 80 seconds. Donor (Cy3B(3S)) and acceptor (LD650-CoA) fluorescence intensity traces were extracted after subtracting background signals and correcting cross-talks. The energy transfer efficiency (FRET) traces were generated from fluorescence intensity traces, according to FRET= I_A_/(γI_D_+I_A_), where I_D_ and I_A_ are the fluorescence intensities of donor and acceptor, respectively, γ is the correlation coefficient compromising the discrepancy in quantum yields and detection efficiencies of two fluorophores. FRET is sensitive to changes in distances between the donor and the acceptor over time, ultimately translating into the conformational profiles and dynamics of S on S-MEN particles. S-MEN particles that contain incomplete FRET-paired fluorophores or more than one FRETing pairs of donor and acceptor on a single virus particle were automatically filtered from virus pools for further analysis. FRET traces of fluorescently-labeled S-MEN particles which meet the criteria of sufficient signal-to-noise ratio and anti-correlated fluctuations in donor and acceptor fluorescence intensity are indicative of live molecules. These FRET traces, indicated the number of traces in Figure 3, were then compiled into FRET histograms in Figure 3. Each FRET histogram was fitted into the sum of four Gaussian distributions in Matlab, where each Gaussian distribution represents one conformation and the area under each Gaussian curve estimates the occupancy.

#### Quantification and Statistical Analysis

Data were analyzed and plotted using GraphPad Prism software (La Jolla, CA, USA). Statistical significance for pairwise comparisons were derived by applying non-parametric Mann-Whitney test (two-tailed). To obtain statistical significance for survival curves, grouped data were compared by log-rank (Mantel-Cox) test. To obtain statistical significance for grouped data we employed 2-way ANOVA followed by Dunnett’s or Tukey’s multiple comparison tests. p values lower than 0.05 were considered statistically significant. P values were indicated as ∗, p < 0.05; ∗∗, p < 0.01; ∗∗∗, p < 0.001; ∗∗∗∗, p < 0.0001.

#### Schematics

Schematics for showing experimental design in figures and graphical abstract were created with BioRender.com.

## Supplementary Figure Legends

**Figure S1. Widespread SARS-CoV-2 Infection in K18-hACE2 Mice. Related to Figure 1.**

(A) Images of cryosections from indicated tissues of SARS-CoV-2-nLuc infected K18-hACE2 mouse harvested at 6 dpi. Actin (green), nucleocapsid (red) and hACE2 (magenta) were detected using phalloidin and respective antibodies. Notably, hACE2 appeared as puncta on the surface of infected neurons and lung tissue compared to other organs where the signal was more uniform and stained a region of the cell surface. Scale bar: 50 µm

(B) Images of cryosections from brain tissues of SARS-CoV-2-nLuc infected K18-hACE2 mouse harvested at 6 dpi to characterize infected cells. Glial cells (top panel) were identified using antibodies to markers CD68 (magenta) and CD11b (green). Neurons (lower panel) were identified using antibodies to MAP2 (green) and mature astrocytes were identified using antibodies to GFAP (magenta). SARS-CoV-2 infected cells were identified using antibodies to nucleocapsid (red). Nucleocapsid positive cells were predominantly positive for MAP2. Scale bar: 20 µm

(C) Images of cryosections from lung and brain tissues of SARS-CoV-2-nLuc infected K18-hACE2 mouse harvested at 6 dpi to visualize neutrophil and inflammatory monocyte infiltration. Monocytes and neutrophils were identified using antibodies to markers Ly6C (magenta) and Ly6G (green) respectively. SARS-CoV-2 infected cells were identified using antibodies to nucleocapsid (red) and actin (blue) was stained using phalloidin conjugated to AF_405_. Scale bar: 20 µm

**Figure S2. Widespread Biodistribution of CV3-25 and CV3-1 NAbs in Mice 24 h After Intraperitoneal Delivery. Related to Figure 3.**

(A) C57BL/6J mice were either mock treated (PBS) or intraperitoneally administered 12.5 mg/kg body weight of CV3-25 monoclonal antibody conjugated to Alexa Fluor 594 (CV3-25 AF_594_). 24 h later, indicated tissues were imaged using the fluorescence module in IVIS spectrum to detect AF_594_. The plot shows the quantified radiance detected in indicated tissues after normalization with corresponding organs from control mouse.

(B) Images of cryosections from indicated tissues from CV3-25 AF_594_-treated mouse as in A. Actin and CV3-25 were detected using phalloidin-AF_488_ and AF_647_ conjugated anti-human IgG respectively. Scale bar: 20 µm

(C) C57BL/6J mice were either mock treated (PBS) or intraperitoneally administered 12.5 mg/kg of CV3-1 monoclonal antibody conjugated to Alexa Fluor 647 (CV3-1 AF_647_). 24 h later, indicated tissues were imaged using the fluorescence module in IVIS spectrum to detect AF_647_. The plot shows the quantified radiance detected in indicated tissues after normalization with corresponding organs from control mouse. (B) Images of cryosections from indicated tissues from CV3-1 AF_647_-treated mouse as in A. Actin was detected using phalloidin-AF_488_. CV3-1 AF_647_ was detected in the red channel using Alexa Fluor 568 conjugated anti-human IgG. Scale bar: 20 µm

**Figure S3. Assessment of CV3-1 and CV3-25 NAbs Biodistribution in Mice Using ELISA and Immunohistology. Related to Figure 4.**

(A-D) Estimation of CV3-25 and CV3-1 NAbs biodistribution in mice using ELISA. Measurement of anti-Spike NAbs levels in organs was performed using quantitative ELISA. (A-B) Recombinant SARS-CoV-2 RBD and (C-D) S-6P proteins were used to quantify CV3-1 and CV3-25 antibody levels, respectively. Linear standard curves using known concentrations of CV3-1 or CV3-25 NAbs were established for inferring the antibody concentration in organ homogenates. Serial dilutions of homogenized mice organs were prepared in PBS and incubate on antigen-coated plates. The presence of anti-Spike NAbs was revealed using HRP-conjugated anti-human IgG secondary Abs. The signal obtained with BSA (negative control) was subtracted for each organ. Relative light unit (RLU) values were transformed into a NAb concentrations based on the standard curve and the dilution factor. Subsequently, these concentration values were multiplied with the homogenization volume and divided by the total organ weight.

(E) Persistence and redistribution of neutralizing NAbs in SARS-CoV-2 infected mice. Images of brain tissue from K18-hACE2 mice infected with SARS-CoV-2-nLuc at 6 dpi that were prophylactically treated with CV3-1 or CV3-25 (12.5 mg/kg body weight), 24 h before infection. Actin (green) was labelled using phalloidin, CV3-1 and CV3-25 (magenta) were detected using anti-hIgG conjugated to AF_647_ and infected cells (red) were identified using antibodies to SARS-CoV-2 N. CV3-1 localizes to the endothelial walls of blood vessels and CV3-25 redistributes to decorate infected neurons in addition to endothelium (seen in UI mice; Figure S2). Scale bar: 50 µm

(F) Assessment of neutrophil infiltration in the brains of CV3-25 and CV3-1-pretreated mice for an experiment as in E. Neutrophils were identified using antibodies to marker Ly6G (magenta) and SARS-CoV-2 infected cells were identified using antibodies to nucleocapsid (red) and actin (blue) was stained using phalloidin conjugated to AF_405_. Scale bar: 20 µm

**Figure S4. Efficacious Dose for CV3-1 NAb During Prophylaxis. Related to Figure 4.**

(A) A scheme showing experimental design for testing the dose of CV3-1 NAb to achieve protection for lethal SARS-CoV-2-nLuc infection. Indicated concentration of CV3-1 NAb was delivered (i.p.) 1 day before challenging K18-hACE2 mice with 1 x 10^5^ FFU of SARS-CoV-2 nLuc. Human IgG1-treated (12.5 mg/kg) mice were used as control (isotype treated). Mice were followed by non-invasive BLI every 2 days from the start of infection using IVIS Spectrum after retroorbital administration of furimazine (nLuc substrate).

(B) SARS-CoV-2 replication and dissemination in K18-hACE2 transgenic mice (n = 4-6 per group) for experiment as in A, were monitored via BLI in ventral (v) and dorsal (d) positions at the indicated days post infection every 2 days. Images from two mice under CV3-1 prophylaxis (0.7 mg/kg) are shown where one mouse succumbed at 6 dpi and the other survived despite weak but observable neuroinvasion. Images from one representative experiment are shown for the rest.

(C-D) A plot showing temporal quantification of nLuc signal acquired non-invasively and displayed as photon flux (photons/sec) in whole body or brain region of SARS-CoV2-nLuc infected K18- hACE2 mice for an experiment as in A. Each curve represents luminescent signal computed for individual mouse. Scale bars denote radiance in photons per second per square centimeter per steradian (p/sec/cm2/sr).

(E) A plot showing temporal body weight changes in indicated groups of K18-hACE2 mice at indicated days post infection for an experiment shown in A. Each curve represents one animal. The body weight at the start of the experiment was set to 100 %.

(F) Kaplan-Meier survival curves of mice statistically compared by log-rank (Mantel-Cox) test for experiment as in A.

(G) Plot showing viral loads as nLuc activity/mg of indicated organs using Vero E6 cells as targets.

Undetectable virus amounts were set to 1 for display on log plots.

Grouped data in (C)-(E) and G were analyzed by 2-way ANOVA followed by Dunnett’s or Tukey’s multiple comparison tests. Statistical significance: group comparisons to isotype control are shown in black; group comparisons to CV3-1 (0.7 mg/kg) are shown in red. ∗, p < 0.05; ∗∗, p < 0.01; ∗∗∗, p < 0.001; ∗∗∗∗, p < 0.0001; Mean values ± SD are depicted.

**Figure S5. LALA Mutations Diminish and GASDALIE Mutations Enhance Antibody Effector Functions of NAbs without Compromising Neutralizing Activity. Related to Figure 6.**

(A) Cell-surface staining of CEM.NKr cells stably expressing full-length SARS-CoV-2 Spike (CEM.NKr-Spike) using CV3-1 and CV3-25 mAbs or their LALA and GASDALIE mutant counterparts. The graph shown represent the mean fluorescence intensities (MFI) obtained. with titrated concentrations of anti-Spike NAbs. MFI values obtained with parental CEM.NKr were subtracted.

(B) Pseudoviral particles encoding for the luciferase reporter gene and bearing the SARS-CoV-2 S glycoproteins were used to infect 293T-ACE2 cells. Neutralizing activity was measured by incubating pseudoviruses with titrated concentrations of anti-Spike NAbs at 37°C for 1 h prior to infection of 293T-ACE2 cells. Neutralization half maximal inhibitory antibody concentration (IC_50_) values were determined using a normalized non-linear regression using GraphPad Prism software.

(C) Using a FACS-based ADCC assay, CEM.NKr parental cells were mixed at a 1:1 ratio with CEM.NKr-Spike cells and were used as target cells. PBMCs from uninfected donors were used as effector cells. The graph shown represent the percentages of ADCC obtained in the presence of titrated concentrations of anti-Spike NAbs.

(D) Using a FACS-based ADCP assay, CEM.NKr-Spike cells were used as target cells and THP-1 monocytic cell line was used as effector cells. The graph shown represent the percentages of effector cells that had phagocytosed target cells obtained in the presence of titrated concentrations of anti-Spike NAbs. Statistical significance was tested using a non-parametric Mann-Whitney test for pairwise comparison between WT and LALA NAbs (*, p < 0.05; **, p < 0.01; ns, not significant)

(E-F) Biodistribution of LALA and GASDALIE mutants of indicated NAbs in mice 24 h after i.p. delivery. B6 mice were either isotype treated (control) or intraperitoneally administered of Alexa Fluor 647 conjugated NAb mutants (12.5 mg/kg body weight). 24 h later, indicated tissues were imaged using the fluorescence module in IVIS spectrum to detect Alexa Flour 647. The plot shows the quantified radiance detected in indicated tissues after normalization with corresponding organs from control mouse.

(G, H) Images of cryosections from brain tissues of K18-hACE2 mice pretreated with LALA mutants of CV3-25 or CV3-1 (i.p., 12.5 mg/g body weight) at 6 dpi. Actin was detected using phalloidin-Alexa Fluor 488. CV3-25 and CV3-1 (magenta) were detected using Alexa Fluor 647 conjugated anti-human IgG respectively. Infected cells were detected using antibodies to SARS-CoV-2 nucleocapsid (red). Images show penetration of both CV3-25 and CV3-1 mAbs into the brain and localization to the surface of infected neurons. Scale bar: 20 µm

(I) SARS-CoV-2 can establish infection in nasal cavity and lungs during CV3-1 prophylaxis. A scheme showing experimental design to test establishment of virus infection in K18-hACE2 mice pretreated with CV3-1 NAb (i.p.,12.5 mg/kg body weight), 1 day before challenging with 1 x 10^5^ FFU of SARS-CoV-2 nLuc. Mice treated similarly with Isotype matched hIgG1 were used as controls. The mice were followed by non-invasive BLI at 0 and 3 dpi using IVIS Spectrum after retroorbital administration of furimazine (nLuc substrate).

(J) SARS-CoV-2 replication and dissemination in indicated groups of K18-hACE2 transgenic mice (n = 5-3 per group) for experiment as in I, were monitored via BLI at the indicated times. The mice were euthanized at 3 dpi and imaged again after necropsy. Images from one representative experiment are shown.

(K) A plot showing temporal quantification of nLuc signal acquired non-invasively and displayed as photon flux (photons/sec) in whole body of SARS-CoV2-nLuc infected K18-hACE2 mice for an experiment as in I. Each line represents luminescent signal computed for individual mouse.

(L, M) *Ex vivo* imaging of indicated organs after necropsy at 3 dpi and quantification of nLuc signal displayed as photon flux (photons/sec) in K18-ACE2 mice for experiment as in I.

(N) A plot showing real-time PCR analyses to detect SARS-CoV-2 nucleocapsid (N) gene mRNA in indicated organs of K18-hACE2 mice for an experiment as in I. The data were normalized to N gene mRNA seen in uninfected mice and *Gapdh* mRNA levels. Scale bars denote radiance (photons/sec/cm^2^/steradian). *p* values obtained by non-parametric Mann-Whitney test for pairwise comparison with isotype-treated controls; *, p < 0.05; **, p < 0.01; ***, p < 0.001; ****, p < 0.0001; individual data points along with mean values ± SD are depicted.

**Figure S6. Fc-mediated Antibody Effector Functions Contribute to *In Vivo* Efficacy of CV3-1 and CV3-25 During Prophylaxis. Related to Figure 6.**

(A) A scheme showing experimental design for testing *in vivo* efficacy of NAbs and their corresponding Leucine to Alanine (LALA) or G236A/S239D/A330L/I332E (GASDALIE) mutants (12.5 mg/kg body weight) delivered intraperitoneally (i.p.) 1 day before challenging K18-hACE2 mice with 1 x 10^5^ FFU of SARS-CoV-2 nLuc. Human IgG1-treated (12.5 mg/kg body weight) mice were used as control (Iso). Mice were followed by non-invasive BLI every 2 days from the start of infection using IVIS Spectrum after retroorbital administration of furimazine (nLuc substrate).

(B) SARS-CoV-2 replication and dissemination in K18-hACE2 transgenic mice (n = 4-8 per group) for experiment as in A, were monitored via BLI in ventral (v) and dorsal (d) positions at the indicated days post infection every 2 days. The mice were euthanized on indicated days and imaged again after necropsy. Images from one representative experiment are shown.

(C-D) A plot showing temporal quantification of nLuc signal acquired non-invasively and displayed as Flux (photons/sec) in whole body or brain region of SARS-CoV2-nLuc infected K18-hACE2 mice for an experiment as in A. Each curve represents luminescent signal computed for individual mouse.

(E) A plot showing temporal body weight changes of K18-hACE2 mice at indicated days post infection for an experiment shown in A. Each curve represents one animal. The body weight at the start of the experiment was set to 100%.

(F) Kaplan-Meier survival curves of mice statistically compared by log-rank (Mantel-Cox) test for experiment as in A.

(G) Plot showing viral loads as nLuc activity/mg of indicated organs using Vero E6 cells as targets. Nluc activity was determined 24 h after infection. Undetectable virus amounts were set to 1 for display on log plots.

(H, I) A plot showing mRNA levels of indicated cytokines from lung and brain tissues of K18- hACE2 mice at the time of euthanasia as shown in F. The mRNA amounts were normalized to the levels seen in uninfected mice and the house keeping *Gapdh* mRNA.

(J) A scheme showing experimental design for testing *in vivo* efficacy of CV3-1 and its corresponding Leucine to Alanine (LALA) mutant (12.5 mg/kg body weight) delivered intraperitoneally (i.p.) 1 day before challenging 12-14 weeks old C57BL/6 (B6) mice with 5 x 10^5^ FFU of mouse-adapted SARS-CoV-2 MA10. Human IgG1-treated (12.5 mg/kg body weight) mice were used as control (Iso).

(K) A plot showing temporal body weight changes of B6 mice at indicated days post infection for an experiment shown in J. Each curve represents one animal. The body weight at the start of the experiment was set to 100%.

(L) Kaplan-Meier survival curves of mice statistically compared by log-rank (Mantel-Cox) test for experiment as in A.

(M-O) A plot showing fold changes in mRNA levels of I SARS-CoV-2 N or indicated cytokines from specified tissues of B6 mice at 7 dpi for an experiment as in J. The mRNA amounts were normalized to the levels seen in uninfected mice and the house keeping *Gapdh* mRNA.

Scale bars in (B) denote radiance (photons/sec/cm2/steradian). Grouped data in (C-E) and (G-I), were analyzed by 2-way ANOVA followed by Dunnett’s or Tukey’s multiple comparison tests. The data in (K) and (M-O) were analyzed using unpaired Mann-Whitney test. Statistical significance: group comparisons to isotype control are shown in black; group comparisons between CV3-25 LALA to CV3-25 and CV3-25 GASDALIE-treated cohorts are shown in red; group comparison between CV3-1 LALA and CV3-1 treated cohorts are shown in purple. ∗, p < 0.05; ∗∗, p < 0.01; ∗∗∗, p < 0.001; ∗∗∗∗, p < 0.0001; Mean values ± SD are depicted.

**Figure S7. NK Cells Contribute Marginally to *In Vivo* Efficacy During CV3-1 Prophylaxis. Related to Figure 7.**

(A) A scheme showing experimental design for testing the contribution of NK cells in K18-hACE2 mice pretreated with CV3-1 NAb (i.p.,12.5 mg/kg body weight), 1 day before challenging with 1 x 10^5^ FFU of SARS-CoV-2 nLuc. αNK1.1 mAb (i.p., 20 mg/kg body weight) was used to deplete NK cells at indicated time points. Corresponding human (for CV3-1) and rat (for αNK1.1) antibodies served as non-specific isotype controls. The mice were followed by non-invasive BLI every 2 days from the start of infection using IVIS Spectrum after retroorbital administration of furimazine (nLuc substrate).

(B) SARS-CoV-2 replication and dissemination in indicated groups of K18-hACE2 transgenic mice (n = 5 per group) for experiment as in A, were monitored via BLI at ventral (v) and dorsal (d) positions at the indicated days post infection every 2 days. The mice were euthanized at indicated times and imaged again after necropsy. Images from one representative experiment are shown. (C-D) A plot showing temporal quantification of nLuc signal acquired non-invasively and displayed as photon flux (photons/sec) in whole body or brain region of SARS-CoV2-nLuc infected K18- hACE2 mice for an experiment as in A. Each curve represents luminescent signal computed for individual mouse. Scale bars denote radiance (photons/sec/cm^2^/steradian).

(E) A plot showing temporal body weight changes in designated groups of K18-hACE2 mice at indicated days post infection for an experiment shown in A. Each curve represents one animal. The body weight at the start of the experiment was set to 100%.

(F) Kaplan-Meier survival curves of mice statistically compared by log-rank (Mantel-Cox) test for experiment as in A.

(G) Plot showing viral loads as nLuc activity/mg of indicated organs using Vero E6 cells as targets. Nluc activity was determined 24 h after infection. Undetectable virus amounts were set to 1 for display on log plots.

(H, I) A plot showing mRNA levels of indicated cytokines from lung and brain tissues of K18- hACE2 mice at the time of euthanasia as shown in F. The mRNA amounts were normalized to the levels seen in uninfected mice and the house keeping gene GAPDH.

(J, K) Representative FACS plots showing the gating strategy to identify NK cells (CD3^-^NK1.1^+^) and quantification to ascertain their depletion in PBMCs of indicated groups of mice.

(L, M) Representative FACS plots showing the gating strategy to identify neutrophils cells (CD45^+^ CD11b^+^Ly6G^+^) and quantification to ascertain their depletion in PBMCs of indicated groups of mice.

(N, O) Representative FACS plots showing the gating strategy to identify Ly6C^hi^ monocytes and quantification to ascertain their depletion in PBMCs of indicated groups of mice.

Grouped data in (C)-(E) and (G)-(I) were analyzed by 2-way ANOVA followed by Dunnett’s or Tukey’s multiple comparison tests. Statistical significance: group comparisons to isotype control are shown in black; group comparisons to Iso^+^CV3-1 treated cohort are in red. Pairwise comparisons in (K), (M) and (O) were analyzed using non-parametric Mann-Whitney test. ∗, p < 0.05; ∗∗, p < 0.01; ∗∗∗, p < 0.001; ∗∗∗∗, p < 0.0001; Mean values ± SD are depicted.

## Supplementary Table Legends

**Table S1**. A Comparison of SARS-CoV-2 Nucleocapsid (N) and *hACE2* mRNA Levels in K18- hACE2 Mice Across Various Organs, Related to Figure 1 and S1

*p* values obtained by non-parametric Mann-Whitney test for pairwise comparison

## Supplementary Multimedia Files

**Video S1. Longitudinal Non-invasive BLI of SARS-CoV-2-nLuc Infection and Dissemination in K18-hACE2 Mice, Related to Figure 1.**

SARS-CoV-2-nLuc challenged mice were imaged daily in dorsal (d) and ventral (v) positions for 6 days using IVIS Spectrum to monitor virus spread in the whole body as well as neuroinvasion.

**Video S2. Tomographic Reconstruction of SARS-CoV-2 Infected Lung Tissue, Related to Figure S2, panels B-D.**

Virus particles are found within membrane-enclosed exit compartments of two adjacent pulmonary capillary endothelial cells. The movie traverses the reconstructed volume to illustrate the compartments (red arrowheads) then increases in magnification to detail the virions within the compartments.

**Video S3. Cellular Overview and Tomogram of SARS-CoV-2 Infected Region in Lung Tissue, Related to Figure S2, panels B-D.**

SARS-CoV-2 virions are found in regions containing identifiable immune cells. Movie begins with a large-field montaged overview, highlighting alveolar macrophages (blue), AT2 cells (green), AT1 cells (yellow) and pulmonary blood veins (red). The upper of two blood veins is detailed at higher magnification, showing 3 red blood cells (rbc) and the surrounding capillary endothelium. A region containing portions of two endothelial cells is selected for tomographic reconstruction, showing caveolae at the cell surfaces and localizing SARS-CoV-2 virions within cytoplasmic exit compartments.

**Video S4. Tomographic Reconstruction of SARS-CoV-2 Infected Brain Tissue, Related to Figure S2, panel F.**

Virus particles are found within neurons, often appearing in linear groups within compartments bordering the edges of neuronal projections. The movie details the distinction between presumptive SARS-CoV-2 virions and typical synaptic neurotransmitter vesicles found in an adjacent synaptic terminal.

**Video S5. Tomographic Reconstruction of SARS-CoV-2 Infected Testis Tissue, Related to Figure S2, panel M.**

Virus particles are found within membrane-enclosed compartments of Sertoli cells. Additional material and structures coexist with the virions in these compartments, suggesting they may be defined as lysosomes. Presumptive SARS-CoV-2 virions can be discerned from the other structures.

## References

Alsoussi, W.B., Turner, J.S., Case, J.B., Zhao, H., Schmitz, A.J., Zhou, J.Q., Chen, R.E., Lei, T., Rizk, A.A., McIntire, K.M., et al. (2020). A Potently Neutralizing Antibody Protects Mice against SARS-CoV-2 Infection. J Immunol 205, 915–922.

Amanat, F., White, K.M., Miorin, L., Strohmeier, S., McMahon, M., Meade, P., Liu, W.C., Albrecht, R.A., Simon, V., Martinez-Sobrido, L., et al. (2020). An In Vitro Microneutralization Assay for SARS-CoV-2 Serology and Drug Screening. Curr Protoc Microbiol 58, e108.

Anand, S.P., Prevost, J., Nayrac, M., Beaudoin-Bussieres, G., Benlarbi, M., Gasser, R., Brassard, N., Laumaea, A., Gong, S.Y., Bourassa, C., et al. (2021a). Longitudinal analysis of humoral immunity against SARS-CoV-2 Spike in convalescent individuals up to 8 months post-symptom onset. bioRxiv 2021.01.25.428097.

Anand, S.P., Prevost, J., Richard, J., Perreault, J., Tremblay, T., Drouin, M., Fournier, M.J., Lewin, A., Bazin, R., and Finzi, A. (2021b). High-throughput detection of antibodies targeting the SARS-CoV-2 Spike in longitudinal convalescent plasma samples. Transfusion 10.1111/trf.16318.

Baum, A., Ajithdoss, D., Copin, R., Zhou, A., Lanza, K., Negron, N., Ni, M., Wei, Y., Mohammadi, K., Musser, B., et al. (2020). REGN-COV2 antibodies prevent and treat SARS-CoV-2 infection in rhesus macaques and hamsters. Science 370, 1110–1115.

Beaudoin-Bussieres, G., Laumaea, A., Anand, S.P., Prevost, J., Gasser, R., Goyette, G., Medjahed, H., Perreault, J., Tremblay, T., Lewin, A., et al. (2020). Decline of Humoral Responses against SARS-CoV-2 Spike in Convalescent Individuals. mBio 11, 10.1128/mBio.02590-02520.

Bolles, M., Deming, D., Long, K., Agnihothram, S., Whitmore, A., Ferris, M., Funkhouser, W., Gralinski, L., Totura, A., and Heise, M. (2011). A double-inactivated severe acute respiratory syndrome coronavirus vaccine provides incomplete protection in mice and induces increased eosinophilic proinflammatory pulmonary response upon challenge. Journal of virology 85, 12201–12215.

Bournazos, S., DiLillo, D.J., Goff, A.J., Glass, P.J., and Ravetch, J.V. (2019). Differential requirements for FcγR engagement by protective antibodies against Ebola virus. Proceedings of the National Academy of Sciences 116, 20054–20062.

Bournazos, S., Klein, F., Pietzsch, J., Seaman, M.S., Nussenzweig, M.C., and Ravetch, J.V. (2014). Broadly neutralizing anti-HIV-1 antibodies require Fc effector functions for in vivo activity. Cell 158, 1243–1253.

Calomeni, E., Satoskar, A., Ayoub, I., Brodsky, S., Rovin, B.H., and Nadasdy, T. (2020). Multivesicular bodies mimicking SARS-CoV-2 in patients without COVID-19. Kidney Int 98, 233–234.

Carossino, M., Montanaro, P., O’Connell, A., Kenney, D., Gertje, H., Grosz, K.A., Kurnick, S.A., Bosmann, M., Saeed, M., Balasuriya, U.B.R., et al. (2021). Fatal neuroinvasion of SARS-CoV-2 in K18-hACE2 mice is partially dependent on hACE2 expression. bioRxiv, 2021.2001.2013.425144.

Chen, P., Nirula, A., Heller, B., Gottlieb, R.L., Boscia, J., Morris, J., Huhn, G., Cardona, J., Mocherla, B., Stosor, V., et al. (2021). SARS-CoV-2 Neutralizing Antibody LY-CoV555 in Outpatients with Covid-19. N Engl J Med 384, 229–237.

Cohen, A.A., Gnanapragasam, P.N.P., Lee, Y.E., Hoffman, P.R., Ou, S., Kakutani, L.M., Keeffe, J.R., Wu, H.J., Howarth, M., West, A.P., et al. (2021). Mosaic nanoparticles elicit cross-reactive immune responses to zoonotic coronaviruses in mice. Science 371, 735–741.

Dinnon, K.H., 3rd, Leist, S.R., Schafer, A., Edwards, C.E., Martinez, D.R., Montgomery, S.A., West, A., Yount, B.L., Jr., Hou, Y.J., Adams, L.E., et al. (2020). A mouse-adapted model of SARS-CoV-2 to test COVID-19 countermeasures. Nature 586, 560–566.

Dekkers, G., Bentlage, A.E.H., Stegmann, T.C., Howie, H.L., Lissenberg-Thunnissen, S., Zimring, J., Rispens, T., and Vidarsson, G. (2017). Affinity of human IgG subclasses to mouse Fc gamma receptors. MAbs 9, 767–773.

Del Valle, D.M., Kim-Schulze, S., Huang, H.H., Beckmann, N.D., Nirenberg, S., Wang, B., Lavin, Y., Swartz, T.H., Madduri, D., Stock, A., et al. (2020). An inflammatory cytokine signature predicts COVID-19 severity and survival. Nat Med 26, 1636–1643.

DiLillo, D.J., Tan, G.S., Palese, P., and Ravetch, J.V. (2014). Broadly neutralizing hemagglutinin stalk–specific antibodies require FcγR interactions for protection against influenza virus in vivo. Nature medicine 20, 143–151.

Ding, S., Laumaea, A., Benlarbi, M., Beaudoin-Bussieres, G., Gasser, R., Medjahed, H., Pancera, M., Stamatatos, L., McGuire, A.T., Bazin, R., et al. (2020). Antibody Binding to SARS-CoV-2 S Glycoprotein Correlates with but Does Not Predict Neutralization. Viruses 12, 10.1101/2020.1109.1108.287482.

Ellul, M.A., Benjamin, L., Singh, B., Lant, S., Michael, B.D., Easton, A., Kneen, R., Defres, S., Sejvar, J., and Solomon, T. (2020). Neurological associations of COVID-19. Lancet Neurol 19, 767–783.

Fagre, A.C., Manhard, J., Adams, R., Eckley, M., Zhan, S., Lewis, J., Rocha, S.M., Woods, C., Kuo, K., and Liao, W. (2020). A potent SARS-CoV-2 neutralizing human monoclonal antibody that reduces viral burden and disease severity in Syrian hamsters. Frontiers in immunology 11, 10.3389/fimmu.2020.614256.

Falzarano, D., Groseth, A., and Hoenen, T. (2014). Development and application of reporter-expressing mononegaviruses: current challenges and perspectives. Antiviral Res 103, 78–87.

Finzi, A., Xiang, S.H., Pacheco, B., Wang, L., Haight, J., Kassa, A., Danek, B., Pancera, M., Kwong, P.D., and Sodroski, J. (2010). Topological layers in the HIV-1 gp120 inner domain regulate gp41 interaction and CD4-triggered conformational transitions. Mol Cell 37, 656–667.

Golden, J., Cline, C., Zeng, X., Garrison, A., Carey, B., Mucker, E., White, L., Shamblin, J., Brocato, R., and Liu, J. (2020). Human angiotensin-converting enzyme 2 transgenic mice infected with SARS-CoV-2 develop severe and fatal respiratory disease. JCI Insight 10.1172/jci.insight.142032.

Gorman, M.J., Patel, N., Guebre-Xabier, M., Zhu, A., Atyeo, C., Pullen, K.M., Loos, C., Goez-Gazi, Y., Carrion, R., Tian, J.-H., et al. (2021). Collaboration between the Fab and Fc contribute to maximal protection against SARS-CoV-2 in nonhuman primates following NVX-CoV2373 subunit vaccine with Matrix-M™ vaccination. bioRxiv, 2021.2002.2005.429759.

Graham, R.L., and Baric, R.S. (2020). SARS-CoV-2: Combating Coronavirus Emergence. Immunity 52, 734–736.

Halstead, S.B., and Katzelnick, L. (2020). COVID-19 Vaccines: Should We Fear ADE? The Journal of infectious diseases 222, 1946–1950.

Hansen, J., Baum, A., Pascal, K.E., Russo, V., Giordano, S., Wloga, E., Fulton, B.O., Yan, Y., Koon, K., and Patel, K. (2020). Studies in humanized mice and convalescent humans yield a SARS-CoV-2 antibody cocktail. Science 369, 1010–1014.

Hassan, A.O., Case, J.B., Winkler, E.S., Thackray, L.B., Kafai, N.M., Bailey, A.L., McCune, B.T., Fox, J.M., Chen, R.E., Alsoussi, W.B., et al. (2020). A SARS-CoV-2 Infection Model in Mice Demonstrates Protection by Neutralizing Antibodies. Cell 182, 744–753 e744.

He, W., Ladinsky, M.S., Huey-Tubman, K.E., Jensen, G.J., McIntosh, J.R., and Bjorkman, P.J. (2008). FcRn-mediated antibody transport across epithelial cells revealed by electron tomography. Nature 455, 542–546.

Hoffmann, M., Kleine-Weber, H., Schroeder, S., Kruger, N., Herrler, T., Erichsen, S., Schiergens, T.S., Herrler, G., Wu, N.H., Nitsche, A., et al. (2020). SARS-CoV-2 Cell Entry Depends on ACE2 and TMPRSS2 and Is Blocked by a Clinically Proven Protease Inhibitor. Cell 181, 271–280 e278.

Hoffmann, M., Muller, M.A., Drexler, J.F., Glende, J., Erdt, M., Gutzkow, T., Losemann, C., Binger, T., Deng, H., Schwegmann-Wessels, C., et al. (2013). Differential sensitivity of bat cells to infection by enveloped RNA viruses: coronaviruses, paramyxoviruses, filoviruses, and influenza viruses. PLoS One 8, e72942.

Hofmann, H., Pyrc, K., van der Hoek, L., Geier, M., Berkhout, B., and Pohlmann, S. (2005). Human coronavirus NL63 employs the severe acute respiratory syndrome coronavirus receptor for cellular entry. Proc Natl Acad Sci U S A 102, 7988–7993.

Hsieh, C.L., Goldsmith, J.A., Schaub, J.M., DiVenere, A.M., Kuo, H.C., Javanmardi, K., Le, K.C., Wrapp, D., Lee, A.G., Liu, Y., et al. (2020). Structure-based design of prefusion-stabilized SARS-CoV-2 spikes. Science 369, 1501–1505.

Iwasaki, A., and Yang, Y. (2020). The potential danger of suboptimal antibody responses in COVID-19. Nat Rev Immunol 20, 339–341.

Jennewein, M.F., MacCamy, A.J., Akins, N.R., Feng, J., Homad, L.J., Hurlburt, N.K., Seydoux, E., Wan, Y.H., Stuart, A.B., Edara, V.V., et al. (2021). Isolation and Characterization of Cross-Neutralizing Coronavirus Antibodies from COVID-19+ Subjects. bioRxiv 2021, 10.1101/2021.03.23.436684

Johansen, M.D., Irving, A., Montagutelli, X., Tate, M.D., Rudloff, I., Nold, M.F., Hansbro, N.G., Kim, R.Y., Donovan, C., Liu, G., et al. (2020). Animal and translational models of SARS-CoV-2 infection and COVID-19. Mucosal Immunol 13, 877–891.

Juette, M.F., Terry, D.S., Wasserman, M.R., Altman, R.B., Zhou, Z., Zhao, H., and Blanchard, S.C. (2016). Single-molecule imaging of non-equilibrium molecular ensembles on the millisecond timescale. Nat Methods 13, 341–344.

Ke, Z., Oton, J., Qu, K., Cortese, M., Zila, V., McKeane, L., Nakane, T., Zivanov, J., Neufeldt, C.J., Cerikan, B., et al. (2020). Structures and distributions of SARS-CoV-2 spike proteins on intact virions. Nature 588, 498–502.

Klasse, P.J., and Moore, J.P. (2020). Antibodies to SARS-CoV-2 and their potential for therapeutic passive immunization. Elife 9, 10.7554/eLife.57877.

Klein, S., Cortese, M., Winter, S.L., Wachsmuth-Melm, M., Neufeldt, C.J., Cerikan, B., Stanifer, M.L., Boulant, S., Bartenschlager, R., and Chlanda, P. (2020). SARS-CoV-2 structure and replication characterized by in situ cryo-electron tomography. Nat Commun 11, 5885.

Ladinsky, M.S., Huey-Tubman, K.E., and Bjorkman, P.J. (2012). Electron tomography of late stages of FcRn-mediated antibody transcytosis in neonatal rat small intestine. Mol Biol Cell 23, 2537–2545.

Leist, S.R., Dinnon, K.H., 3rd, Schafer, A., Tse, L.V., Okuda, K., Hou, Y.J., West, A., Edwards, C.E., Sanders, W., Fritch, E.J., et al. (2020a). A Mouse-Adapted SARS-CoV-2 Induces Acute Lung Injury and Mortality in Standard Laboratory Mice. Cell 183, 1070–1085 e1012.

Leist, S.R., Schäfer, A., and Martinez, D.R. (2020b). Cell and animal models of SARS-CoV-2 pathogenesis and immunity. Disease Models & Mechanisms 13, dmm046581.

Li, D., Edwards, R.J., Manne, K., Martinez, D.R., Schäfer, A., Alam, S.M., Wiehe, K., Lu, X., Parks, R., Sutherland, L.L., et al. (2021). The functions of SARS-CoV-2 neutralizing and infection-enhancing antibodies in vitro and in mice and nonhuman primates. bioRxiv, 2020.2012.2031.424729.

Li, W., Chen, C., Drelich, A., Martinez, D.R., Gralinski, L.E., Sun, Z., Schafer, A., Kulkarni, S.S., Liu, X., Leist, S.R., et al. (2020). Rapid identification of a human antibody with high prophylactic and therapeutic efficacy in three animal models of SARS-CoV-2 infection. Proc Natl Acad Sci U S A 117, 29832–29838.

Liu, L., Wang, P., Nair, M.S., Yu, J., Rapp, M., Wang, Q., Luo, Y., Chan, J.F., Sahi, V., Figueroa, A., et al. (2020). Potent neutralizing antibodies against multiple epitopes on SARS-CoV-2 spike. Nature 584, 450–456.

Liu, Z., Pan, Q., Ding, S., Qian, J., Xu, F., Zhou, J., Cen, S., Guo, F., and Liang, C. (2013). The interferon-inducible MxB protein inhibits HIV-1 infection. Cell Host Microbe 14, 398–410.

Lu, L.L., Suscovich, T.J., Fortune, S.M., and Alter, G. (2018). Beyond binding: antibody effector functions in infectious diseases. Nature Reviews Immunology 18, 46.

Lu, M., Ma, X., Castillo-Menendez, L.R., Gorman, J., Alsahafi, N., Ermel, U., Terry, D.S., Chambers, M., Peng, D., Zhang, B., et al. (2019). Associating HIV-1 envelope glycoprotein structures with states on the virus observed by smFRET. Nature 568, 415–419.

Lu, M., Uchil, P.D., Li, W., Zheng, D., Terry, D.S., Gorman, J., Shi, W., Zhang, B., Zhou, T., Ding, S., et al. (2020). Real-Time Conformational Dynamics of SARS-CoV-2 Spikes on Virus Particles. Cell Host Microbe 28, 880–891 e888.

Mack, M., Cihak, J., Simonis, C., Luckow, B., Proudfoot, A.E., Plachý, J.í., Brühl, H., Frink, M., Anders, H.-J., and Vielhauer, V. (2001). Expression and characterization of the chemokine receptors CCR2 and CCR5 in mice. The Journal of Immunology 166, 4697–4704.

Mastronarde, D.N. (2005). Automated electron microscope tomography using robust prediction of specimen movements. J Struct Biol 152, 36–51.

Mastronarde, D.N. (2008). Correction for non-perpendicularity of beam and tilt axis in tomographic reconstructions with the IMOD package. J Microsc 230, 212–217.

Mastronarde, D.N., and Held, S.R. (2017). Automated tilt series alignment and tomographic reconstruction in IMOD. J Struct Biol 197, 102–113.

McCray, P.B., Pewe, L., Wohlford-Lenane, C., Hickey, M., Manzel, L., Shi, L., Netland, J., Jia, H.P., Halabi, C., and Sigmund, C.D. (2007). Lethal infection of K18-hACE2 mice infected with severe acute respiratory syndrome coronavirus. Journal of virology 81, 813–821.

Munro, J.B., Gorman, J., Ma, X., Zhou, Z., Arthos, J., Burton, D.R., Koff, W.C., Courter, J.R., Smith, A.B., 3rd, Kwong, P.D., et al. (2014). Conformational dynamics of single HIV-1 envelope trimers on the surface of native virions. Science 346, 759–763.

Noy-Porat, T., Mechaly, A., Levy, Y., Makdasi, E., Alcalay, R., Gur, D., Aftalion, M., Falach, R., Ben-Arye, S.L., Lazar, S., et al. (2021). Therapeutic antibodies, targeting the SARS-CoV-2 spike N-terminal domain, protect lethally infected K18-hACE2 mice. bioRxiv, 2021.2002.2002.428995.

Park, J.E., Li, K., Barlan, A., Fehr, A.R., Perlman, S., McCray, P.B., Jr., and Gallagher, T. (2016). Proteolytic processing of Middle East respiratory syndrome coronavirus spikes expands virus tropism. Proc Natl Acad Sci U S A 113, 12262–12267.

Prevost, J., Gasser, R., Beaudoin-Bussieres, G., Richard, J., Duerr, R., Laumaea, A., Anand, S.P., Goyette, G., Benlarbi, M., Ding, S., et al. (2020). Cross-Sectional Evaluation of Humoral Responses against SARS-CoV-2 Spike. Cell Rep Med 1, 100126.

Rogers, T.F., Zhao, F., Huang, D., Beutler, N., Burns, A., He, W.T., Limbo, O., Smith, C., Song, G., Woehl, J., et al. (2020). Isolation of potent SARS-CoV-2 neutralizing antibodies and protection from disease in a small animal model. Science 369, 956–963.

Ruckwardt, T.J., Morabito, K.M., and Graham, B.S. (2019). Immunological lessons from respiratory syncytial virus vaccine development. Immunity 51, 429–442.

Saunders, K.O. (2019). Conceptual Approaches to Modulating Antibody Effector Functions and Circulation Half-Life. Front Immunol 10, 1296.

Sauer, M.M., Tortorici, M.A., Park, Y.J., Walls, A.C., Homad, L., Acton, O.J., Bowen, J.E., Wang, C., Xiong, X., de van der Schueren, W., et al. (2021). Structural basis for broad coronavirus neutralization. Nat Struct Mol Biol 28, 478–486.

Schafer, A., Muecksch, F., Lorenzi, J.C.C., Leist, S.R., Cipolla, M., Bournazos, S., Schmidt, F., Maison, R.M., Gazumyan, A., Martinez, D.R., et al. (2021). Antibody potency, effector function, and combinations in protection and therapy for SARS-CoV-2 infection in vivo. J Exp Med 218, 10.1084/jem.20201993.

Shi, P.Y., Plante, J., Liu, Y., Liu, J., Xia, H., Johnson, B., Lokugamage, K., Zhang, X., Muruato, A., Zou, J., et al. (2020a). Spike mutation D614G alters SARS-CoV-2 fitness and neutralization susceptibility. Res Sq, 10.21203/rs.21203.rs-70482/v21201.

Shi, R., Shan, C., Duan, X., Chen, Z., Liu, P., Song, J., Song, T., Bi, X., Han, C., and Wu, L. (2020b). A human neutralizing antibody targets the receptor-binding site of SARS-CoV-2. Nature 584, 120–124.

Silvas, J., Morales-Vasquez, D., Park, J.-G., Chiem, K., Torrelles, J.B., Platt, R.N., Anderson, T., Ye, C., and Martinez-Sobrido, L. (2021). Contribution of SARS-CoV-2 accessory proteins to viral pathogenicity in K18 hACE2 transgenic mice. bioRxiv, 2021.2003.2009.434696.

Smith, P., DiLillo, D.J., Bournazos, S., Li, F., and Ravetch, J.V. (2012). Mouse model recapitulating human Fcγ receptor structural and functional diversity. Proceedings of the National Academy of Sciences 109, 6181–6186.

Stamatatos, L., Czartoski, J., Wan, Y.-H., Homad, L.J., Rubin, V., Glantz, H., Neradilek, M., Seydoux, E., Jennewein, M.F., MacCamy, A.J., et al. (2021). Antibodies elicited by SARS-CoV-2 infection and boosted by vaccination neutralize an emerging variant and SARS-CoV-1. medRxiv, 2021.2002.2005.21251182.

ter Meulen, J., van den Brink, E.N., Poon, L.L., Marissen, W.E., Leung, C.S., Cox, F., Cheung, C.Y., Bakker, A.Q., Bogaards, J.A., van Deventer, E., et al. (2006). Human monoclonal antibody combination against SARS coronavirus: synergy and coverage of escape mutants. PLoS Med 3, e237.

Tortorici, M.A., Beltramello, M., Lempp, F.A., Pinto, D., Dang, H.V., Rosen, L.E., McCallum, M., Bowen, J., Minola, A., Jaconi, S., et al. (2020). Ultrapotent human antibodies protect against SARS-CoV-2 challenge via multiple mechanisms. Science 370, 950–957.

Turonova, B., Sikora, M., Schurmann, C., Hagen, W.J.H., Welsch, S., Blanc, F.E.C., von Bulow, S., Gecht, M., Bagola, K., Horner, C., et al. (2020). In situ structural analysis of SARS-CoV-2 spike reveals flexibility mediated by three hinges. Science 370, 203–208.

Ventura, J.D., Beloor, J., Allen, E., Zhang, T., Haugh, K.A., Uchil, P.D., Ochsenbauer, C., Kieffer, C., Kumar, P., Hope, T.J., et al. (2019). Longitudinal bioluminescent imaging of HIV-1 infection during antiretroviral therapy and treatment interruption in humanized mice. PLoS Pathog 15, e1008161.

Voss, W.N., Hou, Y.J., Johnson, N.V., Kim, J.E., Delidakis, G., Horton, A.P., Bartzoka, F., Paresi, C.J., Tanno, Y., Abbasi, S.A., et al. (2020). Prevalent, protective, and convergent IgG recognition of SARS-CoV-2 non-RBD spike epitopes in COVID-19 convalescent plasma. bioRxiv, 10.1101/2020.1112.1120.423708.

Weinreich, D.M., Sivapalasingam, S., Norton, T., Ali, S., Gao, H., Bhore, R., Musser, B.J., Soo, Y., Rofail, D., Im, J., et al. (2021). REGN-COV2, a Neutralizing Antibody Cocktail, in Outpatients with Covid-19. N Engl J Med 384, 238–251.

Winkler, E.S., Bailey, A.L., Kafai, N.M., Nair, S., McCune, B.T., Yu, J., Fox, J.M., Chen, R.E., Earnest, J.T., and Keeler, S.P. (2020). SARS-CoV-2 infection of human ACE2-transgenic mice causes severe lung inflammation and impaired function. Nature immunology 21, 1327–1335.

Winkler, E.S., Gilchuk, P., Yu, J., Bailey, A.L., Chen, R.E., Chong, Z., Zost, S.J., Jang, H., Huang, Y., Allen, J.D., et al. (2021). Human neutralizing antibodies against SARS-CoV-2 require intact Fc effector functions for optimal therapeutic protection. Cell, 10.1016/j.cell.2021.1002.1026.

Xie, X., Muruato, A., Lokugamage, K.G., Narayanan, K., Zhang, X., Zou, J., Liu, J., Schindewolf, C., Bopp, N.E., Aguilar, P.V., et al. (2020a). An Infectious cDNA Clone of SARS-CoV-2. Cell Host Microbe 27, 841–848 e843.

Xie, X., Muruato, A.E., Zhang, X., Lokugamage, K.G., Fontes-Garfias, C.R., Zou, J., Liu, J., Ren, P., Balakrishnan, M., Cihlar, T., et al. (2020b). A nanoluciferase SARS-CoV-2 for rapid neutralization testing and screening of anti-infective drugs for COVID-19. Nat Commun 11, 5214.

Yao, H., Song, Y., Chen, Y., Wu, N., Xu, J., Sun, C., Zhang, J., Weng, T., Zhang, Z., Wu, Z., et al. (2020). Molecular Architecture of the SARS-CoV-2 Virus. Cell 183, 730–738 e713.

Zhang, L., Jackson, C.B., Mou, H., Ojha, A., Peng, H., Quinlan, B.D., Rangarajan, E.S., Pan, A., Vanderheiden, A., Suthar, M.S., et al. (2020). SARS-CoV-2 spike-protein D614G mutation increases virion spike density and infectivity. Nat Commun 11, 10.1038/s41467-41020-19808-41464.

Zost, S.J., Gilchuk, P., Case, J.B., Binshtein, E., Chen, R.E., Nkolola, J.P., Schäfer, A., Reidy, J.X., Trivette, A., and Nargi, R.S. (2020a). Potently neutralizing and protective human antibodies against SARS-CoV-2. Nature 584, 443–449.

Zost, S.J., Gilchuk, P., Chen, R.E., Case, J.B., Reidy, J.X., Trivette, A., Nargi, R.S., Sutton, R.E., Suryadevara, N., and Chen, E.C. (2020b). Rapid isolation and profiling of a diverse panel of human monoclonal antibodies targeting the SARS-CoV-2 spike protein. Nature medicine 26, 1422–1427.

