## Supplementary table 1_Fig_1-7 for "Live Imaging of SARS-CoV-2 Infection in Mice Reveals Neutralizing Antibodies Require Fc Function for Optimal Efficacy"

**Table S1**

| Organs | Fold increase in GAPDH-normalized mRNA levels in tissues compared to uninfected B6 mice | |
|---|---|---|
| | **SARS-CoV-2 N (Mean ± SD) (n=6) (Infected K18-hACE2 mice, 6 dpi) (*p value*)** | **hACE2 (Mean ± SD) (n= 4) (UI K18-hACE2 mice) (*p value*)** |
| Brain | 303476.00 (± 27526) *(0.0022)* | 353.62 (± 145) *(0.0022)* |
| Lung | 2383.07 (± 2666) *(0.0022)* | 1131.08 (± 415) *(0.0022)* |
| Nose | 6081.23 (± 5577) *(0.0022)* | 518.60 (± 390) *(0.0022)* |
| cLNs | 541.27 (± 541) *(0.0022)* | 688.96 (± 379) *(0.0022)* |
| Trachea | 218.11 (± 206) *(0.0022)* | 375.90 (± 315) *(0.0022)* |
| Heart | 29.50 (± 17) *(0.0022)* | 159.56 (± 55) *(0.0022)* |
| Liver | 2.82 (± 1.5) *(0.0087)* | 49.46 (± 12) (*0.0022)* |
| Spleen | 11.65 (± 9) *(0.0087)* | 19.75 (± 8) *(0.0022)* |
| Kidney | 43.33 (± 21) *(0.0022)* | 487.90 (± 254) *(0.0022)* |
| Gut | 30.85 (± 20) *(0.0022)* | 320.03 (± 110) *(0.0022)* |
| Genital tract | 936.94 (± 364) *(0.0022)* | 724.26 (± 275) *(0.0022)* |

Figure S1

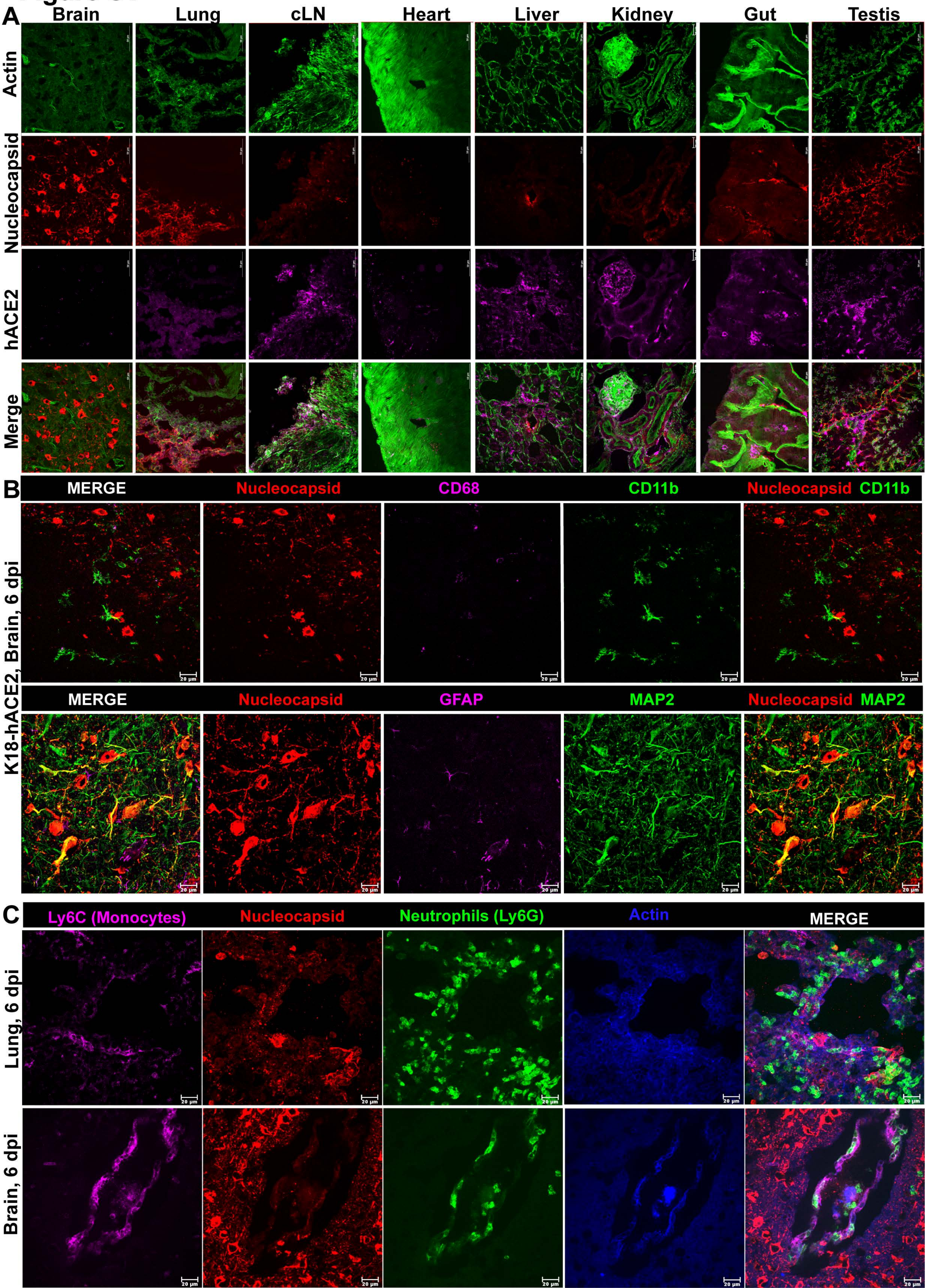
A
Brain
Lung
cLN
Heart
Liver
Kidney
Gut
Testis
Actin
Nucleocapsid
hACE2
Merge
B
MERGE
Nucleocapsid
CD68
CD11b
Nucleocapsid CD11b
K18-hACE2, Brain, 6 dpi
MERGE
Nucleocapsid
GFAP
MAP2
Nucleocapsid MAP2
20 µm
C
Ly6C (Monocytes)
Nucleocapsid
Neutrophils (Ly6G)
Actin
MERGE
Lung, 6 dpi
Brain, 6 dpi

### Figure S2

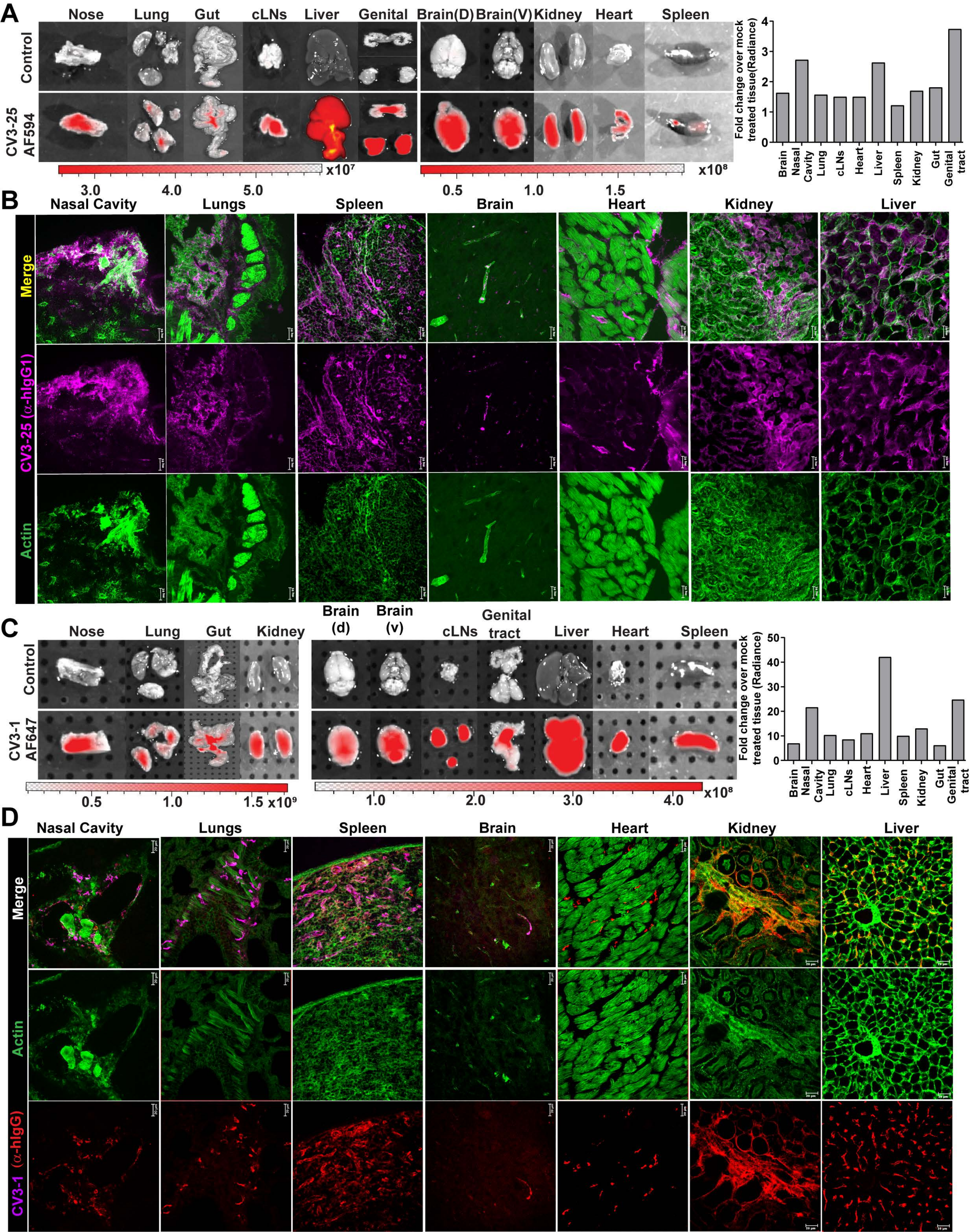
A
Nose
Lung
Gut
cLNs
Liver
Genital
Brain(D)
Brain(V)
Kidney
Heart
Spleen
Control
CV3-25 AF594
x10⁷
3.0
4.0
5.0
x10⁸
0.5
1.0
1.5
Fold change over mock treated tissue(Radiance)
0
1
2
3
4
Brain
Nasal Cavity
Lung
cLNs
Heart
Liver
Spleen
Kidney
Gut
Genital tract
B
Nasal Cavity
Lungs
Spleen
Brain
Heart
Kidney
Liver
Merge
CV3-25 (α-hIgG1)
Actin
C
Nose
Lung
Gut
Kidney
Brain (d)
Brain (v)
cLNs
Genital tract
Liver
Heart
Spleen
Control
CV3-1 AF647
0.5
1.0
1.5 x10⁹
1.0
2.0
3.0
4.0
x10⁸
Fold change over mock treated tissue (Radiance)
0
10
20
30
40
50
Brain
Nasal Cavity
Lung
cLNs
Heart
Liver
Spleen
Kidney
Gut
Genital tract
D
Nasal Cavity
Lungs
Spleen
Brain
Heart
Kidney
Liver
Merge
Actin
CV3-1 (α-hIgG)

### Figure S3

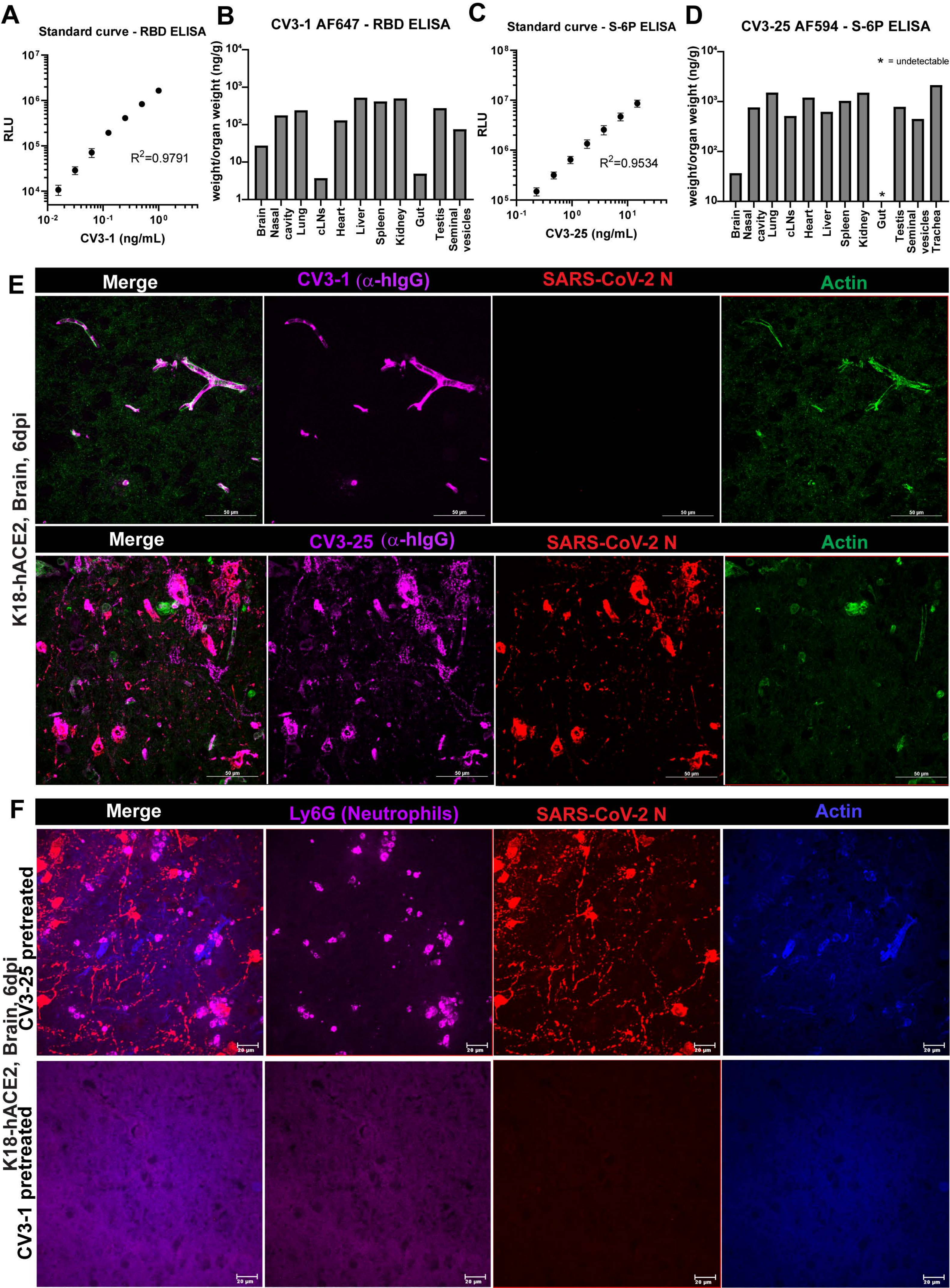
A
Standard curve - RBD ELISA
RLU
CV3-1 (ng/mL)
R2=0.9791
B
CV3-1 AF647 - RBD ELISA
weight/organ weight (ng/g)
Brain
Nasal cavity
Lung
cLNs
Heart
Liver
Spleen
Kidney
Gut
Testis
Seminal vesicles
C
Standard curve - S-6P ELISA
RLU
CV3-25 (ng/mL)
R2=0.9534
D
CV3-25 AF594 - S-6P ELISA
* = undetectable
weight/organ weight (ng/g)
Brain
Nasal cavity
Lung
cLNs
Heart
Liver
Spleen
Kidney
Gut
Testis
Seminal vesicles
Trachea
E
Merge
CV3-1 (α-hIgG)
SARS-CoV-2 N
Actin
K18-hACE2, Brain, 6dpi
Merge
CV3-25 (α-hIgG)
SARS-CoV-2 N
Actin
50 µm
F
Merge
Ly6G (Neutrophils)
SARS-CoV-2 N
Actin
K18-hACE2, Brain, 6dpi
CV3-25 pretreated
CV3-1 pretreated
20 µm

### Figure S4

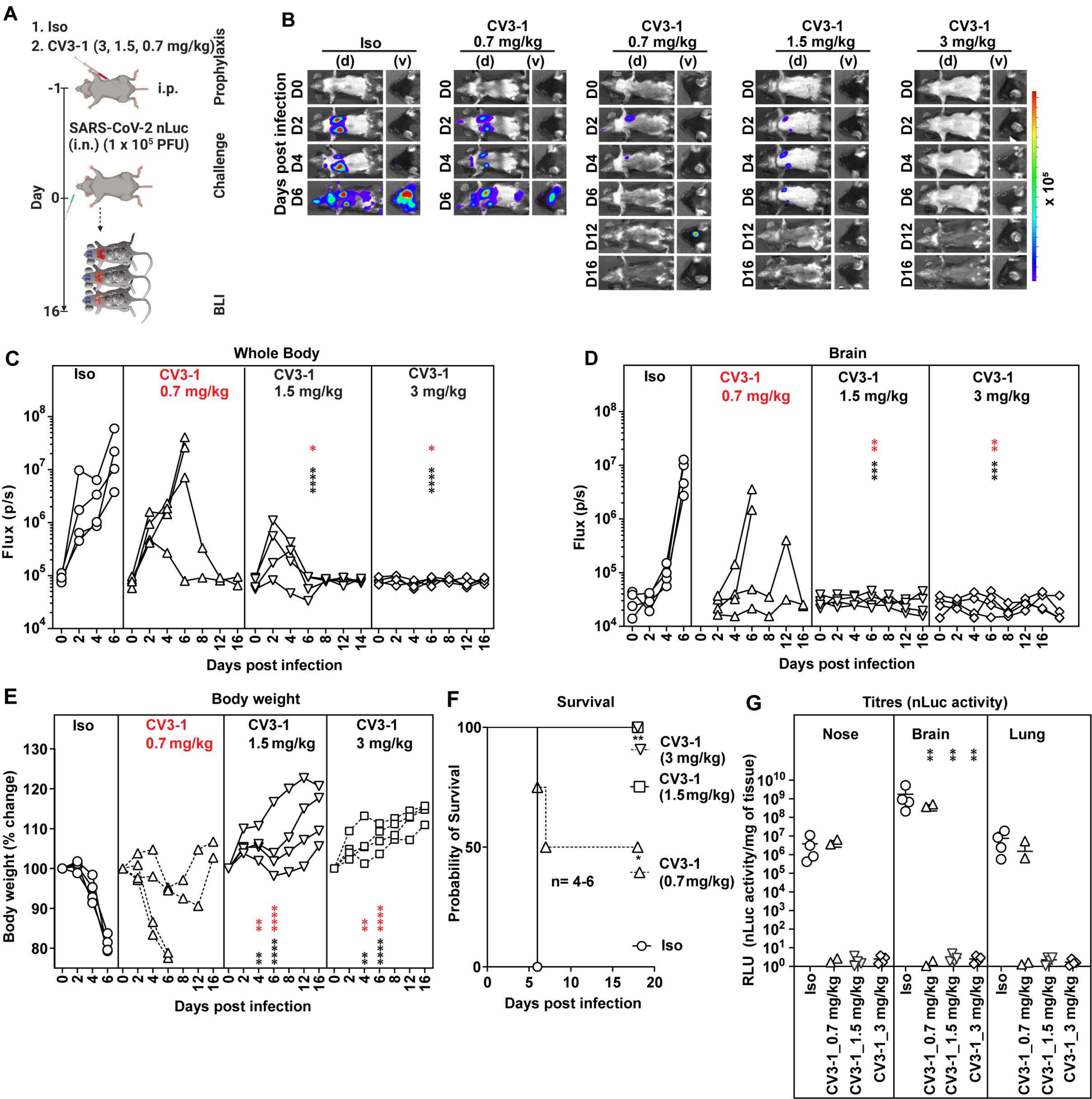
A
1. Iso
2. CV3-1 (3, 1.5, 0.7 mg/kg)
i.p.
Prophylaxis
SARS-CoV-2 nLuc (i.n.) (1 x 10^5 PFU)
Challenge
Day
BLI
B
Iso
CV3-1 0.7 mg/kg
CV3-1 1.5 mg/kg
CV3-1 3 mg/kg
(d) (v)
Days post infection
D0 D2 D4 D6 D12 D16
x 10^5
C
Whole Body
Flux (p/s)
Days post infection
D
Brain
E
Body weight
Body weight (% change)
F
Survival
Probability of Survival
CV3-1 (3 mg/kg)
CV3-1 (1.5mg/kg)
CV3-1 (0.7mg/kg)
Iso
n= 4-6
G
Titres (nLuc activity)
Nose
Brain
Lung
RLU (nLuc activity/mg of tissue)
CV3-1_0.7 mg/kg
CV3-1_1.5 mg/kg
CV3-1_3 mg/kg

### Figure S5

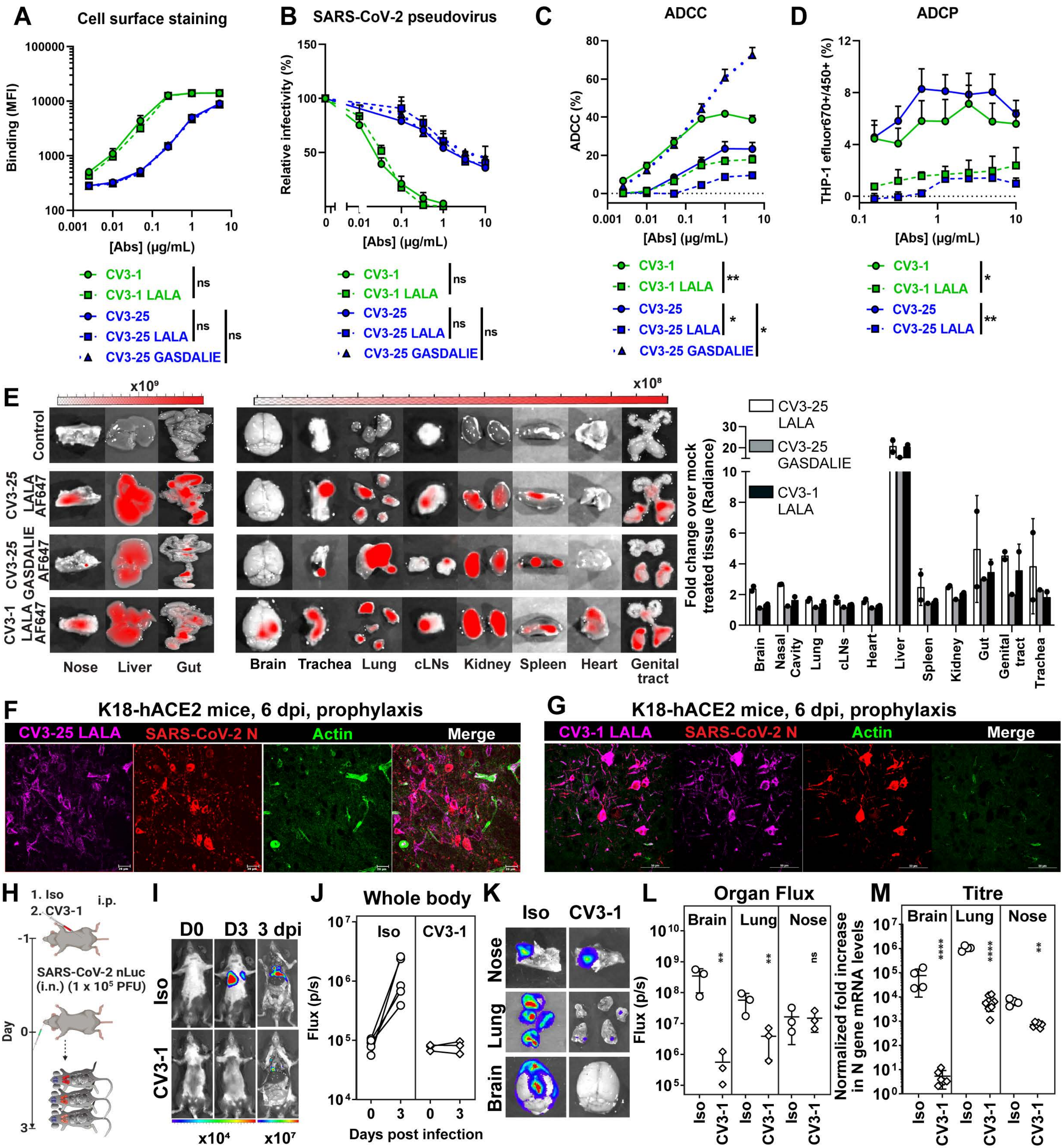
A
Cell surface staining
Binding (MFI)
[Abs] (μg/mL)
CV3-1
CV3-1 LALA
CV3-25
CV3-25 LALA
CV3-25 GASDALIE
ns
B
SARS-CoV-2 pseudovirus
Relative infectivity (%)
C
ADCC
ADCC (%)
D
ADCP
THP-1 efluor670+/450+ (%)
E
x10⁹
x10⁸
Control
CV3-25 LALA AF647
CV3-25 GASDALIE AF647
CV3-1 LALA AF647
Nose
Liver
Gut
Brain
Trachea
Lung
cLNs
Kidney
Spleen
Heart
Genital tract
Fold change over mock treated tissue (Radiance)
Nasal Cavity
F
K18-hACE2 mice, 6 dpi, prophylaxis
CV3-25 LALA
SARS-CoV-2 N
Actin
Merge
G
CV3-1 LALA
H
1. Iso
2. CV3-1
i.p.
SARS-CoV-2 nLuc (i.n.) (1 x 10⁵ PFU)
Day
I
D0
D3
3 dpi
Iso
CV3-1
x10⁴
x10⁷
J
Whole body
Flux (p/s)
Days post infection
K
L
Organ Flux
M
Titre
Normalized fold increase in N gene mRNA levels

### Figure S6

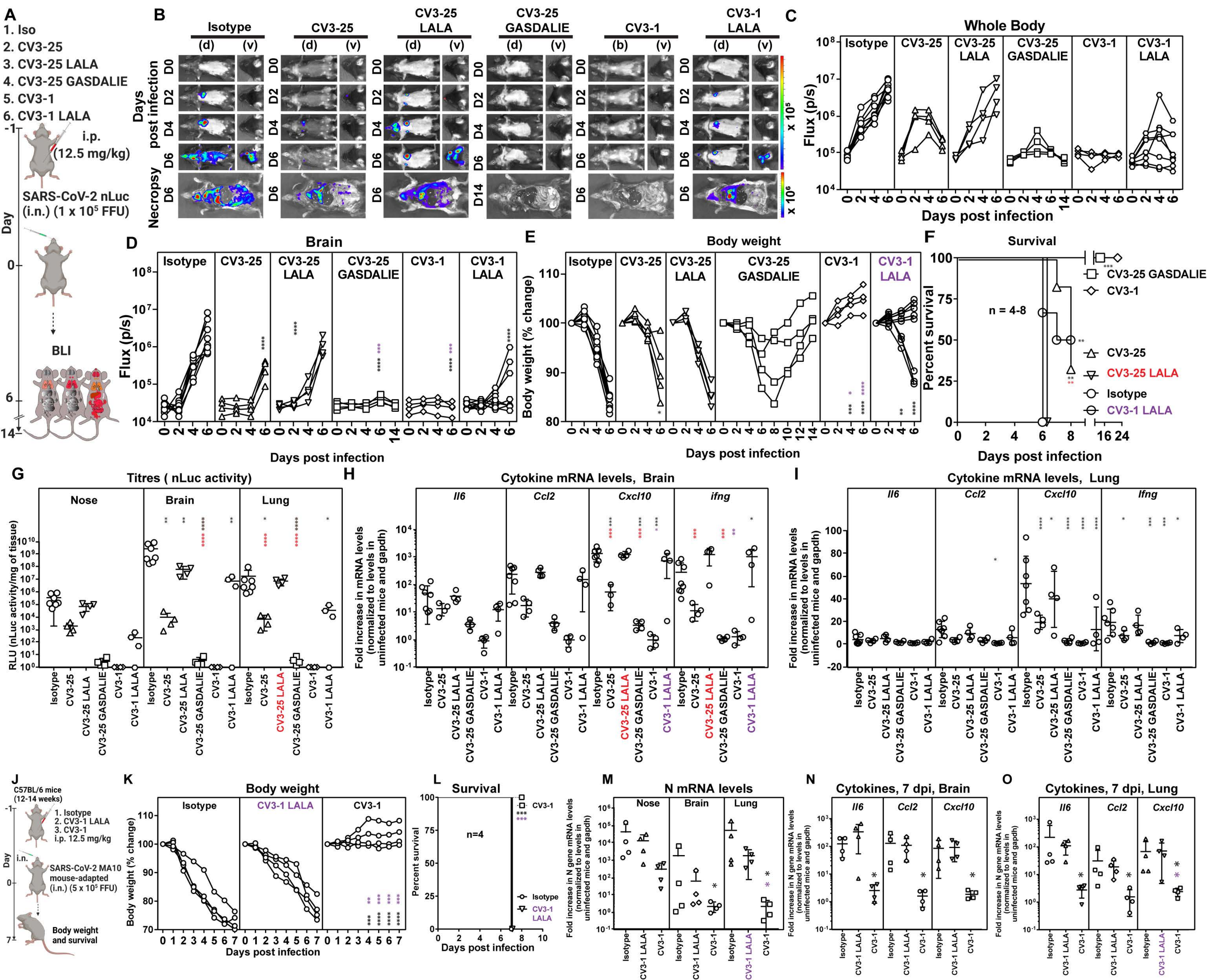

A
1. Iso
2. CV3-25
3. CV3-25 LALA
4. CV3-25 GASDALIE
5. CV3-1
6. CV3-1 LALA
i.p. (12.5 mg/kg)
SARS-CoV-2 nLuc (i.n.) (1 x 10^5 FFU)
BLI
Day
B
Isotype
CV3-25
CV3-25 LALA
CV3-25 GASDALIE
CV3-1
CV3-1 LALA
Days post infection
Necropsy
C
Whole Body
Flux (p/s)
Days post infection
D
Brain
Flux (p/s)
Days post infection
E
Body weight
Body weight (% change)
Days post infection
F
Survival
Percent survival
n = 4-8
CV3-25 GASDALIE
CV3-1
CV3-25
CV3-25 LALA
Isotype
CV3-1 LALA
Days post infection
G
Titres ( nLuc activity)
Nose
Brain
Lung
RLU (nLuc activity/mg of tissue)
H
Cytokine mRNA levels, Brain
Il6
Ccl2
Cxcl10
ifng
Fold increase in mRNA levels (normalized to levels in uninfected mice and gapdh)
I
Cytokine mRNA levels, Lung
Il6
Ccl2
Cxcl10
Ifng
J
C57BL/6 mice (12-14 weeks)
1. Isotype
2. CV3-1 LALA
3. CV3-1
i.p. 12.5 mg/kg
i.n.
SARS-CoV-2 MA10 mouse-adapted (i.n.) (5 x 10^5 FFU)
Body weight and survival
K
Body weight
Isotype
CV3-1 LALA
CV3-1
Body weight (% change)
Days post infection
L
Survival
n=4
CV3-1
Isotype
CV3-1 LALA
Percent survival
Days post infection
M
N mRNA levels
Nose
Brain
Lung
Fold increase in N gene mRNA levels (normalized to levels in uninfected mice and gapdh)
N
Cytokines, 7 dpi, Brain
Il6
Ccl2
Cxcl10
O
Cytokines, 7 dpi, Lung
Il6
Ccl2
Cxcl10

### Figure S7

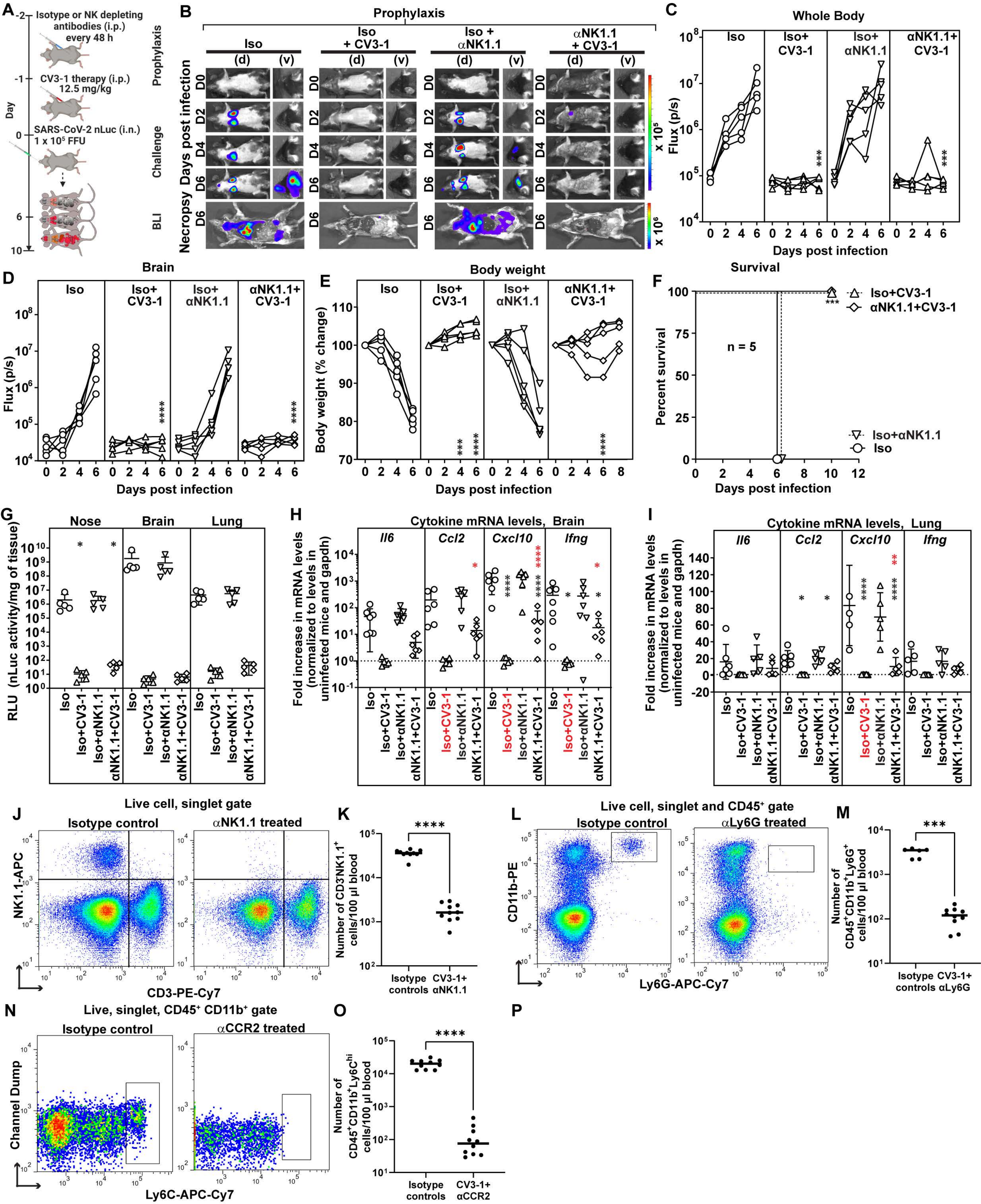

A
Isotype or NK depleting antibodies (i.p.) every 48 h
CV3-1 therapy (i.p.) 12.5 mg/kg
SARS-CoV-2 nLuc (i.n.) 1 x 10^5 FFU
Day
Prophylaxis
Challenge
B
Prophylaxis
Iso
Iso + CV3-1
Iso + αNK1.1
αNK1.1 + CV3-1
Days post infection
Necropsy
BLI
C
Whole Body
Flux (p/s)
Days post infection
D
Brain
E
Body weight
Body weight (% change)
F
Survival
Percent survival
n = 5
Iso+CV3-1
αNK1.1+CV3-1
Iso+αNK1.1
Iso
G
Nose
Brain
Lung
RLU (nLuc activity/mg of tissue)
H
Cytokine mRNA levels, Brain
Fold increase in mRNA levels (normalized to levels in uninfected mice and gapdh)
I
Cytokine mRNA levels, Lung
J
Live cell, singlet gate
Isotype control
αNK1.1 treated
NK1.1-APC
CD3-PE-Cy7
K
Number of CD3⁻NK1.1⁺ cells/100 µl blood
Isotype controls
CV3-1+ αNK1.1
L
Live cell, singlet and CD45⁺ gate
αLy6G treated
CD11b-PE
Ly6G-APC-Cy7
M
Number of CD45⁺CD11b⁺Ly6G⁺ cells/100 µl blood
CV3-1+ αLy6G
N
Live, singlet, CD45⁺ CD11b⁺ gate
αCCR2 treated
Channel Dump
Ly6C-APC-Cy7
O
Number of CD45⁺CD11b⁺Ly6C^hi cells/100 µl blood
CV3-1+ αCCR2
P
